# Dynamic birth and death of Argonaute gene family functional repertoire across *Caenorhabditis* nematodes

**DOI:** 10.1101/2024.10.27.620551

**Authors:** Daniel D. Fusca, Katja R. Kasimatis, Hongyu Vicky Zhu, Asher D. Cutter

## Abstract

Diverse small RNA pathways, comprised of Argonaute effector proteins and their bound small RNA molecules, define critical systems for regulating gene expression in all domains of life. Some small RNA pathways have undergone significant evolutionary change in nematode roundworms, including gains of novel Argonaute genes and losses of entire pathways. Differences in the functional complement of Argonautes among species therefore profoundly influence the available repertoire of mechanisms for gene regulation. Despite intensive study of Argonaute function in *Caenorhabditis elegans*, the extent of Argonaute gene family dynamism and functional breadth remains unknown. We therefore comprehensively surveyed Argonautes across 51 *Caenorhabditis* species, yielding over 1200 genes from 11 subfamilies. We documented multiple cases of diversification, including the birth of a potentially novel Argonaute subfamily and the origin of the ALG-5 microRNA Argonaute near the base of the *Caenorhabditis* phylogeny, as well as evidence of adaptive sequence evolution and gain of a new splice isoform for CSR-1 in a clade of 31 species. We also detected repeated independent losses of multiple components of the piRNA pathway, mirroring other instances of piRNA pathway loss across the phylum. Gene gain and loss occurs significantly faster than expected within several Argonaute subfamilies, potentially associated with transposable element proliferation coevolving with WAGO-9/10/12 copy number variation. Our characterization of Argonaute diversity across *Caenorhabditis* demonstrates exceptional functional dynamism in the evolution of gene regulation, with broad implications for mechanisms of control over ontogenetic development and genome integrity.

**Author Summary:** For organisms to develop properly to survive and reproduce, they must express their genes in the right amount, in the appropriate cell types and time during development. One important mechanism that organisms use to regulate gene expression involves small RNA pathways, where short molecules of RNA serve as targeting guides by binding to Argonaute effector proteins. To understand how small RNA pathways evolve over time, we searched for Argonaute genes throughout the genomes of 51 species of *Caenorhabditis* nematode worms and found over 1200 Argonaute genes belonging to 11 different Argonaute subfamilies. We then documented cases where species have evolved potentially new types of Argonautes, or new protein isoforms of existing Argonautes. We also identified repeated cases of evolutionary loss of entire Argonaute subfamilies, including for the PRG-1 Argonaute needed in the piRNA regulatory pathway, and characterized how some Argonaute subfamilies gain and lose genes significantly faster than expected. Our findings demonstrate substantial variation in the functional repertoire of Argonaute genes found among *Caenorhabditis* species, with this evolutionary dynamism implicating fundamental differences between species in how they regulate gene expression across their genomes throughout development.

## Introduction

The development, survival, and reproduction of all organisms depends crucially on the proper regulation of gene expression. One fundamental system of gene regulation employed by all known domains of life are small RNA pathways, in which short (20 – 30 nucleotides long) RNA molecules serve as sequence-specific guides for Argonaute effector proteins [1, 2].

Together, Argonaute and small RNA form the RNA Induced Silencing Complex (RISC), which is then able to regulate the expression of target sequences based on the sequence of the small RNA guide and the cell types in which the Argonaute and small RNA are expressed. Depending on the type of Argonaute, the effect on target expression can range from repressive (e.g., PRG-1) to licensing (e.g., CSR-1) [3–6] and the regulated sequences can include loci such as protein-coding genes, pseudogenes, transposable elements, and noncoding RNA genes [7]. How distinct types of Argonaute proteins evolve to use different classes of small RNA thus defines the regulatory repertoire available for RISC-mediated genome regulation in a given species.

Small RNA pathways are exceptionally well-characterized in the nematode roundworm *Caenorhabditis elegans* [8–10]. *C. elegans* expresses three different classes of small RNAs: microRNAs and piRNAs, which are genomically encoded, and endogenous small interfering RNAs (endo-siRNAs), which are transcribed by RNA-dependent RNA polymerases (RdRPs) that use the target transcripts as a template for small RNA production. The *C. elegans* genome encodes 20 protein-coding Argonaute genes, though one (WAGO-5) may not have functional activity [7]. These *C. elegans* Argonautes bind to microRNAs (RDE-1, ALG-1, ALG-2, and ALG-5), piRNAs (PRG-1; piRNAs are also known as 21U-RNAs in *Caenorhabditis*), and two classes of endo-siRNAs known as 22G-RNAs and 26G-RNAs, named based on their stereotypical length and first nucleotide. 26G-RNAs are bound by ALG-3, ALG-4, and ERGO-1, whereas 22G-RNAs are bound by a host of a dozen Argonautes (the WAGO Argonautes CSR-1, VSRA-1, WAGO-1, WAGO-3/PPW-2, WAGO-4, WAGO-6/SAGO-2, WAGO-7/PPW-1, WAGO-8/SAGO-1, WAGO-9/HRDE-1, WAGO-10, and WAGO-12/NRDE-3; as well as RDE-1) [7]. RDE-1 additionally binds to small RNAs derived from exogenous double stranded RNA, known as exo-siRNAs [11]. Loss of certain individual Argonautes in *C. elegans* can lead to lethality or sterility [11–13], whereas other individual Argonautes are dispensable for survival and reproduction [7, 14], highlighting both their crucial role in maintaining proper gene regulation and the diversity of potential selective pressures acting on this family of proteins.

Given that changes in gene regulation between species can provide the basis for evolution of adaptive phenotypes [15–17], characterizing the evolution of small RNA pathways may be useful for shedding light on the process of adaptation. Indeed, microRNAs and protein-coding genes involved in small RNA pathways are known to have experienced positive selection in some species [18–21], and differences in small RNA expression between populations may contribute to certain local adaptations in nature [22, 23]. To this end, while studies on nematodes such as *C. elegans* have yielded a great deal of insight into the mechanistic functioning of small RNA pathways, nematodes also represent an interesting study system for understanding how these pathways evolve over time.

The large repertoire of 20 Argonautes in *C. elegans*, compared to just 8 family members in humans, for example [11, 24, 25], implicates the potential for nematodes to show exceptional gene family dynamism with important consequences for divergence in gene regulation. Certain nematode lineages have gained and lost various types of Argonautes and small RNA pathways. For example, while microRNA-binding and piRNA-binding Argonautes are found broadly across animals [25–27], WAGO Argonautes are found exclusively in nematodes and have diversified to control a number of novel different subtypes of small RNAs [11, 24]. In addition, the piRNA pathway that is crucial to transposable element regulation in animals such as *Drosophila* and mice [28, 29] was lost independently in multiple nematode clades, though these clades nevertheless maintain the ability to silence transposable elements via different small RNA pathways [30]. Despite these profound changes to small RNA pathways in the nematode phylum, it is less clear to what extent Argonautes and small RNA pathways evolve at more recent timescales. The *Caenorhabditis* genus is an exceptional study system for addressing such questions, owing to the abundance of Argonautes and small RNA pathways in combination with increasing availability of genome sequencing data for diverse species and populations [31, 32].

Indeed, *Caenorhabditis* species such as *C. inopinata* and *C. plicata* were reported to lack copies of key Argonaute genes that are present in *C. elegans* (ERGO-1 and PRG-1, respectively) [33, 34]. Studies in *C. elegans* also showed that small RNA pathway genes can exhibit substantial variability within-species both in coding sequences and in gene expression [35]. These case studies motivate a comprehensive comparison of Argonautes across *Caenorhabditis* to characterize the full extent to which the Argonaute gene family varies to underlie divergent small RNA regulatory repertoires across species.

To investigate how small RNA regulatory pathways evolve, we comprehensively surveyed Argonaute genes in *Caenorhabditis*. From over 1200 identified Argonaute genes in 51 species, we tested for diversification into novel Argonaute subfamilies and functional forms, losses of entire pathways, and distinctive rates of family size expansion and contraction relative to other gene families encoded in these nematode genomes. This resource reveals previously unappreciated dynamism in Argonaute gene family diversification and function, which will serve to catalyze the creation and testing of new hypotheses about the evolution of small RNA pathways mediated by Argonaute proteins.

## Results

### *Caenorhabditis* genomes encode a large and varied complement of Argonaute genes

To understand the evolution of small RNA pathways in *Caenorhabditis*, we searched for Argonaute genes in the genomes and transcriptomes of 51 *Caenorhabditis* species, ranging from the closest relatives of *C. elegans* to some of the earliest-diverging members of the genus. Based on a combination of orthology to known Argonautes in *C. elegans* and interrogation of sequences for the presence of functional domains that are highly specific to Argonautes (Fig. 1A), and after filtering out spurious candidate gene copies (see Materials & Methods), we identified a total of 1213 Argonaute genes in these 51 species. These Argonautes predominantly clustered into 11 distinct subfamilies of orthologous genes, labeled with reference to the *C. elegans* gene(s) contained in each subfamily (Fig. 1B), and which varied substantially in protein sequence conservation between species. WAGO families typically showed the lowest sequence conservation (Fig. 1C, Fig. S1); for example, orthologs of the WAGO VSRA-1 exhibited only 30.0% average protein sequence identity to each other compared to 87.3% average identity among ALG-3/4 orthologs.

**Figure 1.**
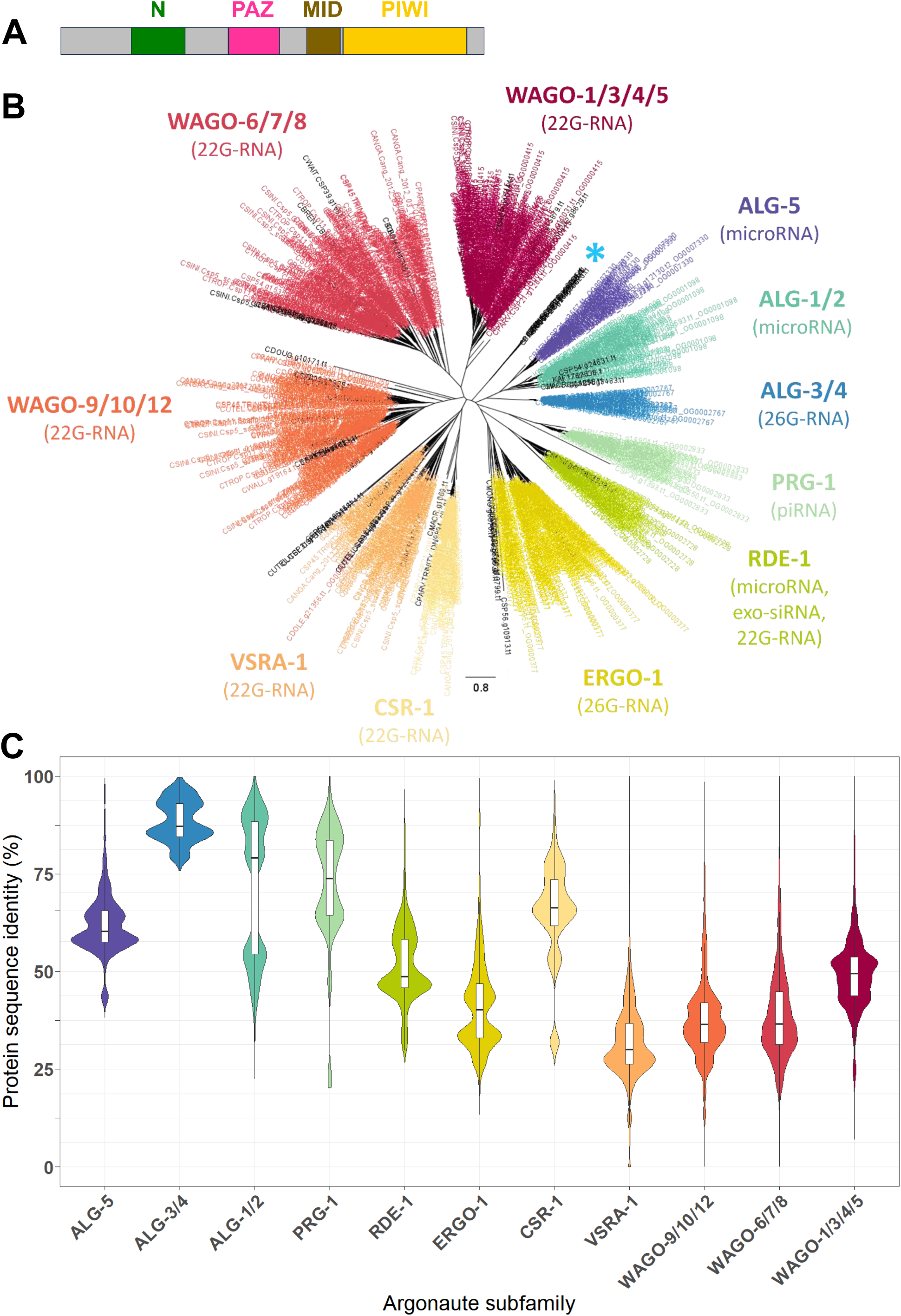
Identification of 1213 *Caenorhabditis* Argonautes in 11 distinct subfamilies. (A) Stereotypical domain structure (N-terminal, PAZ, MID, and PIWI domains) of a eukaryotic Argonaute protein. Domain locations are based off of their positions in ALG-1 in *C. elegans*. (B) Maximum-likelihood gene tree of 1213 identified Argonaute protein sequences across 51 species of *Caenorhabditis*. Colors correspond to the 11 Argonaute orthogroups identified by OrthoFinder. Gene names in black represent genes that were not grouped with Argonautes by OrthoFinder, but nevertheless contain Argonaute functional domains. “*” indicates a group of 7 divergent Argonautes found in *C. panamensis*. (C) Violin plots showing the distributions of pairwise protein sequence identities, measured between all pairs of different Argonautes belonging to the same subfamily, for each of the 11 Argonaute subfamilies.

Argonaute proteins are organized with a stereotypical domain structure in eukaryotes (N-PAZ-MID-PIWI domains) (Fig. 1A) to carry out functions such as small RNA binding and target cleavage [1]. As expected, nearly all 1213 identified Argonaute proteins (97.6%) contained a PIWI domain that could be identified by the domain-annotation software InterProScan when using the Pfam domain database. However, we detected PAZ, MID, and N domains in only 63.7%, 14.8%, and 18.3% of Argonautes, respectively, with these domain annotations being undetected even in some *C. elegans* Argonaute proteins (e.g., ALG-1/2/3/4/5 were the only *C. elegans* Argonautes with MID domains annotated by InterProScan). To determine whether these apparent domain absences were genuine, we aligned the three-dimensional protein structure of *C. elegans* ALG-1 to the 19 other *C. elegans* Argonaute protein structures, as InterProScan was able to identify all four domain types in ALG-1. We found that all 20 Argonautes in *C. elegans* contained regions that structurally aligned to the N, PAZ, MID, and PIWI domains of ALG-1 (Fig. S2). This observation suggests that all four domain types are likely present in almost all Argonautes across *Caenorhabditis*, despite limitations of sequence-based domain recognition to universally identify them.

We observed substantial Argonaute copy number variation both between *Caenorhabditis* species and between Argonaute families (Fig. 2, Table S1), even after adjusting the gene copy counts to correct for instances of spurious annotation due to gene fragmentation, fusion, and allelism (see Materials & Methods). Species encoded a minimum of 9 Argonaute gene copies (*C. astrocarya*) to a maximum of 46 Argonautes (*C. brenneri*), with a median of 23 Argonaute genes per species. The 11 Argonaute subfamilies differed substantially in how variable their copy numbers were among species (Fig. 2, Table 1). For example, ALG-3/4 and ALG-5 were found in exactly a single copy in almost every species, whereas ERGO-1 was completely absent in 7 species but present with 5 or more copies in 10 species (Fig. 2). Given that ALG-5 was present in every species except for *C. monodelphis*, the most basal species included here, it is possible that this absence indicates that ALG-5 was gained in the other species rather than lost in *C. monodelphis*. Consistent with this possibility, we were unable to identify ALG-5 in the annotated protein sequences of 5 non-*Caenorhabditis* nematode species (*Diploscapter coronatus, Diploscapter pachys*, *Haemonchus contortus*, *Oscheius tipulae*, and *Pristionchus pacificus*), with *D. coronatus* and *D. pachys* being the most closely related nematodes with sequenced genomes to *Caenorhabditis* [36]. However, an alternative explanation is that ALG-5 was gained in the ancestor of all *Caenorhabditis* species but then subsequently lost in *C. monodelphis*. In support of this, we identified a potential ALG-5 ortholog in *Caenorhabditis krikudae*, a species not included in our Argonaute survey as its transcriptome was sequenced after we began our survey [37]. Phylogenies have placed *C. krikudae* in a clade with *C. monodelphis* at the very base of the genus [37], so assuming this placement is correct, the birth of ALG-5 likely occurred near the common ancestor of the *Caenorhabditis* genus.

**Figure 2.**
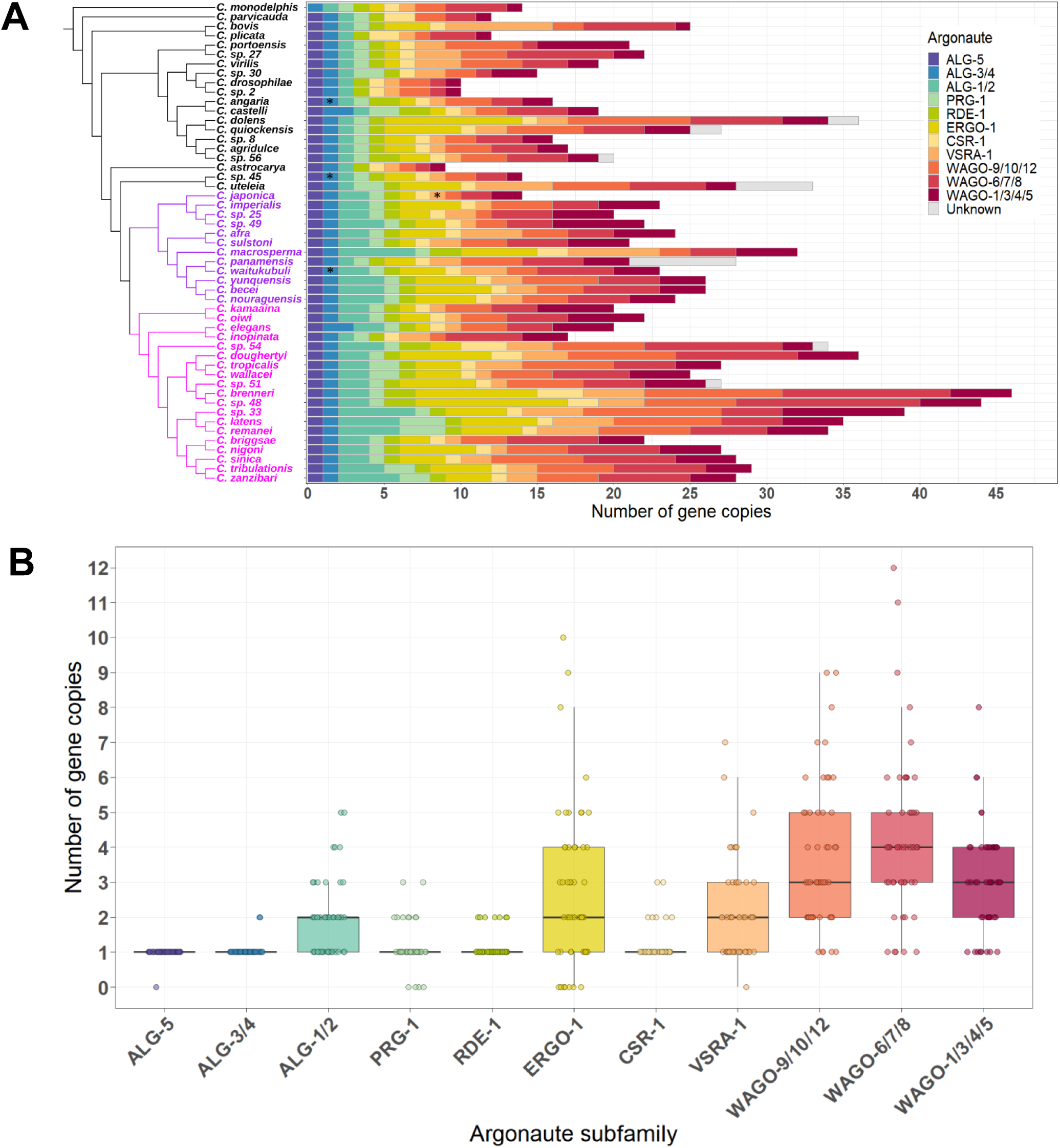
Argonaute complement varies between *Caenorhabditis* species. (A) Number of Argonaute genes in all 51 species belonging to each of the 11 Argonaute subfamilies. “Unknown” indicates genes with Argonaute domains that could not be confidently assigned to any known Argonaute subfamily. “*” indicates cases where a subfamily could only be detected in a species as one or more short gene fragments. Purple and magenta on the species tree represent the Japonica group and the Elegans group respectively, which collectively form the Elegans supergroup. (B) Boxplots showing the number of Argonaute genes identified in each individual species, separated by Argonaute subfamily.

**Table 1:**
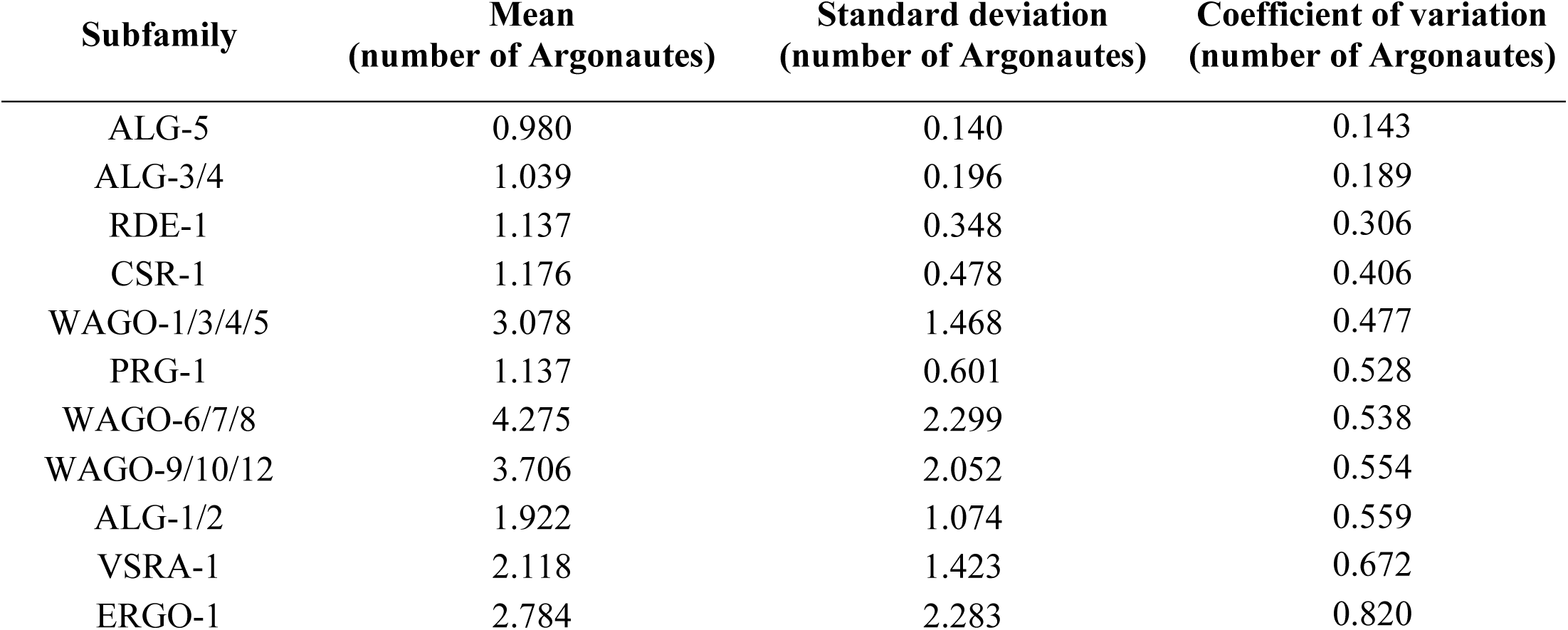
Mean, standard deviation, and coefficient of variation for the number of Argonaute genes found in each species, separately for all 11 Argonaute subfamilies.

| Subfamily | Mean<br>(number of Argonautes) | Standard deviation<br>(number of Argonautes) | Coefficient of variation<br>(number of Argonautes) |
| --- | --- | --- | --- |
| ALG-5 | 0.980 | 0.140 | 0.143 |
| ALG-3/4 | 1.039 | 0.196 | 0.189 |
| RDE-1 | 1.137 | 0.348 | 0.306 |
| CSR-1 | 1.176 | 0.478 | 0.406 |
| WAGO-1/3/4/5 | 3.078 | 1.468 | 0.477 |
| PRG-1 | 1.137 | 0.601 | 0.528 |
| WAGO-6/7/8 | 4.275 | 2.299 | 0.538 |
| WAGO-9/10/12 | 3.706 | 2.052 | 0.554 |
| ALG-1/2 | 1.922 | 1.074 | 0.559 |
| VSRA-1 | 2.118 | 1.423 | 0.672 |
| ERGO-1 | 2.784 | 2.283 | 0.820 |

The three species with 10 or fewer total Argonautes (*C. astrocarya*, *C. drosophilae*, and *C.* sp. 2) likely reflect genuine gene family contractions rather than genome assembly errors, given that they showed high BUSCO completeness scores for their genome assemblies (94.6%, 93.5%, and 94.4% of BUSCO genes were complete and single-copy for the respective species) that were similar to other species with many Argonaute copies (e.g., *C.* sp. 48 has 44 Argonaute genes and 95.6% of BUSCO genes complete and single-copy) (Table S2). The overall number of Argonaute genes in a species correlated positively with genome size (*P* = 0.0034, PGLS regression), but not with N50 of the genome assembly (*P* = 0.33), likely reflecting the general positive associations of protein-coding gene complement with both genome size (*P =* 1.6x10^-6^) and number of Argonaute genes (*P* = 0.0010) rather than some aspect of assembly quality (Fig. S3). The functional complement of Argonaute genes encoded by a given species has thus experienced a large amount of evolution across *Caenorhabditis*.

### Argonaute gene family size is dynamic across the phylogeny

Given that some Argonaute subfamilies such as ERGO-1 appear to evolve rapidly in terms of copy number variation across species, we aimed to quantify and compare Argonaute gene family dynamics relative to all of the gene families in *Caenorhabditis*. We employed CAFÉ [38] to test a genome-wide set of 11,442 gene families for evidence of significant expansions or contractions at any point in the *Caenorhabditis* phylogeny, compared to the expected rate of gene birth and death. We detected significant gene family size changes in 430 of these gene families, which included 3 of the 11 Argonaute subfamilies (ERGO-1, WAGO-6/7/8, and WAGO-9/10/12; this result was robust to the exclusion of the five species with the lowest-quality genome assemblies based on BUSCO scores). *Caenorhabditis* Argonautes were thus 7- fold enriched among these gene families with evidence of significant gene family size changes (27.3% of Argonaute families had significant size changes compared to 3.8% of all gene families tested; *P* = 0.007, Fisher’s exact test). After stringent multiple-testing correction (FDR ≤ 0.05) reducing the genome-wide total to 274 significant gene families, ERGO-1 was the only Argonaute subfamily that remained significant. In *C. elegans*, ERGO-1 mediates regulation of its targets by binding to 26G-RNAs, leading to the downstream production of certain WAGO-associated 22G-RNAs [7, 39]. We conclude that ERGO-1 (and potentially also WAGO-6/7/8 and WAGO-9/10/12) underwent significant changes in gene family size in *Caenorhabditis* (Fig. S4) that are not attributable to the background rate of gene birth and death, or to artifacts of genome assembly and gene annotation.

Having established that copy number evolves rapidly for several Argonaute subfamilies within *Caenorhabditis*, we next aimed to explore potential evolutionary forces that could drive gene gain and loss dynamism. Because some Argonautes target transposable elements for silencing [7], we hypothesized that genomic repeat content might correlate with the number of encoded Argonaute genes. To test this hypothesis, we quantified the proportion of the genome consisting of repetitive elements for each of 50 *Caenorhabditis* genome assemblies (*C.* sp. 45 was excluded as only a transcriptome assembly was available) and generated a phylogenetically- controlled statistical model of repeat content as a linear function of Argonaute gene counts, genome size, and total number of non-Argonaute protein-coding genes. We found that repeat content did not associate significantly with the total number of Argonaute genes (*P* = 0.21, PGLS regression). However, when incorporating each of the 11 Argonaute gene subfamilies as separate model terms, we observed repeat content to negatively associate with the number of orthologs of WAGO-9/10/12 (*P* = 0.0073) and CSR-1 (*P* = 0.0037), after accounting for variation in genome size and number of non-Argonaute genes. These findings for WAGO-9/10/12 are robust to exclusion of the five species with the lowest-quality gene annotations (WAGO-9/10/12 *P* = 0.049), but a positive relationship of repeat content with RDE-1 replaces the association with CSR-1 (RDE-1 *P* = 0.020, albeit with range of just 1 to 2 copies of RDE-1). Focusing our modeling on individual classes of repeats, we found that WAGO-9/10/12 ortholog number was significantly negatively associated with both retrotransposon content (*P* = 0.024) and DNA transposon content (*P* = 0.0011) but not rolling-circle transposon content (*P* = 0.20), with these relationships again persisting when the five species with the lowest-quality gene annotations were excluded. No other Argonaute subfamily showed a significant relationship with any of these transposon subtypes, though on average only 20% of the repeat content in each species could be classified to a known subtype. In *C. elegans*, Argonautes in the WAGO-9/10/12 subfamily are known to target transposable elements and work to prevent their proliferation [7, 40, 41]. Consequently, our findings are consistent with selection driving higher copy numbers of WAGO-9/10/12 subfamily members to control repetitive element activity in genomes (Fig. S5).

### Microevolution of Argonautes within species recapitulates macroevolutionary patterns

Based on our observation at the macroevolutionary scale that the rate at which species gain and lose gene copies is much higher for some Argonaute families (e.g., ERGO-1) than others (e.g., ALG-5) (Table 1), we sought to determine whether these trends also hold at the more recent evolutionary timescale of microevolution within species. To do so, we extracted data on within-species genetic variation for Argonaute genes in the genomes of 550 isotypes of *C. elegans* and 641 isotypes of *C. briggsae* from the *Caenorhabditis* Natural Diversity Resource (CaeNDR) [32]. In both species, we observed that mutations that likely create pseudogene alleles (e.g., frameshift indels that introduce premature stop codons) tend to occur with higher incidence in those Argonaute families that show high rates of gene gain and loss between species (Fig. 3A, B; Fig. S6A, B). This positive relationship between interspecies gene copy number variation and the number of isotypes carrying pseudogene alleles was statistically significant for *C. elegans* (*P* = 0.041, Spearman’s rank correlation) but not *C. briggsae* (*P* = 0.32). This finding suggests that the relative rates of Argonaute subfamily evolution are similar at both macroevolutionary and microevolutionary timescales.

**Figure 3.**
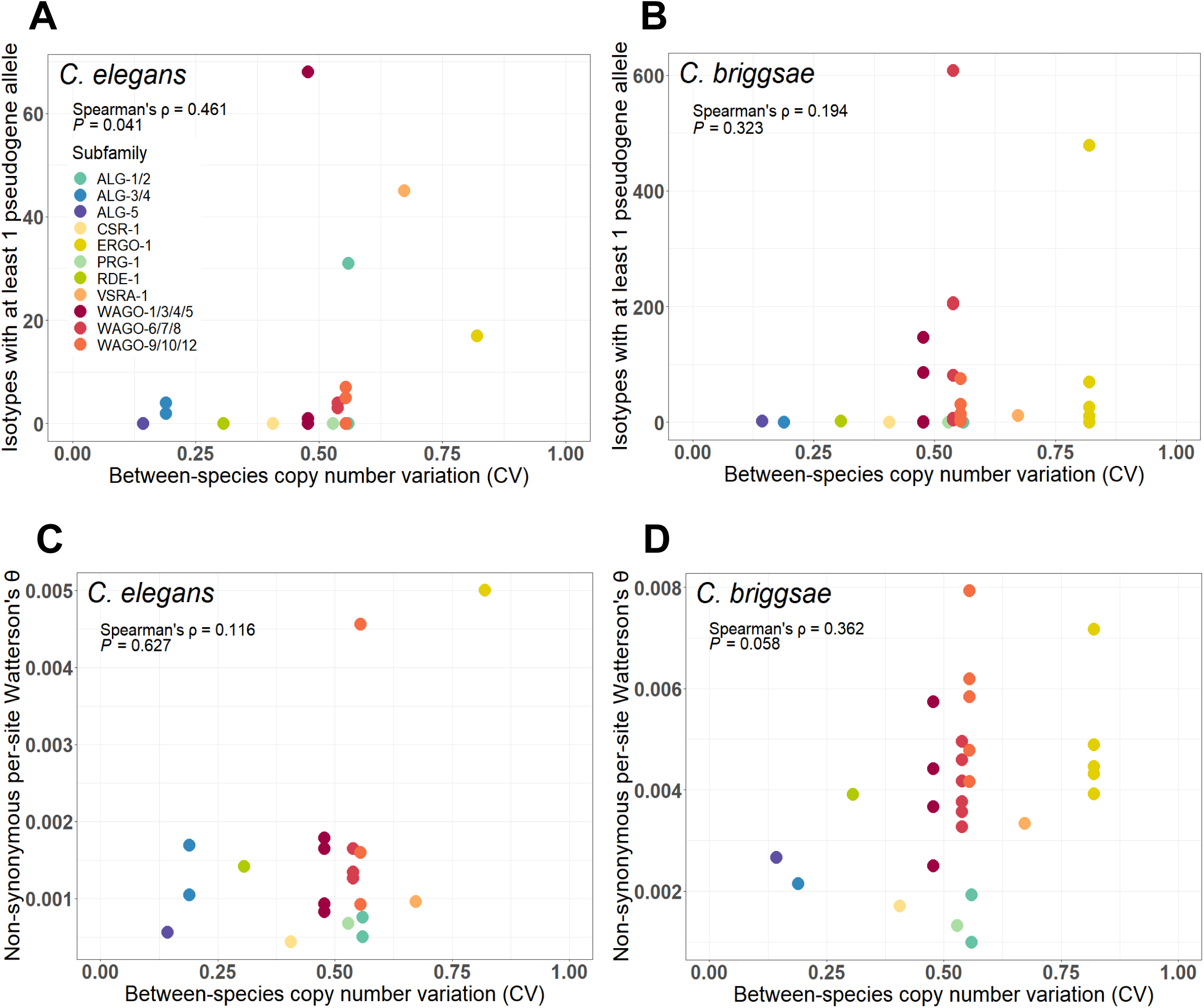
Argonaute gene microevolution in *C. elegans* and *C. briggsae*. For every Argonaute gene in *C. elegans* (A, C) and *C. briggsae* (B, D), comparisons of the between-species copy number variation for that Argonaute subfamily (measured by the coefficient of variation), and either the number of isotypes carrying at least 1 pseudogene allele for that Argonaute gene (A, B), or the per-site Watterson’s θ at non-synonymous sites for that Argonaute gene (C, D).

As an example, we identified at most two isotypes with potentially-pseudogenizing mutational variants of CSR-1 and ALG-5 in *C. elegans* and *C. briggsae*, consistent with the extreme conservation of exactly one single copy of these genes in most species. In contrast, VSRA-1 and ERGO-1 are present as a single copy in fewer than half of the 51 species. We identified pseudogene variants of VSRA-1 in 45 *C. elegans* isotypes and 12 *C. briggsae* isotypes, and found that most ERGO-1 gene copies in these species harbor pseudogene alleles in 10 or more different isotypes. Metrics of within-species nucleotide diversity at non-synonymous sites, measured by Watterson’s θ as a proxy for the strength of purifying selection, also tended to be higher for Argonaute subfamilies with more rapid between-species gains and losses (e.g., ERGO-1) compared to Argonaute subfamily members that usually are encoded with a single-copy (e.g., CSR-1) (Fig. 3C, D; Fig. S6C, D). However, this positive trend between non-synonymous sequence diversity and interspecies gene copy number variation was not significant for *C. elegans* (*P* = 0.63, Spearman’s rank correlation) and marginally non-significant for *C. briggsae* (*P* = 0.058). Our results indicate that Argonaute genes with higher rates of duplications and deletions between species may also be generally less constrained by purifying selection to maintain conserved coding sequences within species.

### Ablation and diversification of Argonaute function across the phylogeny

Our analyses revealed several cases where a species appeared to be missing an Argonaute subfamily entirely that could be indicative of the ablation of some small RNA regulatory functionality. For example, we observed that *C*. *plicata* did not appear to encode any copies of the piRNA Argonaute PRG-1, as previously described [34]. We leveraged publicly-available RNA-seq datasets to confirm absence of expression for cases of putative Argonaute loss where possible. We were unable to detect expression of PRG-1 in *C. plicata*, ALG-5 in *C. monodelphis*, VSRA-1 in *C. parvicauda*, and ERGO-1 in *C. inopinata*, *C. plicata*, and *C. parvicauda*, but recovered expression of CSR-1 in *C. parvicauda* despite it not having been detected in the genome assembly (aside from a highly-diverged duplicate). Although some species with putatively missing Argonautes lack available RNA-seq datasets, this transcriptomic approach provided additional evidence for these cases of Argonaute gene subfamily absence.

In addition to losses of entire Argonaute subfamilies in certain species, we also observed several cases of Argonaute diversification. Several species appear to encode highly-diverged duplicates in addition to their more conserved copy of an otherwise strongly conserved Argonaute gene, as for the case of PRG-1 in *C.* sp. 30 and CSR-1 in *C. macrosperma* and *C. parvicauda*. Using PCR amplification and Sanger sequencing, we could verify the sequence of 7 of the 11 exons of the *C.* sp. 30 PRG-1 duplicate, though we were unable to verify the sequence of the putatively divergent copy of CSR-1 in *C. macrosperma*. Divergent duplicate Argonaute gene copies provide candidates for possible instances of neofunctionalization, in which a duplicated gene goes on to diverge in function from its parent gene.

The most notable instance of diversification that we identified occurred in *C. panamensis*: 7 Argonaute genes clustered with each other, and separate from all other Argonautes, on our gene tree (CSP28.g16020.t1, CSP28.g16021.t1, CSP28.g16024.t1, CSP28.g16025.t1, CSP28.g16026.t1, CSP28.g16027.t1, and CSP28.g8935.t1) (Fig. 1B).

Interestingly, 6 of these 7 *C. panamensis* Argonautes (all aside from CSP28.g8935.t1) also clustered together physically in the genome as a series of tandem duplicates on scaffold CSP28.scaffold3_cov87 (Fig. 4A). Comparisons of these 7 putatively novel Argonautes to *C. elegans* revealed greatest protein sequence similarity to ALG-1 and ALG-2, but absolute similarity scores were low (24.3% - 28.5% identical in sequence) (Fig. 4B). After performing PCR and Sanger sequencing, we were able to verify the accuracy of the annotated coding sequences for the 6 tandemly-duplicated Argonautes (aside from the first 5 annotated exons of CSP28.g16021.t1, which appears to be an adjacent but separate gene). These results rule out genome assembly artifacts as a cause of the high sequence divergence for 6 of the putatively novel Argonaute genes in *C. panamensis*.

**Figure 4.**
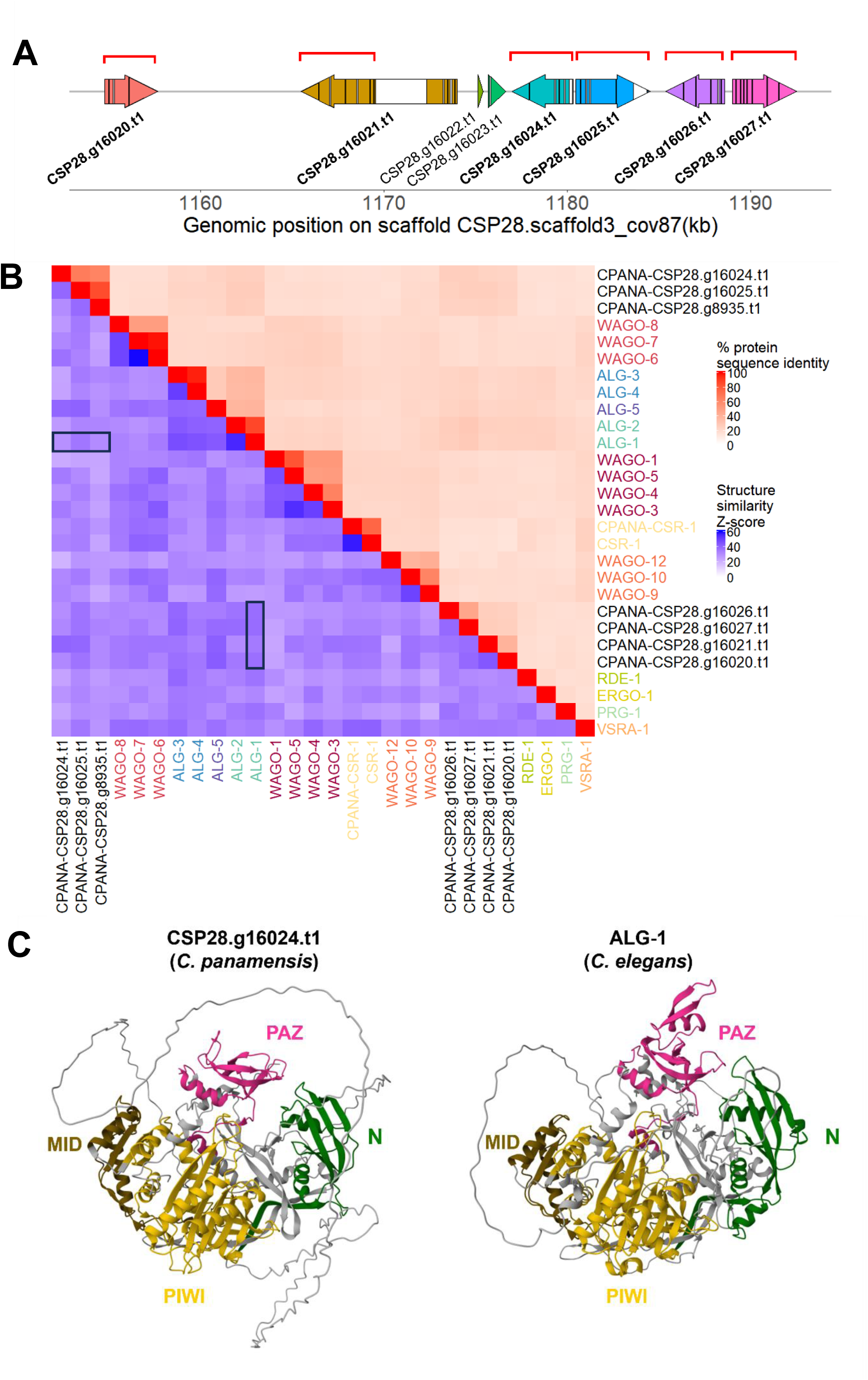
*C. panamensis* contains a cluster of divergent Argonaute genes. (A) Positions of genes in the cluster of *C. panamensis* divergent Argonautes on scaffold CSP28.scaffold3_cov87. Bolded gene names denote the 6 Argonautes. Colored blocks represent exons, and intervening white blocks represent introns. Red bars above genes indicate coding sequences that we were able to verify with Sanger sequencing. (B) Heatmap of pairwise protein sequence identity (above and along the diagonal, in red) and structural similarity (below the diagonal, in blue) between all 20 *C. elegans* Argonautes, the 7 *C. panamensis* divergent Argonautes, and *C. panamensis* CSR-1 as a control. Higher structure similarity Z-scores indicate more similar protein structures. “CPANA” denotes proteins from *C. panamensis*; all other proteins are from *C. elegans*. Black outlined boxes indicate structural comparisons between *C. elegans* ALG-1 and the 7 *C. panamensis* divergent Argonautes, as shown in panel C. (C) ColabFold-predicted protein structure of CSP28.g16024.t1, a divergent Argonaute from *C. panamensis* (left). For comparison, the structure of *C. elegans* ALG-1 from the AlphaFold Protein Structure Database is also shown (right). Colors indicate the typical Argonaute domains, with domains in CSP28.g16024.t1 identified based on structural alignments to ALG-1. Note that while the PAZ domain is mostly present in CSP28.g16024.t1, it is reduced in size compared to ALG-1.

To better understand the potential function of these 7 putatively novel Argonautes, we predicted their protein structures using ColabFold’s implementation of AlphaFold2 [42, 43], and compared the resulting structures to the existing AlphaFold2 protein structures of all 20 *C. elegans* Argonautes (Fig. 4B, C). As we observed for protein sequences, the 3D-structures of these 7 divergent *C. panamensis* Argonautes were most similar to *C. elegans* ALG subfamily proteins (ALG-2, ALG-3, and ALG-5), though the 5 *C. elegans* ALG structures were almost always more similar to each other than they were to these 7 *C. panamensis* Argonautes (Fig. 4B). This reduced structural similarity is unlikely to be due to general problems with ColabFold, as our ColabFold-predicted structure of *C. panamensis* CSR-1 was highly similar to the structure of CSR-1 in *C. elegans* (Fig. 4B). Thus, *C. panamensis* appears to encode at least 6 duplicates of an ALG-like Argonaute that diverged substantially in both sequence and structure from the known ALG families. Surprisingly, while all of these putatively novel Argonautes have predicted PIWI domains, structural alignments to *C. elegans* ALG-1 revealed that four of them appear to be mostly or completely missing structures corresponding to the PAZ domain that typically functions to anchor the 3′ end of a small RNA (Fig. S7), reminiscent of some prokaryotic “short” Argonautes [1]. Even when present, the PAZ domain was sometimes noticeably reduced in size (Fig. 4C). It is therefore possible that at least some of the distinctive and putatively novel Argonaute genes from *C. panamensis* may function in non-canonical ways or may not act to mediate small RNA regulation.

### Evidence for repeated independent losses of the piRNA pathway in *Caenorhabditis*

In addition to confirming loss of PRG-1 from the *C. plicata* genome [34], we were unable to identify PRG-1 in the genomes of *C. drosophilae*, *C.* sp. 2, and *C. astrocarya*. Reminiscent of repeated losses of PRG-1 across the nematode phylum [30], our analysis implies at least 3 independent losses of PRG-1 within the *Caenorhabditis* genus (*C. drosophilae* and *C.* sp. 2 are sister species). Supporting our conclusion that PRG-1 has been lost in these species, we were unable to PCR amplify exons 4-8 of PRG-1 in *C. plicata* or *C. drosophilae* despite successful amplification in several diverse control species (*C.* sp. 30, *C. virilis, C. macrosperma*, *C. panamensis*, and *C. elegans*).

*C. elegans* piRNAs primarily bind to PRG-1, and *prg-1* mutants have reduced levels of piRNAs [12, 44, 45]. Therefore, we hypothesized that the four *Caenorhabditis* species lacking a copy of PRG-1 do not have functional piRNA silencing pathways. To further test this hypothesis, we selected 7 genes (*prde-1*, *snpc-4*, *tofu-4*, *tofu-5*, *pid-1*, *henn-1*, and *parn-1*) that are involved in piRNA transcription and processing [10] and assessed their presence across *Caenorhabditis*. Among these 7 genes, PARN-1 was missing in all 4 species that lack PRG-1 (Fig. 5A). Aside from these four species, PARN-1 only appeared to be missing in one other species (*C. parvicauda*), meaning that losses of PARN-1 associate primarily with species that do not appear to have a piRNA-binding Argonaute. In *C. elegans*, PARN-1 is an exonuclease that trims the 3′ ends of piRNA precursors, and is required for proper piRNA-mediated silencing [46]. We have therefore identified four *Caenorhabditis* species that have seemingly lost multiple core components of the piRNA pathway.

**Figure 5.**
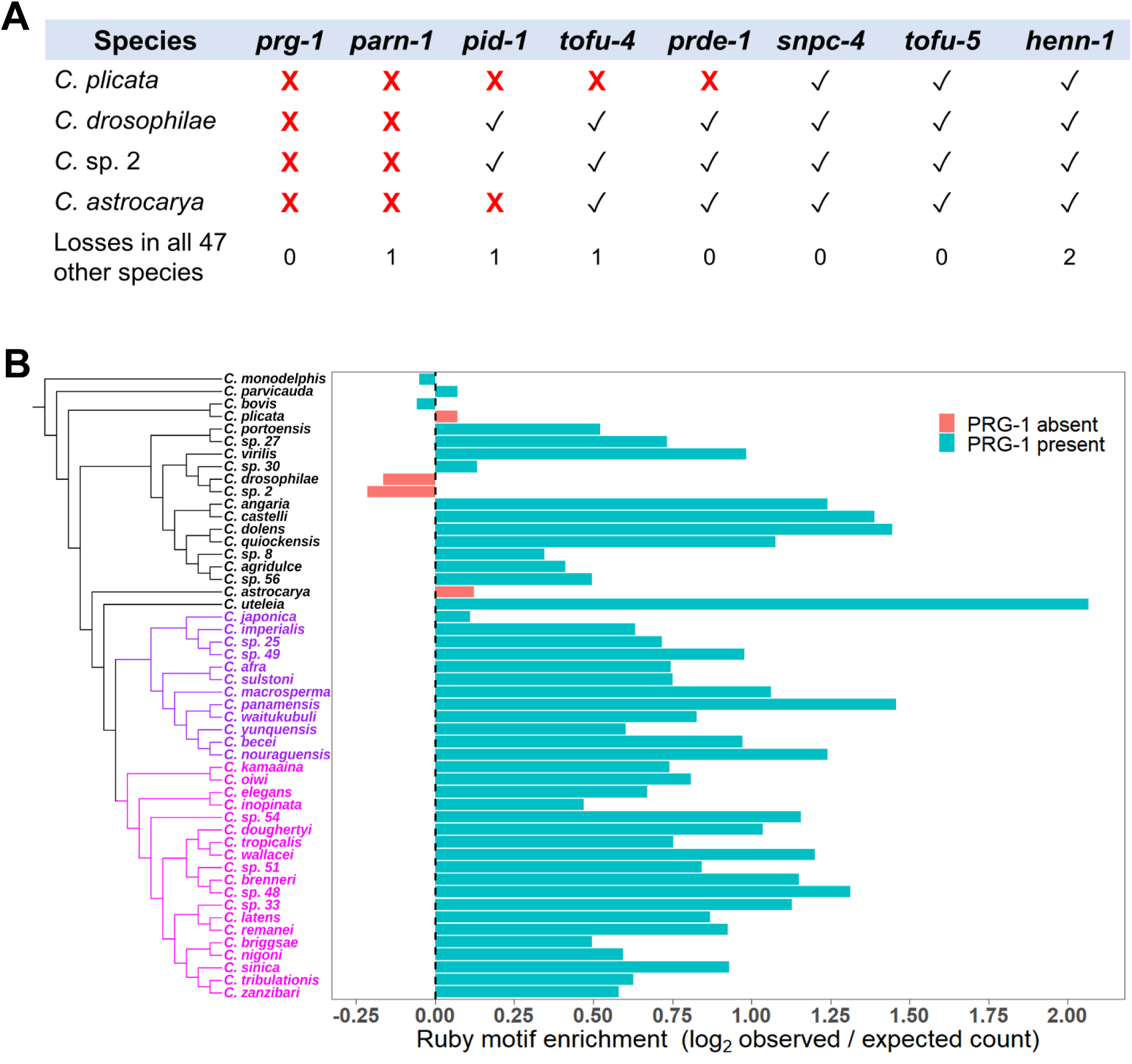
Multiple independent losses of piRNA pathway components in *Caenorhabditis*. (A) Presence of piRNA pathway components in *C. plicata*, *C. drosophilae*, *C.* sp. 2, and *C. astrocarya*. Checkmarks indicate that a gene is present in at least 1 copy in that species, and “X”s indicate that no copies of the gene were detected. The number of other *Caenorhabditis* species that are missing each gene is also indicated. (B) The number of Ruby motifs observed in the noncoding portion of each species’ genome, normalized against the number of Ruby motifs that would be expected to occur by chance. Dashed vertical line represents Ruby motifs being observed exactly as often as would be expected by chance. Purple and magenta on the species tree represent the Japonica group and the Elegans group respectively, which collectively form the Elegans supergroup.

To further assess the integrity of the piRNA pathway in these four species, we quantified the incidence of the DNA motif known as the Ruby motif. The Ruby motif occurs upstream of most piRNA loci in multiple clades of nematode species, and is conserved across *Caenorhabditis* as the 8-nucleotide sequence CNGTTTCA [34, 47]. This motif is bound by a protein complex that promotes piRNA transcription [48], meaning that *Caenorhabditis* species lacking an active piRNA pathway may also show degradation of Ruby motifs in their genomes. We therefore scanned the noncoding portions of *Caenorhabditis* genomes for instances of the Ruby motif, normalizing the observed motif counts in each species by the number expected from a random DNA sequence of the same total length and dinucleotide composition. Detected Ruby motifs in

*C. elegans* and *C. briggsae* were highly enriched in the known piRNA clusters on chromosomes I and IV [47, 49] (Fig. S8), supporting the validity of our approach. We found that all four species that lack PRG-1 were among the set of species with the lowest incidence of the Ruby motif, relative to the random expectation (Fig. 5B). For example, the Ruby motif was found in *C. elegans* (PRG-1 present) 59% more often than expected by chance, whereas in *C. plicata* (PRG-1 absent), this motif was found only 5% more often than expected by chance. *C. drosophilae* and *C.* sp. 2, which also lack PRG-1, had the lowest Ruby motif content out of all our analyzed species, with Ruby motifs occurring in these genomes 11 – 14% less often than would be expected by chance. Several species that encode PRG-1 also showed reduced Ruby motif incidence in a manner suggesting that piRNA gene expression in basal lineages might utilize a distinct regulatory motif, or potentially exposing an error in the topology of the species tree with respect to the sister clade to *C. bovis – C. plicata*. The species *C. bovis* (sister to *C. plicata*), *C. monodelphis* and *C. parvicauda* (basal outgroups from the *C. plicata – C. bovis* lineage), and *C*. sp. 30 (sister to the *C. drosophilae* – *C*. sp. 2 lineage) all showed similar rarity of Ruby motifs to the species lacking PRG-1. Collectively, the fact that the four species lacking PRG-1 also are missing PARN-1 and have reduced genome-wide incidence of Ruby motifs, and previous analysis of *C. plicata* [34], strongly implicates the repeated loss of the piRNA-pathway mechanism for gene regulation in *C. plicata*, *C. astrocarya*, *C. drosophilae* and *C*. sp. 2.

### Diversification of CSR-1 via gain of alternative splice forms in the Elegans supergroup

*C. elegans* expresses two isoforms of the CSR-1 Argonaute (long CSR-1a and short CSR- 1b). These two isoforms differ exclusively due to an N-terminal exon included only in CSR-1a that encodes a structurally disordered sequence of 163 amino acids rich with arginine-glycine repeats (RG and RGG) [50, 51] (Fig. 6A). The CSR-1a and CSR-1b isoforms regulate the expression of distinct sets of target genes in different tissues, with the longer CSR-1a being expressed in the germline during spermatogenesis and some somatic tissues (e.g., intestine) and the shorter CSR-1b being expressed constitutively in the germline for both oogenesis and spermatogenesis [50]. Previous studies have found that CSR-1 in several other *Caenorhabditis* species contains the exon exclusive to CSR-1a [51], but whether both isoforms occur in these or other *Caenorhabditis* species has not been fully determined.

**Figure 6.**
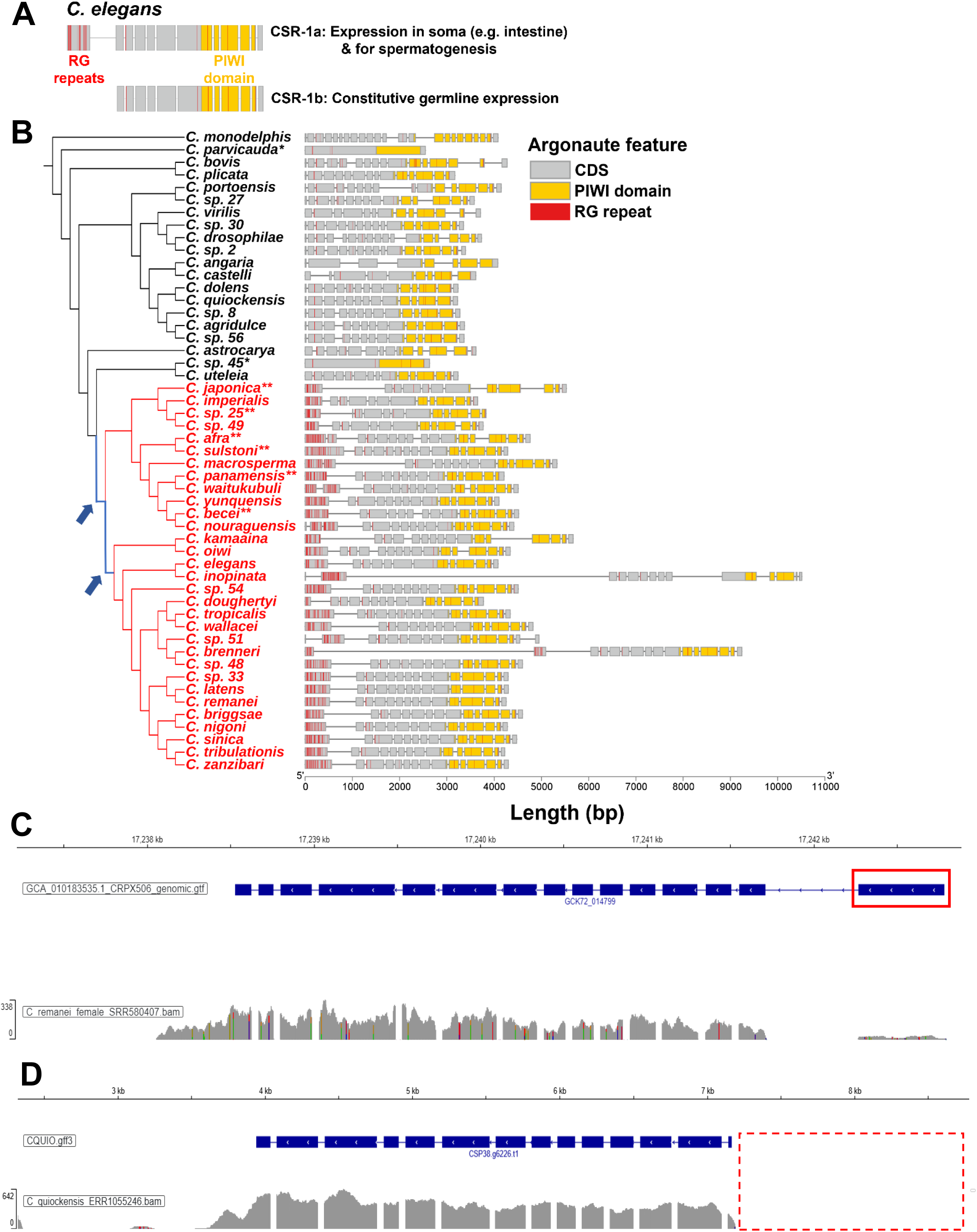
Gain of a novel CSR-1a exon in the Elegans supergroup of species. (A) Schematic of the CSR-1a and CSR-1b isoforms in *C. elegans*. (B) Exon-intron structure of a single representative CSR-1 copy from each *Caenorhabditis* species, highlighting all arginine-glycine (RG) repeats. Species belonging to the Elegans supergroup are shown in red. Blue branches with arrows on the phylogeny are those for which we detected significant positive selection on CSR-1. “*” indicates species where the locations of introns are unknown, as we could only confidently identify CSR-1 in the transcriptomes of these species. “**” indicates species where the CSR-1 gene model was manually updated to include the likely boundaries of the CSR-1a exon. UTRs are not shown due to only being annotated in 4 species. (C) Genome browser track showing the number of aligned RNA-seq reads (grey histogram) along the CSR-1 locus in *C. remanei* (GCK72_014799). Red box indicates the putative CSR-1a exon. (D) Genome browser track showing the number of aligned RNA-seq reads (grey histogram) along the CSR-1 locus in *C. quiockensis* (CSP38.g6226.t1). The upstream region outlined in red is the expected location of a theoretical unannotated CSR-1a exon, if such an exon existed.

To resolve this issue, we first interrogated our set of annotated CSR-1 orthologs to identify species whose CSR-1 gene begins with an RG-rich first exon, as a proxy for species expressing separate CSR-1a and CSR-1b isoforms. We found that all 31 species that we examined within the Elegans supergroup clade contained at least one CSR-1 gene with an RG-rich exon at the 5′ end (sometimes split into two adjacent exons, or preceded by very short exons that are likely annotation errors), consistent with these species expressing a CSR-1a isoform (Fig. 6B). Six species in the Elegans supergroup (*C. japonica*, *C.* sp. 25, *C. afra*, *C. sulstoni*, *C. panamensis*, and *C. becei*) did not have this CSR-1a exon annotated, but manual inspection of these sequences revealed that these absences were due to gene annotation errors (e.g., the CSR- 1a exon in *C. japonica* is mistakenly annotated as a UTR; the CSR-1a exon in *C. panamensis* is unannotated but found in the upstream “noncoding” region). To confirm that CSR-1a is actually expressed as a distinct isoform, we examined available RNA-seq data from a variety of species in the Elegans group (*C. remanei*, *C. tribulationis,* and *C. inopinata*) and Japonica group (*C. macrosperma, C. afra, C. nouraguensis,* and *C. waitukubuli*) within the Elegans supergroup. The putative CSR-1a exon showed reduced expression read depth relative to the rest of the gene for each species except *C. inopinata* (Fig. 6C, Fig. S9), supporting the idea that they express both a CSR-1a and a CSR-1b isoform.

Surprisingly, we were unable to identify a potential CSR-1a exon in any species outside of the Elegans supergroup (Fig. 6B), suggesting that these species may only express a single CSR-1 isoform that resembles CSR-1b. Given that the CSR-1a exon was present but not annotated in several species within the Elegans supergroup, we searched the region upstream of CSR-1 for an unannotated RG-rich exon in a sample of species outside of the Elegans supergroup, but we were unable to find evidence for any such exons. Furthermore, we aimed to verify these absences using RNA-seq data from six species outside of the Elegans supergroup (*C. castelli*, *C. monodelphis*, *C. plicata*, *C. quiockensis*, *C. agridulce*, and *C. uteleia*). However, we found no evidence of an unannotated 5′ exon being expressed in the 5 of 6 species with detectable expression of CSR-1 (Fig 6D, Fig. S10). Again, these data support the absence of a long CSR-1a isoform outside of the Elegans supergroup (though in theory it is possible that RNA-seq data from these species was primarily collected from a sex or developmental stage that may not express the CSR-1a isoform). The most parsimonious explanation for these observations is that the common ancestor of the *Caenorhabditis* genus expressed only a single isoform of CSR-1 resembling CSR-1b, with the origin of a separate CSR-1a isoform containing a novel exon as an evolutionary innovation in the common ancestor of the Elegans supergroup. To investigate the potential origin of this novel exon, we searched for any genomic sequences resembling the *C. elegans* CSR-1a exon in several species outside of the Elegans supergroup (*C. uteleia*, *C.* sp. 45, *C. astrocarya*, *C. castelli*, and *C.* sp. 27), but we were unable to find any significantly similar sequences.

Given that a subset of *Caenorhabditis* species evolved the novel CSR-1a exon, we wondered whether its origin was accompanied by sequence changes in the conserved CSR-1b portion of the gene that could be indicative of specialization of isoform functionality. We therefore selected a single representative CSR-1 gene from each of the 51 *Caenorhabditis* species and performed branch-site tests for positive selection using aBSREL [52]. After testing each branch on the species tree (primarily based on the CSR-1b portion of the gene, as most of the CSR-1a exon could not be reliably aligned), we detected evidence of significant positive selection on the branch leading to the Elegans supergroup (*P* = 0.0033, dN/dS = 4.55 at 10% of tested sites), which is the same branch of the phylogeny where the CSR-1a exon is predicted to have originated (Fig. 6B). We additionally detected signals of positive selection in CSR-1 on the branch leading to the Elegans group of species within the Elegans supergroup, though at a much smaller proportion of sites (*P* = 0.0198, dN/dS = 72.0 at 1.4% of tested sites). Our results are nearly identical when sites from the CSR-1a exon are excluded entirely (Elegans supergroup branch: *P* = 0.0024, dN/dS = 4.60 at 10% of tested sites; Elegans group branch: *P* = 0.0173, dN/dS = 74.6 at 1.4% of tested sites), suggesting that this signal of CSR-1 positive selection is not due to the small proportion of CSR-1a sites that could be tested. We repeated the branch-site tests using all 58 CSR-1 orthologs (rather than only 1 per species) and the inferred topology of the CSR-1 gene tree, and again found significant support for positive selection on both the branch leading to the Elegans supergroup (*P* = 0.0266, dN/dS = 4.52 at 9.3% of tested sites) and the branch leading to the Elegans group (*P* = 0.0102, dN/dS = 56.3 at 1.6% of tested sites). These inferences of positive selection were robust to accounting for the potential confounding effect of synonymous-site substitution rate variation when analyzed using BUSTED [53, 54], for both the Elegans supergroup branch (1 ortholog per species: *P* = 0.0004; all 58 orthologs: *P* = 0.0139) and the Elegans group branch (1 ortholog per species: *P* = 0.0408; all 58 orthologs: *P* = 0.0079). Taken together, these molecular evolutionary tests suggest that the evolution of the novel CSR- 1a exon in the ancestor of the Elegans supergroup was also accompanied by significant sequence diversification in the CSR-1b portion of this gene, consistent with functional specialization of the two CSR-1 isoforms.

## Discussion

In this study, we identified over 1200 Argonaute genes across 51 species of *Caenorhabditis* nematodes, allowing for an in-depth examination of how small RNA regulatory pathways evolve in a single genus. While many studies of small RNAs in *Caenorhabditis* focus only on *C. elegans*, we find that *Caenorhabditis* species can have fewer than half as many Argonaute genes as *C. elegans*, or more than twice as many, illustrating how dynamic small RNA pathways can be. Argonaute gene gains and losses were more common in some subfamilies than others, with some subfamilies (e.g., PRG-1) occasionally and repeatedly being lost entirely. We also identified and characterized notable instances of diversification, such as the gain of a potentially novel subfamily of ALGs in *C. panamensis*, the evolutionary birth of the microRNA-associated ALG-5 near the root of the *Caenorhabditis* phylogeny, as well as the evolution of different CSR-1 splice isoforms in the Elegans supergroup of species. Such diversity of gene regulatory mechanisms between species could contribute to the evolution of species-specific profiles of gene expression controls over development and genome integrity. For example, given that expression of ALG-5 is restricted to the germline (unlike the other microRNA-binding ALGs, ALG-1 and ALG-2) where it regulates the developmental timing of germ cells [7, 14], the gain of ALG-5 may represent a novel microRNA-mediated mechanism for regulating germline development in these species.

We found that some Argonaute subfamilies (e.g., ERGO-1) show substantial between-species gene copy number variation, but the question remains as to what evolutionary forces and selective pressures might drive such variation. Shi et al. [49] observed that the genomes of the outcrossing species *C. remanei* and *C. brenneri* contained more piRNA loci and Argonaute genes than the self-fertilizing species *C. elegans* and *C. briggsae*, and speculated that having more piRNAs and Argonautes was a defense against foreign transposable elements that would be encountered more frequently by outcrossing species. Reproductive mode itself cannot fully explain Argonaute copy number variation, however, as we found that the self-fertilizing species *C. tropicalis* has more Argonaute genes (27) than many outcrossing species, and that all of the species with the fewest Argonaute copies are outcrossing (Fig. 2A). Despite these observations, it is possible that control over transposon abundance exerts an influence on Argonaute copy number variation for at least some Argonautes, as we found that species with more orthologs of Argonautes in the WAGO-9/10/12 subfamily had significantly less repetitive genomes. Consistent with this possibility, WAGO-9/HRDE-1, WAGO-10, and WAGO-12/NRDE-3 are known to target pseudogenes and transposons for regulation [7, 40, 41], and a missense variant in WAGO-12 was found to be significantly associated with intraspecies variation in MIRAGE1 transposon abundance in *C. elegans* [55]. However, we did not detect this association between Argonaute gene copy number and genomic repeat content for other Argonaute subfamilies that target transposons, such as WAGO-1/3/4/5. Additionally, the piRNA Argonaute PRG-1 shows similar levels of between-species copy number variation as the microRNA Argonautes ALG-1 and ALG-2 (Table 1), despite the fact that transposons are thought to be targeted by piRNAs but not microRNAs in *Caenorhabditis* [9, 56]. Control of transposable elements is therefore likely not the sole explanation for differences in Argonaute copy number between species. Evolution of Argonaute copy number may be a response in part to the unique stressors and pathogens encountered by *Caenorhabditis* species, which occupy distinct habitats across the world [57, 58]. In line with this idea, immune response genes in *C. elegans* are enriched among targets of Argonautes ERGO-1, WAGO-6/SAGO-2, WAGO-8/SAGO-1, and WAGO-12/NRDE-3 [7]. As we observed the most significant copy number variation in the ERGO-1, WAGO-6/7/8, and WAGO-9/10/12 subfamilies, this Argonaute variation might correspond to the evolution of stress responses in different species, though this possibility needs to be tested further.

We observed that the piRNA Argonaute PRG-1, and likely the piRNA pathway as a whole, are absent in four of our studied *Caenorhabditis* species (*C. plicata*, *C. astrocarya*, *C. drosophilae*, and *C.* sp. 2). Given the topology of the species tree that we used (Fusca et al., in preparation), this likely represents three independent losses of the piRNA pathway across the genus. However, errors in our inferred species tree could theoretically alter this interpretation. The topology of our species tree is highly consistent with published *Caenorhabditis* phylogenies [31, 37, 59], though a few areas of the phylogeny carry some degree of uncertainty. For example, the exact placement of *C. virilis* on the phylogeny differs depending on the phylogenetic reconstruction method used [31, 37], which could affect the placement of its close relatives *C. drosophilae* and *C.* sp. 2. There also is uncertainty around the placement of *C. astrocarya*, owing to very short internal branch lengths (Fusca et al., in preparation). Despite these issues, there is currently no evidence that *C. plicata* or *C. astrocarya* form a clade with the sister species *C. drosophilae*/*C.* sp. 2 or with each other, and so having 3 independent piRNA pathway losses is currently the most parsimonious explanation for our results. We additionally found that some of the earliest-diverging *Caenorhabditis* species (e.g., *C. monodelphis*) have very low incidences of the piRNA Ruby motif, despite them still encoding PRG-1 (Fig. 5B). While one possible interpretation is that the Ruby motif was absent at the root of *Caenorhabditis* and then gained later on, it is more likely that these early-diverging species have lost the Ruby motif because nematode taxa outside of *Caenorhabditis* (e.g., *Diploscapter*, *Haemonchus*) also employ the core of this sequence motif upstream of piRNAs [34]. Interestingly, one of the early-diverging *Caenorhabditis* with low Ruby motif presence (*C. parvicauda*) also appears to have lost the piRNA processing factor PARN-1, despite retaining PRG-1, suggesting the possibility of a compromised piRNA pathway in this species as well.

In general, we observed that Argonaute subfamilies that showed more variable copy numbers between species also tended to contain more pseudogene-causing alleles in strains of *C. elegans* and *C. briggsae* (Fig. 3A, B). However, our detection of pseudogene alleles was limited only to coding sequences (e.g., frameshift mutations leading to premature stop codons), and thus we did not count potential pseudogene alleles that disrupt noncoding regulatory regions such as promoters. It is likely that such noncoding pseudogene alleles play a role in Argonaute functional evolution, given that the N2 strain of *C. elegans* does not appear to express WAGO-5 [7] despite an apparent lack of pseudogene-causing alleles in its coding sequence. Additionally, some pseudogene-causing alleles might not be strongly deleterious if they occur in exons that are not constitutively expressed due to alternative splicing. This may be the case for ALG-2, where all detected pseudogene alleles in *C. elegans* occurred in a region of coding sequence that is included in the longer ALG-2a isoform but not the shorter ALG-2b isoform. Further studies on the *cis*-regulatory motifs that control Argonaute expression in *Caenorhabditis* will allow for a more complete picture of how these genes gain and lose function over different evolutionary timescales.

An important consideration for this study is the quality of the underlying genome assemblies. For instance, we found that *C. japonica* has an uncharacteristically low incidence of Ruby motifs in its genome compared to other members of the Elegans supergroup (Fig. 5B).

However, given that the *C. japonica* genome assembly also has the highest amount of missing BUSCO genes of any Elegans supergroup species (11.1% missing), it is plausible that the rarity of Ruby motifs reflects an artifact of genome assembly for this repeat-rich genome (47.6% of the assembly consists of repeats). Apparent species-specific gene losses can be caused by issues with genome sequencing and assembly of the underlying genomic region, as was the case for the CSR-1 locus in *C. parvicauda* that we detected only from RNA-seq data. An opposing challenge is that apparent gene duplications might in reality correspond to multiple alleles of a single gene that were assembled improperly (as we corrected for in *C. brenneri*). We also observed errors in annotated gene models (e.g., erroneous fusions of adjacent genes into a single gene) that could bias analyses of sequence evolution if not properly accounted for. While we have accounted for many such genome quality issues in our analysis (such as by excluding species with the lowest-quality genome assemblies for some analyses that may be sensitive to these outliers), and in light of plausible biological copy number variation within species, it is likely that some Argonaute genes copies remain to be discovered. Current efforts by the research community to generate chromosome-scale genome assemblies for *Caenorhabditis* and other nematodes [60, 61, 62] will greatly assist future studies of comparative and population genomics in these species.

The computational predictions we provide serve as starting points for further discoveries about small RNA function. We identified the existence of a group of highly divergent ALG-like Argonautes in *C. panamensis* that may represent a novel subfamily, but it remains to be determined when and in which tissues these genes get expressed, which classes of small RNA they might bind to (e.g., microRNAs as is the case for ALG-1, ALG-2, and ALG-5, or 26G- RNAs as is the case for ALG-3 and ALG-4 [7]), and what functional role they might play in gene regulation or other molecular programs. Moreover, experimental validation of our prediction that the piRNA pathway has been lost repeatedly in different species will require sequencing their expressed small RNAs to assess the presence or absence of piRNA (i.e., 21U- RNA) expression. Our observation that CSR-1 appears to have diversified from a single CSR-1b- like isoform into separate CSR-1a and CSR-1b isoforms also provides several interesting possibilities for further study. In *C. elegans*, CSR-1a and CSR-1b show differing patterns of spatiotemporal expression: CSR-1a in spermatogenic tissue and some somatic tissues such as the intestine, and CSR-1b constitutively in the germline [50]. Our understanding the evolution of CSR-1 would therefore benefit from profiling its expression in species outside of the Elegans supergroup that we predict to express only a single CSR-1b-like isoform: is the ancestral CSR-1 expressed in both the soma and the germline, taking on the role of both CSR-1a and CSR-1b, or is it limited to the germline like CSR-1b? Such experimental work will help to confirm and expand upon the results presented here. In conclusion, our comprehensive survey of Argonaute genes across *Caenorhabditis* has revealed substantial functional diversity with compelling implications for mechanisms of gene expression regulation that also provides a valuable resource for comparative studies of gene regulatory evolution between species.

## Materials & Methods

### Identification of Argonaute genes

Reference genome assemblies for 50 *Caenorhabditis* species, as well as a reference transcriptome for 1 species (*C.* sp. 45), were obtained from a combination of NCBI GenBank [63], WormBase ParaSite [64, 65], and the Caenorhabditis Genomes Project [66] (Table S3). We assessed the quality of all 51 genome and transcriptome assemblies by running BUSCO v5.2.2 [67] to detect the single-copy genes in the nematoda_odb10 gene set (Table S2). Full details on all methods can be found in the Supplementary Methods (Appendix S1).

We defined groups of orthologous genes (orthogroups) between all 51 *Caenorhabditis* species with OrthoFinder v2.5.2 [68]. From the output of OrthoFinder, we took any protein belonging to the same orthogroup as a known *C. elegans* Argonaute [9] to be a candidate Argonaute. Additionally, proteins that were not identified as Argonautes by OrthoFinder, but that contained at least 1 domain characteristic of Argonaute proteins (“Piwi domain”, “PAZ domain”, “Mid domain of argonaute”, “N-terminal domain of argonaute”, “Argonaute linker 1 domain”, or “Argonaute linker 2 domain”) according to InterProScan v5.52-86.0 [69] were added to our set of Argonautes, and assigned to one of the 11 Argonaute subfamilies when possible. Identified Argonautes were further filtered based on length to remove short gene fragments. A maximum likelihood gene tree of all 1213 identified Argonautes that passed our length filtering was inferred with IQ-TREE v2.1.2 [70–72] after creating a multiple sequence alignment of their protein sequences using the Clustal Omega web server [73].

We used our identified Argonaute genes to count the number of Argonautes from all 11 subfamilies in each of the 51 species, correcting our counts for identified issues of genome assembly and gene annotation (see Supplementary Methods for details). Briefly, we identified 4 cases where a species did not appear to have a certain type of Argonaute gene that passed our length cutoff, but which had one or more gene fragments resembling the “missing” Argonaute – for counting purposes we considered these species to have 1 copy of the corresponding Argonaute. We also identified 2 cases where a pair of tandem duplicate genes was erroneously annotated as a single fused gene, which we counted as separate genes. Lastly, we found 9 pairs of Argonautes in *C. brenneri* that appeared to be allelic variants of the same gene based on protein sequence identity and synonymous substitution rate, and we therefore only counted one Argonaute from each of these 9 pairs (Table S4, Fig. S11). Apparent losses of an Argonaute subfamily in a given species were verified by using tblastn v2.9.0 [74] to search for the gene in the corresponding genome sequence.

### Within-species Argonaute sequence variation

Within-species variation data for *C. elegans* and *C. briggsae* Argonautes were downloaded from the Variant Annotation tool on CaeNDR release 20220216 (for *C. elegans*) and release 20240129 (for *C. briggsae*) [32]. For each Argonaute gene in both species, we retrieved all unique variants annotated as being “synonymous” or “missense” (i.e., non-synonymous), or whose annotations suggest they may be pseudogene-causing (all unique variants carrying the annotation “frameshift”, “start_lost”, “stop_gained”, “stop_lost”, “splice_acceptor”, or “splice_donor”). For each Argonaute in both *C. elegans* and *C. briggsae*, we calculated the number of unique isotypes carrying at least 1 variant of each type (synonymous, non-synonymous, and potentially pseudogene-causing). We additionally calculated per-site Watterson’s θ (the normalized number of variants per site) at both synonymous and non-synonymous sites for each Argonaute, using degenotate v1.3 (https://github.com/harvardinformatics/degenotate) to count the number of 0-fold, 2-fold, 3-fold, and 4-fold degenerate sites in the coding sequence of each Argonaute.

### 3D structural analysis

Files of computationally-predicted 3D structures for all 20 *C. elegans* Argonaute proteins, as well as *C. elegans* DCR-1 (for use as an outgroup) were downloaded from the AlphaFold Protein Structure Database [75] (Table S5). For each of the 7 *C. panamensis* Argonaute proteins that potentially belong to a novel subfamily of ALGs (as well as CSR-1 from *C. panamensis* as a control), we generated AlphaFold predictions of their 3D structures using ColabFold v1.5.5, by running the AlphaFold2_mmseqs2 Google Colab notebook with default settings [42, 43]. Prior to structure prediction, we manually trimmed the protein sequences of two of these putatively novel Argonautes (CSP28.g8935.t1 and CSP28.g16021.t1) to remove sequences from adjacent genes that were erroneously included in these gene models. To determine which Argonaute proteins had the most similar structures to our “novel” *C. panamensis* Argonautes, we used the DALI web server [76] to perform structural alignments between all pairs of our 29 protein structures (21 from *C. elegans* and 8 from *C. panamensis*), and used the reported Z-scores for each alignment as our measure of structural similarity. We additionally used these DALI structural alignments to assess whether each protein structure contained an N, PAZ, PIWI, and MID domain based on alignments to *C. elegans* ALG-1 (in which all 4 domain types were annotated). Heatmaps of protein sequence identity and structural similarity were created using ComplexHeatmap v2.12.1 [77], clustering rows and columns with hierarchical clustering based on sequence identity. 3D protein structures were visualized using the Mol* Viewer [78] on RCSB Protein Data Bank [79].

### RNA-seq analysis

Raw gene expression data from SRA was retrieved for 14 different *Caenorhabditis* species: *C. afra, C. agridulce*, *C. castelli*, *C. inopinata, C. macrosperma, C. monodelphis*, *C. nouraguensis, C. parvicauda*, *C. plicata*, *C. quiockensis*, *C. remanei*, *C. tribulationis, C. uteleia,* and *C. waitukubuli* (Table S6). The quality of each FASTQ file was assessed with FastQC v0.11.7 (https://www.bioinformatics.babraham.ac.uk/projects/fastqc/), and sequencing adapters were trimmed from reads using TrimGalore v0.6.6 (https://github.com/FelixKrueger/TrimGalore). Trimmed RNA-seq reads were aligned to their respective reference genomes using STAR v2.5.3a [80] and filtered with SAMtools v1.8 [81] to exclude reads that did not align uniquely, reads that did not align as a proper pair (for paired-end files), and duplicate reads (for paired-end files). Filtered RNA-seq read alignments at the CSR-1 locus for each species were visualized with the IGV web server [82]. Additionally, we used Trinity v2.13.2 [83] to make a de novo transcriptome assembly for each of the 14 species, using the trimmed FASTQ files as input. To verify apparent gene losses in species with available RNA-seq data (e.g., CSR-1 in *C. parvicauda*, PRG-1 in *C. plicata*), we used tblastn v2.9.0 [74] to search the corresponding transcriptome assembly for that gene.

### Worm culture and DNA isolation

Molecular biology confirmation was conducted in the following species: *C. drosophilae* (DF5112), *C. macrosperma* (JU1857), *C. panamensis* (QG702), *C. plicata* (SB355), and *C.* sp. 30 (DF5152). *C. elegans* (N2) and *C. virilis* (JU1528) were used as controls for species inside and outside the Elegans supergroup, respectively. All strains were cultured on 10 cm NGM-agar plates seeded with *Escherichia coli* OP50 and maintained at room temperature (∼24°C), except *C. elegans* which was maintained at 20°C [84, 85]. Prior to DNA extractions, strains were cleaned and age synchronized through hypochlorite treatment [85] or antibiotic treatment (100ug/mL carbenicillin and 400ug/mL kanamycin). A complete list of strain information can be found in Table S7.

High quality genomic DNA was purified using the Monarch Genomic DNA Purification Kit (NEB), rinsing and pelleting at least 1,000 adult worms. Following the manufacturer recommendations for animal tissues, 200uL of tissue lysis buffer and 10uL of Proteinase K were added to each sample, incubated in a shaking incubator at 56°C, and RNase-A treated. The eluted DNA was further cleaned using the DNA Clean and Concentrator-5 kit (Zymo). DNA concentration and purity was assessed using Nanodrop (Table S7).

Genomic DNA for *C. virilis* was purified by isolating 10 adult worms in 6uL of 20mg/uL Proteinase K in elution buffer. A total of eight samples (n = 80 worms) were collected. Samples were freeze-cracked at -80°C for 30 minutes two times. They were then incubated at 58°C for 60 minutes followed by 10 minutes at 95°C to inactivate the Proteinase K.

### Molecular biology

PCR products were generated using the Q5 High-Fidelity Master Mix (NEB) and 100ng of purified DNA in accordance with manufacturer instructions. We verified that the DNA isolation process was successful in all species by amplifying a 535bp product in the 18S ribosomal DNA region. A complete list of primers used in this study can be found in Table S8.

To verify the putative duplicate copy of *csr-1* in *C. macrosperma*, we attempted to amplify through the entire gene, starting 220bp upstream of the start codon through 154bp downstream of the stop codon. We additionally attempted to amplify exons 1-3 and exons 12-17 (i.e., the PIWI domain). PCR products of approximately the correct size were gel extracted and purified using the Zymoclean Gel DNA Recovery Kit (Zymo) for verification by Sanger sequencing.

To verify the putative duplicate copy of *prg-1* in *C.* sp. 30, we amplified through the entire gene, starting 219bp upstream of the start codon through 172bp downstream of the stop codon. Additionally, we amplified products starting upstream of the start codon through exon 5 and through exon 10. PCR products for all primer combinations were gel extracted and purified using the Zymoclean Gel DNA Recovery Kit (Zymo) for verification by Sanger sequencing.

To examine the potential loss of *prg-1* in *C. drosophilae* and *C. plicata*, we designed primers using a reduced alignment of seven species for which we had DNA along with *C. bovis*. From this alignment, we designed degenerate primers with no more than four ambiguous base pairs in each 20nt forward and reverse primer that amplified exons 4-8. PCRs using the degenerate primers were run at three temperatures: the lowest potential Tm, the highest potential Tm, and an intermediate Tm. All of the positive controls (*C. elegans*, *C. macrosperma*, *C. panamensis*, *C.* sp. 30, and *C. virilis*) amplified correctly for at least one temperature.

To verify the divergent *C. panamensis* ALG duplicate CSP28.g8935.t1, we first attempted to amplify a product starting 117bp upstream of the start codon through 106bp downstream of the stop codon. This reaction produced product. We then attempted to amplify the coding sequence only (exons 1-15). PCR products of approximately the correct size were gel extracted and purified using Zymoclean Gel DNA Recovery Kit (Zymo) for verification by Sanger sequencing. The remaining 6 divergent *C. panamensis* ALG duplicates (which formed a cluster of tandem duplicates) were analyzed using three approaches. First, we attempted to amplify between genes by anchoring the forward primer in the final exon of the upstream gene and the reverse primer in the first or second exon of the downstream gene. We additionally attempted to amplify ∼1kb upstream and downstream of each gene independently. Finally, we attempted to amplify the coding sequence of each gene for verification by Sanger sequencing.

### Gene family evolution

Significant gene family expansions and contractions across *Caenorhabditis* were detected with CAFE v5.1 [38], using the dated *Caenorhabditis* species tree from Fusca et al. (in preparation). The number of genes in each orthogroup reported by OrthoFinder were used as the gene family size for CAFE (aside from the 11 Argonaute orthogroups, for which we instead used our manually-corrected gene counts). As CAFE only tests gene families that are inferred to be present at the root of the species tree, we excluded *C. monodelphis* and *C. parvicauda* in order to test all 11 Argonaute families, since these early-diverging species lack copies of ALG-5 and ERGO-1 respectively. We additionally reran CAFE while also excluding the 5 species with the lowest-quality gene annotations according to BUSCO (either fewer than 85% of BUSCO genes complete and single-copy, or over 5% of BUSCO genes complete but duplicated) – these 5 species are *C. angaria*, *C. brenneri*, *C. japonica*, *C.* sp. 49, and *C. waitukubuli*. CAFE detected significant gene families using a P-value threshold of 0.05, and we additionally performed multiple-testing correction on these P-values by calculating False Discovery Rates in R v4.2.1 [86], taking gene families with an FDR ≤ 0.05 to be significant.

To model genomic repeat content as a function of Argonaute gene copy number, we identified repeat families in each of our 50 reference genome assemblies using RepeatModeler v2.0.3 [87] and RepeatMasker v4.1.2 (https://www.repeatmasker.org/). We defined repeat content for a species as the proportion of that species’ genome that was masked by RepeatMasker (Table S1). All phylogenetic generalized least squares (PGLS) regressions in this study were performed in R with caper v1.0.2 (https://cran.r-project.org/web/packages/caper/index.html), using our dated species tree as the phylogeny and excluding *C.* sp. 45 (as we only have a transcriptome assembly for this species). Genomic repeat content was modeled using the following two models: (1) Repeat content ∼ genome size + total number of non-Argonaute protein-coding genes + total number of Argonaute genes; and (2) Repeat content ∼ genome size + total number of non-Argonaute protein-coding genes + number of RDE-1 orthologs + number of ALG-1/2 orthologs + number of ALG-3/4 orthologs + number of ALG-5 orthologs + number of CSR-1 orthologs + number of ERGO-1 orthologs + number of PRG-1 orthologs + number of VSRA-1 orthologs + number of WAGO-1/3/4/5 orthologs + number of WAGO-6/7/8 orthologs + number of WAGO-9/10/12 orthologs. Both regressions were repeated after excluding the 5 species with the lowest-quality gene annotations (*C. angaria*, *C. brenneri*, *C. japonica*, *C.* sp. 49, and *C. waitukubuli*), and also repeated for three specific types of repeats (retrotransposons, DNA transposons, and rolling-circle transposons), taking the dependent variable to be the proportion of the genome annotated by RepeatMasker as belonging to that repeat type.

### piRNA pathway analysis

The presence or absence of 7 genes involved in piRNA transcription and processing (*prde-1*, *snpc-4*, *tofu-4*, *tofu-5*, *pid-1*, *henn-1*, and *parn-1*) [10] in all 51 species was determined based on whether OrthoFinder was able to identify any orthologs of these genes in each species, verifying losses by using tblastn v2.9.0 [74] to search for the gene in the corresponding genome sequence (as well as our assembled transcriptome, when possible). Ruby motifs were detected in the noncoding portion of all 50 available reference genomes (i.e., every species except for *C.* sp. 45) by first using the “maskfasta” function of BEDTools v2.30.0 [88] to mask all sequences belonging to exons for each genome. The “locate” function of SeqKit v2.3.0 [89] was then used to find and count all instances of the sequence “CNGTTTCA” (where “N” can be any nucleotide) and its reverse complement in each masked genome sequence, based on the known Ruby motif sequence in *Caenorhabditis* [34]. The number of expected Ruby motifs in each species was calculated as the probability of a random 8-mer in a species being “CNGTTTCA” or its reverse complement given the dinucleotide content of its noncoding genome, multiplied by the number of 8-mers in a DNA sequence as long as that species’ noncoding genome (i.e., noncoding genome size – 7). Genomic locations of known piRNA clusters in *C. elegans* and *C. briggsae* were taken from [47] and [49], respectively.

### CSR-1 sequence evolution

DNA coding sequences were retrieved for all identified CSR-1 orthologs, aside from very highly-diverged CSR-1 duplicates in *C. parvicauda* (CSP21.g381.t1) and *C. macrosperma* (CMACR.g1069.t1) which may be genome assembly artifacts, resulting in a total of 58 CSR-1 sequences. In addition to this set of 58 sequences, we also created a subset that contained only a single CSR-1 ortholog for each of the 51 species, by keeping only a single CSR-1 ortholog from the species that had multiple orthologs (*C. plicata*, *C. brenneri*, *C.* sp. 48, *C.* sp. 54, and *C. doughertyi*). Both sets of CSR-1 sequences were separately aligned using PRANK v.170427 [90] in codon-alignment mode. Codons that could not be aligned reliably were filtered out from both alignments with GUIDANCE v2.0.2 [91], using 50 bootstrap runs and a cutoff score of 0.93. For the alignment containing all 58 CSR-1 sequences, a maximum-likelihood gene tree was constructed by running IQ-TREE v2.1.2 [70–72].

Branch-site tests of positive selection were performed on both the full 58-sequence alignment and the reduced 51-sequence alignment using both aBSREL v2.5 [52] and BUSTED v4.1 [54], hosted on the Datamonkey web server [92]. For the corresponding phylogeny needed for these analyses, we used our CSR-1 gene tree for the full 58-sequence alignment, and our *Caenorhabditis* species tree for the 51-sequence alignment. The gene structure of CSR-1 across the genus was visualized using TBtools v1.119 [93], again taking only a single representative CSR-1 ortholog for species with multiple orthologs, exactly as was done for our branch-site tests. The locations of arginine-glycine (RG) repeats in each CSR-1 sequence were detected using the “locate” function of SeqKit v2.3.0 [89]. For this visualization, we manually corrected the CSR-1 gene models for 6 Elegans supergroup species in which the CSR-1a exon was present but not annotated properly (see Supplementary Methods for details). We used tblastn v2.9.0 and blastp v2.9.0 [74] to search for the first 163 amino acids of *C. elegans* CSR-1 (corresponding to the CSR-1a exon) in the genomes (or transcriptome, for *C.* sp. 45) and annotated proteomes of 5 species outside of the Elegans supergroup (*C. uteleia*, *C.* sp. 45, *C. astrocarya*, *C. castelli*, and *C.* sp. 27).

## Data availability

Supplementary data files and code associated with this study are available on our GitHub repository at https://github.com/Cutterlab/Caenorhabditis_AGO_Survey.

## Acknowledgements

We thank Lewis Stevens, Mark Blaxter, and all other members of the Caenorhabditis Genomes Project for prepublication access to sequenced genomes and transcriptomes. Computations were performed on the Niagara supercomputer at the SciNet HPC Consortium. We are grateful to Erik Andersen for assistance in using CaeNDR and Julie Claycomb for providing insights about Argonaute biology and thoughtful feedback on this manuscript. SciNet is funded by Innovation, Science and Economic Development Canada; the Digital Research Alliance of Canada; the Ontario Research Fund: Research Excellence; and the University of Toronto. A.D.C. is supported by Discovery Grant funds from the Natural Sciences and Engineering Research Council of Canada.

## Notes

https://github.com/Cutterlab/Caenorhabditis_AGO_Survey

### Competing Interest Statement

The authors have declared no competing interest.

## References

1. Swarts DC, Makarova K, Wang Y, Nakanishi K, Ketting RF, Koonin EV, et al. The evolutionary journey of Argonaute proteins. Nat Struct Mol Biol. 2014;21(9):743–53. pmid:25192263.

2. Ryazansky S, Kulbachinskiy A, Aravin AA. The Expanded Universe of Prokaryotic Argonaute Proteins. mBio. 2018;9(6). pmid:30563906.

3. Buckley BA, Burkhart KB, Gu SG, Spracklin G, Kershner A, Fritz H, et al. A nuclear Argonaute promotes multigenerational epigenetic inheritance and germline immortality. Nature. 2012;489(7416):447-51. pmid:22810588.

4. Conine CC, Moresco JJ, Gu W, Shirayama M, Conte D, Yates JR, et al. Argonautes promote male fertility and provide a paternal memory of germline gene expression in *C. elegans*. Cell. 2013;155(7):1532–44. pmid:24360276.

5. Seth M, Shirayama M, Gu W, Ishidate T, Conte D, Mello CC. The *C. elegans* CSR-1 Argonaute pathway counteracts epigenetic silencing to promote germline gene expression. Dev Cell. 2013;27(6):656–63. pmid:24360782.

6. Wedeles CJ, Wu MZ, Claycomb JM. Protection of germline gene expression by the *C. elegans* Argonaute CSR-1. Dev Cell. 2013;27(6):664–71. pmid:24360783.

7. Seroussi U, Lugowski A, Wadi L, Lao RX, Willis AR, Zhao W, et al. A comprehensive survey of *C. elegans* argonaute proteins reveals organism-wide gene regulatory networks and functions. Elife. 2023;12. pmid:36790166.

8. Hoogstrate SW, Volkers RJ, Sterken MG, Kammenga JE, Snoek LB. Nematode endogenous small RNA pathways. Worm. 2014;3:e28234. pmid:25340013.

9. Youngman EM, Claycomb JM. From early lessons to new frontiers: the worm as a treasure trove of small RNA biology. Front Genet. 2014;5:416. pmid:25505902.

10. Almeida MV, Andrade-Navarro MA, Ketting RF. Function and Evolution of Nematode RNAi Pathways. Noncoding RNA. 2019;5(1). pmid:30650636.

11. Yigit E, Batista PJ, Bei Y, Pang KM, Chen CC, Tolia NH, et al. Analysis of the *C. elegans* Argonaute family reveals that distinct Argonautes act sequentially during RNAi. Cell. 2006;127(4):747–57. pmid:17110334.

12. Wang G, Reinke V. A *C. elegans* Piwi, PRG-1, regulates 21U-RNAs during spermatogenesis. Curr Biol. 2008;18(12):861–7. pmid:18501605.

13. Conine CC, Batista PJ, Gu W, Claycomb JM, Chaves DA, Shirayama M, et al. Argonautes ALG-3 and ALG-4 are required for spermatogenesis-specific 26G-RNAs and thermotolerant sperm in *Caenorhabditis elegans*. Proc Natl Acad Sci U S A. 2010;107(8):3588–93. pmid:20133686.

14. Brown KC, Svendsen JM, Tucci RM, Montgomery BE, Montgomery TA. ALG-5 is a miRNA-associated Argonaute required for proper developmental timing in the *Caenorhabditis elegans* germline. Nucleic Acids Res. 2017;45(15):9093–107. pmid:28645154.

15. Wray GA. The evolutionary significance of *cis*-regulatory mutations. Nat Rev Genet. 2007;8(3):206–16. pmid:17304246.

16. Romero IG, Ruvinsky I, Gilad Y. Comparative studies of gene expression and the evolution of gene regulation. Nat Rev Genet. 2012;13(7):505–16. pmid:22705669.

17. Signor SA, Nuzhdin SV. The Evolution of Gene Expression in *cis* and *trans*. Trends Genet. 2018;34(7):532–44. pmid:29680748.

18. Simkin A, Wong A, Poh YP, Theurkauf WE, Jensen JD. Recurrent and recent selective sweeps in the piRNA pathway. Evolution. 2013;67(4):1081–90. pmid:23550757.

19. Jovelin R, Cutter AD. Microevolution of nematode miRNAs reveals diverse modes of selection. Genome Biol Evol. 2014;6(11):3049–63. pmid:25355809.

20. Mohammed J, Bortolamiol-Becet D, Flynt AS, Gronau I, Siepel A, Lai EC. Adaptive evolution of testis-specific, recently evolved, clustered miRNAs in *Drosophila*. RNA. 2014;20(8):1195–209. pmid:24942624.

21. Palmer WH, Hadfield JD, Obbard DJ. RNA-Interference Pathways Display High Rates of Adaptive Protein Evolution in Multiple Invertebrates. Genetics. 2018;208(4):1585–99. pmid:29437826.

22. Kelley JL, Desvignes T, McGowan KL, Perez M, Rodriguez LA, Brown AP, et al. microRNA expression variation as a potential molecular mechanism contributing to adaptation to hydrogen sulphide. J Evol Biol. 2021;34(6):977–88. pmid:33124163.

23. Fusca DD, Sharma E, Weiss JG, Claycomb JM, Cutter AD. Temperature-dependent Small RNA Expression Depends on Wild Genetic Backgrounds of *Caenorhabditis briggsae*. Mol Biol Evol. 2022;39(11). pmid:36223483.

24. Buck AH, Blaxter M. Functional diversification of Argonautes in nematodes: an expanding universe. Biochem Soc Trans. 2013;41(4):881–6. pmid:23863149.

25. Sasaki T, Shiohama A, Minoshima S, Shimizu N. Identification of eight members of the Argonaute family in the human genome. Genomics. 2003;82(3):323–30. pmid:12906857.

26. Lewis SH, Salmela H, Obbard DJ. Duplication and Diversification of Dipteran Argonaute Genes, and the Evolutionary Divergence of Piwi and Aubergine. Genome Biol Evol. 2016;8(3):507–18. pmid:26868596.

27. Nong W, Cao J, Li Y, Qu Z, Sun J, Swale T, et al. Jellyfish genomes reveal distinct homeobox gene clusters and conservation of small RNA processing. Nat Commun. 2020;11(1):3051. pmid:32561724.

28. Aravin AA, Sachidanandam R, Girard A, Fejes-Toth K, Hannon GJ. Developmentally regulated piRNA clusters implicate MILI in transposon control. Science. 2007;316(5825):744-7. pmid:17446352.

29. Wang SH, Elgin SC. *Drosophila* Piwi functions downstream of piRNA production mediating a chromatin-based transposon silencing mechanism in female germ line. Proc Natl Acad Sci U S A. 2011;108(52):21164–9. pmid:22160707.

30. Sarkies P, Selkirk ME, Jones JT, Blok V, Boothby T, Goldstein B, et al. Ancient and novel small RNA pathways compensate for the loss of piRNAs in multiple independent nematode lineages. PLoS Biol. 2015;13(2):e1002061. pmid:25668728.

31. Stevens L, Félix MA, Beltran T, Braendle C, Caurcel C, Fausett S, et al. Comparative genomics of 10 new *Caenorhabditis* species. Evol Lett. 2019;3(2):217–36. pmid:31007946.

32. Crombie TA, McKeown R, Moya ND, Evans KS, Widmayer SJ, LaGrassa V, et al. CaeNDR, the *Caenorhabditis* Natural Diversity Resource. Nucleic Acids Res. 2024;52(D1):D850–D8. pmid:37855690.

33. Kanzaki N, Tsai IJ, Tanaka R, Hunt VL, Liu D, Tsuyama K, et al. Biology and genome of a newly discovered sibling species of *Caenorhabditis elegans*. Nat Commun. 2018;9(1):3216. pmid:30097582.

34. Beltran T, Barroso C, Birkle TY, Stevens L, Schwartz HT, Sternberg PW, et al. Comparative Epigenomics Reveals that RNA Polymerase II Pausing and Chromatin Domain Organization Control Nematode piRNA Biogenesis. Dev Cell. 2019;48(6):793–810.e6. pmid:30713076.

35. Chou HT, Valencia F, Alexander JC, Bell AD, Deb D, Pollard DA, et al. Diversification of small RNA pathways underlies germline RNA interference incompetence in wild *Caenorhabditis elegans* strains. Genetics. 2024;226(1). pmid:37865119.

36. Ahmed M, Roberts N, Adediran F, Smythe A, Kocot K, Holovachov O. Phylogenomic Analysis of the Phylum Nematoda: Conflicts and Congruences With Morphology, 18S rRNA, and Mitogenomes. Frontiers in Ecology and Evolution. 2022;9. WOS:000760536100001.

37. Sloat SA, Noble LM, Paaby AB, Bernstein M, Chang A, Kaur T, et al. *Caenorhabditis* nematodes colonize ephemeral resource patches in neotropical forests. Ecol Evol. 2022;12(7):e9124. pmid:35898425.

38. Mendes FK, Vanderpool D, Fulton B, Hahn MW. CAFE 5 models variation in evolutionary rates among gene families. Bioinformatics. 2021;36(22-23):5516–8. pmid:33325502.

39. Vasale JJ, Gu W, Thivierge C, Batista PJ, Claycomb JM, Youngman EM, et al. Sequential rounds of RNA-dependent RNA transcription drive endogenous small-RNA biogenesis in the ERGO-1/Argonaute pathway. Proc Natl Acad Sci U S A. 2010;107(8):3582–7. pmid:20133583.

40. de Albuquerque BF, Placentino M, Ketting RF. Maternal piRNAs Are Essential for Germline Development following De Novo Establishment of Endo-siRNAs in *Caenorhabditis elegans*. Dev Cell. 2015;34(4):448–56. pmid:26279485.

41. Padeken J, Methot S, Zeller P, Delaney CE, Kalck V, Gasser SM. Argonaute NRDE-3 and MBT domain protein LIN-61 redundantly recruit an H3K9me3 HMT to prevent embryonic lethality and transposon expression. Genes Dev. 2021;35(1-2):82–101. pmid:33303642.

42. Jumper J, Evans R, Pritzel A, Green T, Figurnov M, Ronneberger O, et al. Highly accurate protein structure prediction with AlphaFold. Nature. 2021;596(7873):583-9. pmid:34265844.

43. Mirdita M, Schütze K, Moriwaki Y, Heo L, Ovchinnikov S, Steinegger M. ColabFold: making protein folding accessible to all. Nat Methods. 2022;19(6):679–82. pmid:35637307.

44. Batista PJ, Ruby JG, Claycomb JM, Chiang R, Fahlgren N, Kasschau KD, et al. PRG-1 and 21U-RNAs interact to form the piRNA complex required for fertility in *C. elegans*. Mol Cell. 2008;31(1):67–78. pmid:18571452.

45. Das PP, Bagijn MP, Goldstein LD, Woolford JR, Lehrbach NJ, Sapetschnig A, et al. Piwi and piRNAs act upstream of an endogenous siRNA pathway to suppress Tc3 transposon mobility in the *Caenorhabditis elegans* germline. Mol Cell. 2008;31(1):79–90. pmid:18571451.

46. Tang W, Tu S, Lee HC, Weng Z, Mello CC. The RNase PARN-1 Trims piRNA 3’ Ends to Promote Transcriptome Surveillance in *C. elegans*. Cell. 2016;164(5):974–84. pmid:26919432.

47. Ruby JG, Jan C, Player C, Axtell MJ, Lee W, Nusbaum C, et al. Large-scale sequencing reveals 21U-RNAs and additional microRNAs and endogenous siRNAs in *C. elegans*. Cell. 2006;127(6):1193–207. pmid:17174894.

48. Weng C, Kosalka J, Berkyurek AC, Stempor P, Feng X, Mao H, et al. The USTC co-opts an ancient machinery to drive piRNA transcription in *C. elegans*. Genes Dev. 2019;33(1- 2):90–102. pmid:30567997.

49. Shi Z, Montgomery TA, Qi Y, Ruvkun G. High-throughput sequencing reveals extraordinary fluidity of miRNA, piRNA, and siRNA pathways in nematodes. Genome Res. 2013;23(3):497–508. pmid:23363624.

50. Charlesworth AG, Seroussi U, Lehrbach NJ, Renaud MS, Sundby AE, Molnar RI, et al. Two isoforms of the essential *C. elegans* Argonaute CSR-1 differentially regulate sperm and oocyte fertility. Nucleic Acids Res. 2021;49(15):8836–65. pmid:34329465.

51. Nguyen DAH, Phillips CM. Arginine methylation promotes siRNA-binding specificity for a spermatogenesis-specific isoform of the Argonaute protein CSR-1. Nat Commun. 2021;12(1):4212. pmid:34244496.

52. Smith MD, Wertheim JO, Weaver S, Murrell B, Scheffler K, Kosakovsky Pond SL. Less is more: an adaptive branch-site random effects model for efficient detection of episodic diversifying selection. Mol Biol Evol. 2015;32(5):1342–53. pmid:25697341.

53. Wisotsky SR, Kosakovsky Pond SL, Shank SD, Muse SV. Synonymous Site-to-Site Substitution Rate Variation Dramatically Inflates False Positive Rates of Selection Analyses: Ignore at Your Own Peril. Mol Biol Evol. 2020;37(8):2430–9. pmid:32068869.

54. Lucaci AG, Zehr JD, Enard D, Thornton JW, Kosakovsky Pond SL. Evolutionary Shortcuts via Multinucleotide Substitutions and Their Impact on Natural Selection Analyses. Mol Biol Evol. 2023;40(7). pmid:37395787.

55. Laricchia KM, Zdraljevic S, Cook DE, Andersen EC. Natural Variation in the Distribution and Abundance of Transposable Elements Across the *Caenorhabditis elegans* Species. Mol Biol Evol. 2017;34(9):2187–202. pmid:28486636.

56. Ambros V, Ruvkun G. Recent Molecular Genetic Explorations of *Caenorhabditis elegans* MicroRNAs. Genetics. 2018;209(3):651–73. pmid:29967059.

57. Cutter AD. *Caenorhabditis* evolution in the wild. Bioessays. 2015;37(9):983–95. pmid:26126900.

58. Schulenburg H, Félix MA. The Natural Biotic Environment of *Caenorhabditis elegans*. Genetics. 2017;206(1):55–86. pmid:28476862.

59. Sun S, Kanzaki N, Dayi M, Maeda Y, Yoshida A, Tanaka R, et al. The compact genome of *Caenorhabditis niphades* n. sp., isolated from a wood-boring weevil, *Niphades variegatus*. BMC Genomics. 2022;23(1):765. pmid:36418933.

60. Gonzalez de la Rosa PM, Thomson M, Trivedi U, Tracey A, Tandonnet S, Blaxter M. A telomere-to-telomere assembly of *Oscheius tipulae* and the evolution of rhabditid nematode chromosomes. G3 (Bethesda). 2021;11(1). pmid:33561231.

61. Stevens L, Moya ND, Tanny RE, Gibson SB, Tracey A, Na H, et al. Chromosome-Level Reference Genomes for Two Strains of *Caenorhabditis briggsae*: An Improved Platform for Comparative Genomics. Genome Biol Evol. 2022;14(4). pmid:35348662.

62. Kieninger M, Stevens L, Collins JC, Wellcome Sanger Institute Tree of Life Management, Samples and Laboratory team, Wellcome Sanger Institute Tree of Life Core Informatics team, Wellcome Sanger Institute Scientific Operations: Sequencing Operations, Blaxter M. The genome sequence of the nematode *Caenorhabditis drosophilae* (Rhabditida, Rhabditidae) (Kiontke, 1997). Wellcome Open Res. 2024;9:292. pmid:39114493.

63. Sayers EW, Bolton EE, Brister JR, Canese K, Chan J, Comeau DC, et al. Database resources of the national center for biotechnology information. Nucleic Acids Res. 2022;50(D1):D20–D6. pmid:34850941.

64. Howe KL, Bolt BJ, Shafie M, Kersey P, Berriman M. WormBase ParaSite - a comprehensive resource for helminth genomics. Mol Biochem Parasitol. 2017;215:2–10. pmid:27899279.

65. Davis P, Zarowiecki M, Arnaboldi V, Becerra A, Cain S, Chan J, et al. WormBase in 2022-data, processes, and tools for analyzing *Caenorhabditis elegans*. Genetics. 2022;220(4). pmid: 35134929.

66. Stevens L. Data associated with the Caenorhabditis Genomes Project (caenorhabditis.org); 2024 [cited 2024 July 4]. Database: Zenodo [Internet]. Available from: 10.5281/zenodo.12633738

67. Manni M, Berkeley MR, Seppey M, Simão FA, Zdobnov EM. BUSCO Update: Novel and Streamlined Workflows along with Broader and Deeper Phylogenetic Coverage for Scoring of Eukaryotic, Prokaryotic, and Viral Genomes. Mol Biol Evol. 2021;38(10):4647–54. pmid:34320186.

68. Emms DM, Kelly S. OrthoFinder: phylogenetic orthology inference for comparative genomics. Genome Biol. 2019;20(1):238. pmid:31727128.

69. Jones P, Binns D, Chang HY, Fraser M, Li W, McAnulla C, et al. InterProScan 5: genome-scale protein function classification. Bioinformatics. 2014;30(9):1236–40. pmid:24451626.

70. Kalyaanamoorthy S, Minh BQ, Wong TKF, von Haeseler A, Jermiin LS. ModelFinder: fast model selection for accurate phylogenetic estimates. Nat Methods. 2017;14(6):587–9. pmid:28481363.

71. Hoang D, Chernomor O, von Haeseler A, Minh B, Vinh L. UFBoot2: Improving the Ultrafast Bootstrap Approximation. Molecular Biology and Evolution. 2018;35(2):518–22. pmid:29077904.

72. Minh BQ, Schmidt HA, Chernomor O, Schrempf D, Woodhams MD, von Haeseler A, et al. IQ-TREE 2: New Models and Efficient Methods for Phylogenetic Inference in the Genomic Era. Mol Biol Evol. 2020;37(5):1530–4. pmid:32011700.

73. Madeira F, Pearce M, Tivey ARN, Basutkar P, Lee J, Edbali O, et al. Search and sequence analysis tools services from EMBL-EBI in 2022. Nucleic Acids Res. 2022;50(W1):W276–W9. pmid:35412617.

74. Camacho C, Coulouris G, Avagyan V, Ma N, Papadopoulos J, Bealer K, et al. BLAST+: architecture and applications. BMC Bioinformatics. 2009;10:421. pmid:20003500.

75. Varadi M, Anyango S, Deshpande M, Nair S, Natassia C, Yordanova G, et al. AlphaFold Protein Structure Database: massively expanding the structural coverage of protein-sequence space with high-accuracy models. Nucleic Acids Res. 2022;50(D1):D439–D44. pmid:34791371.

76. Holm L. Dali server: structural unification of protein families. Nucleic Acids Res. 2022;50(W1):W210–W5. pmid:35610055.

77. Gu Z, Eils R, Schlesner M. Complex heatmaps reveal patterns and correlations in multidimensional genomic data. Bioinformatics. 2016;32(18):2847–9. pmid:27207943.

78. Sehnal D, Bittrich S, Deshpande M, Svobodová R, Berka K, Bazgier V, et al. Mol* Viewer: modern web app for 3D visualization and analysis of large biomolecular structures. Nucleic Acids Res. 2021;49(W1):W431–W7. pmid:33956157.

79. Burley SK, Bhikadiya C, Bi C, Bittrich S, Chao H, Chen L, et al. RCSB Protein Data Bank (RCSB.org): delivery of experimentally-determined PDB structures alongside one million computed structure models of proteins from artificial intelligence/machine learning. Nucleic Acids Res. 2023;51(D1):D488–D508. pmid:36420884.

80. Dobin A, Davis CA, Schlesinger F, Drenkow J, Zaleski C, Jha S, et al. STAR: ultrafast universal RNA-seq aligner. Bioinformatics. 2013;29(1):15–21. pmid:23104886.

81. Li H, Handsaker B, Wysoker A, Fennell T, Ruan J, Homer N, et al. The Sequence Alignment/Map format and SAMtools. Bioinformatics. 2009;25(16):2078–9. pmid:19505943.

82. Robinson JT, Thorvaldsdóttir H, Winckler W, Guttman M, Lander ES, Getz G, et al. Integrative genomics viewer. Nat Biotechnol. 2011;29(1):24–6. pmid:21221095.

83. Grabherr MG, Haas BJ, Yassour M, Levin JZ, Thompson DA, Amit I, et al. Full-length transcriptome assembly from RNA-Seq data without a reference genome. Nat Biotechnol. 2011;29(7):644–52. pmid:21572440.

84. Brenner S. The genetics of *Caenorhabditis elegans*. Genetics. 1974;77(1):71–94. pmid:4366476.

85. Kenyon C. The nematode *Caenorhabditis elegans*. Science. 1988;240(4858):1448-53. pmid:3287621.

86. R Core Team. R: A language and environment for statistical computing. R Foundation for Statistical Computing, Vienna, Austria. 2022. URL https://www.R-project.org/.

87. Flynn JM, Hubley R, Goubert C, Rosen J, Clark AG, Feschotte C, et al. RepeatModeler2 for automated genomic discovery of transposable element families. Proc Natl Acad Sci U S A. 2020;117(17):9451–7. pmid:32300014.

88. Quinlan AR, Hall IM. BEDTools: a flexible suite of utilities for comparing genomic features. Bioinformatics. 2010;26(6):841–2. pmid:20110278.

89. Shen W, Le S, Li Y, Hu F. SeqKit: A Cross-Platform and Ultrafast Toolkit for FASTA/Q File Manipulation. PLoS One. 2016;11(10):e0163962. pmid:27706213.

90. Löytynoja A, Goldman N. Phylogeny-aware gap placement prevents errors in sequence alignment and evolutionary analysis. Science. 2008;320(5883):1632-5. pmid:18566285.

91. Sela I, Ashkenazy H, Katoh K, Pupko T. GUIDANCE2: accurate detection of unreliable alignment regions accounting for the uncertainty of multiple parameters. Nucleic Acids Res. 2015;43(W1):W7–14. pmid:25883146.

92. Weaver S, Shank S, Spielman S, Li M, Muse S, Pond S. Datamonkey 2.0: A Modern Web Application for Characterizing Selective and Other Evolutionary Processes. Molecular Biology and Evolution. 2018;35(3):773–7. pmid:29301006.

93. Chen C, Chen H, Zhang Y, Thomas HR, Frank MH, He Y, et al. TBtools: An Integrative Toolkit Developed for Interactive Analyses of Big Biological Data. Mol Plant. 2020;13(8):1194–202. pmid:32585190.

