## Appendix S1 for "Dynamic birth and death of Argonaute gene family functional repertoire across *Caenorhabditis* nematodes"

Supplementary Figures

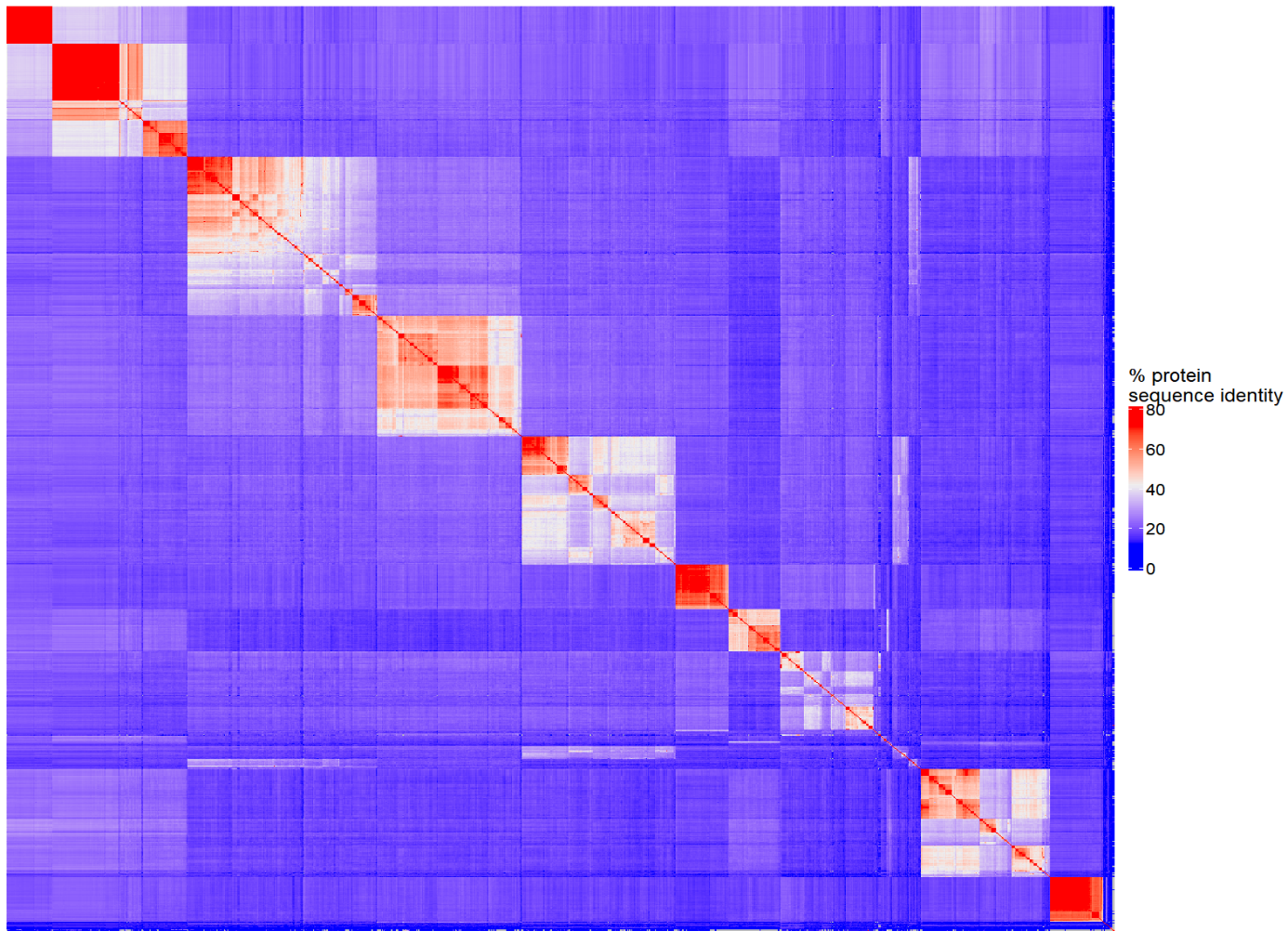

**ALG-5**

**ALG-3/4**

**ALG-1/2**

**WAGO-6/7/8**

**WAGO-1/3/4/5**

**WAGO-9/10/12**

**CSR-1**

**RDE-1**

**VSRA-1**

**ERGO-1**

**PRG-1**

**Figure S1.** Heatmap of protein sequence identity between all pairs of 1213 aligned Argonautes. The clusters corresponding to each of the 11 Argonaute families are indicated.

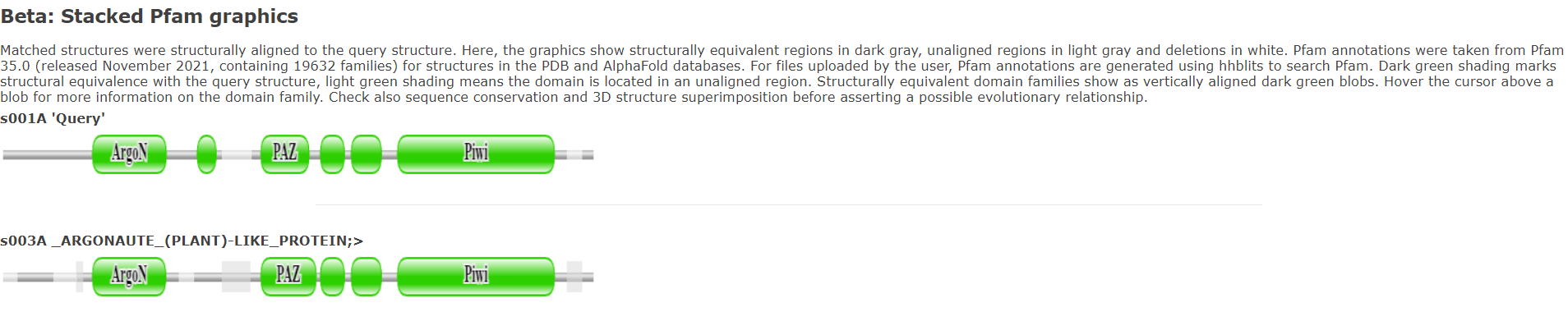

**A**

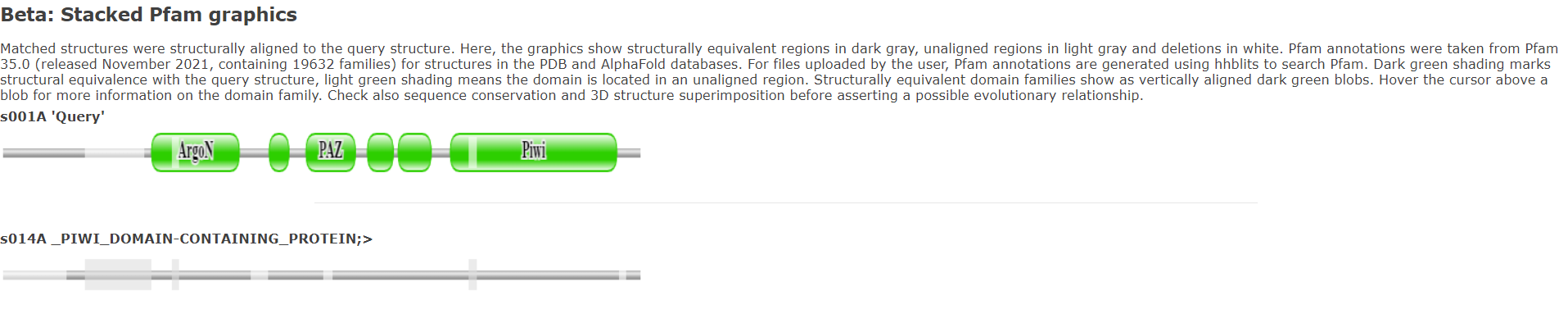

**B**

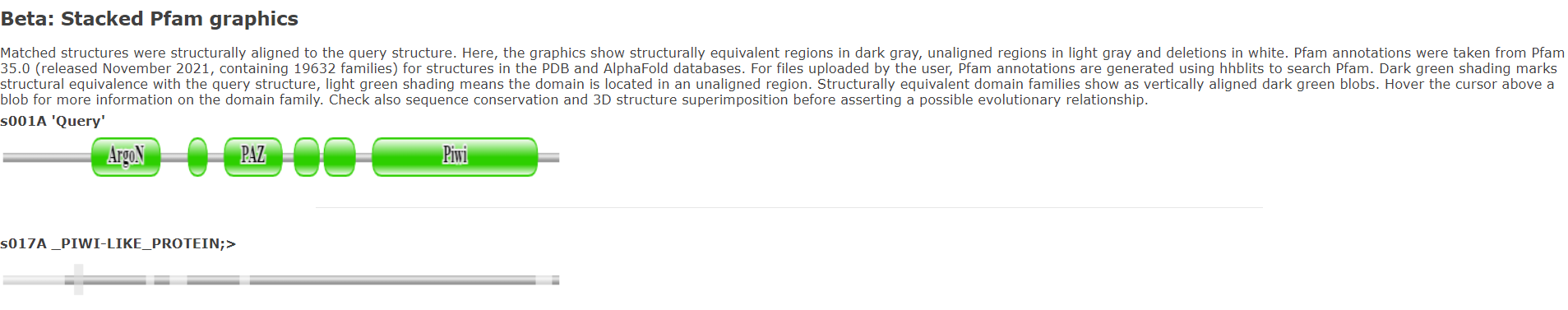

**C**

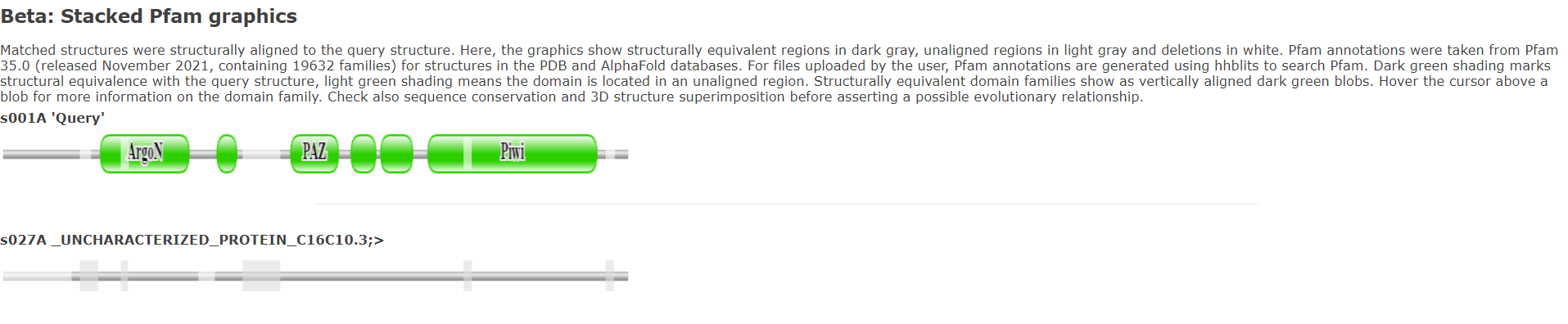

**D**

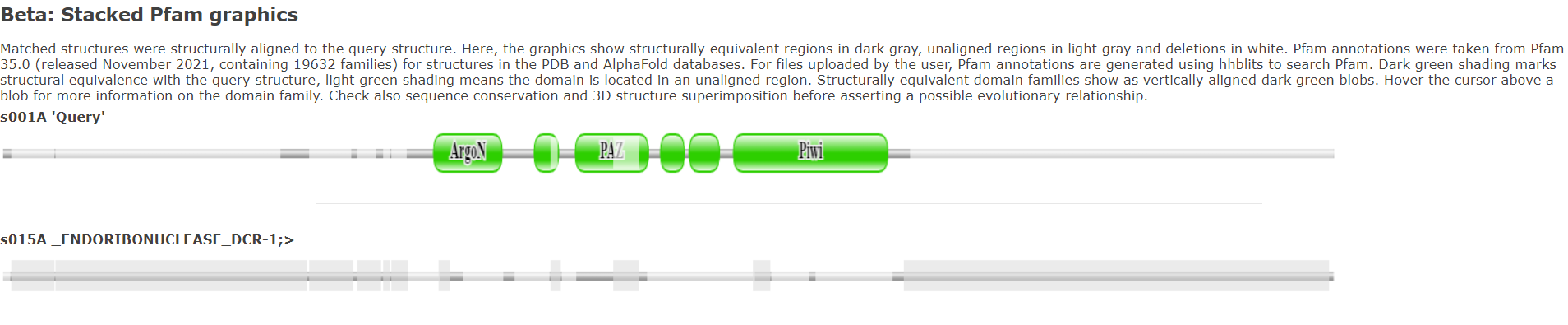

**E**

**Figure S2.** Example structural alignments of *C. elegans* ALG-1 (“Query”, top row) to other *C. elegans* Argonautes (bottom row). Alignments are shown for ALG-3 (A), CSR-1 (B), PRG-1 (C), and WAGO-9 (D). Regions of aligned structures are colored dark grey, with non-aligning structures being light grey and alignment gaps being white. Domains identified by DALI’s Pfam analysis are in green. From left to right, the domains shown for ALG-1 are the N domain, the Argonaute linker 1 domain, the PAZ domain, the Argonaute linker 2 domain, the MID domain, and the PIWI domain. As a control, we also show the alignment for *C. elegans* DCR-1 (E), which has a PAZ domain but no other Argonaute domains. As expected, the PAZ domain is generally the only structure that aligns between ALG-1 and DCR-1.

**B**

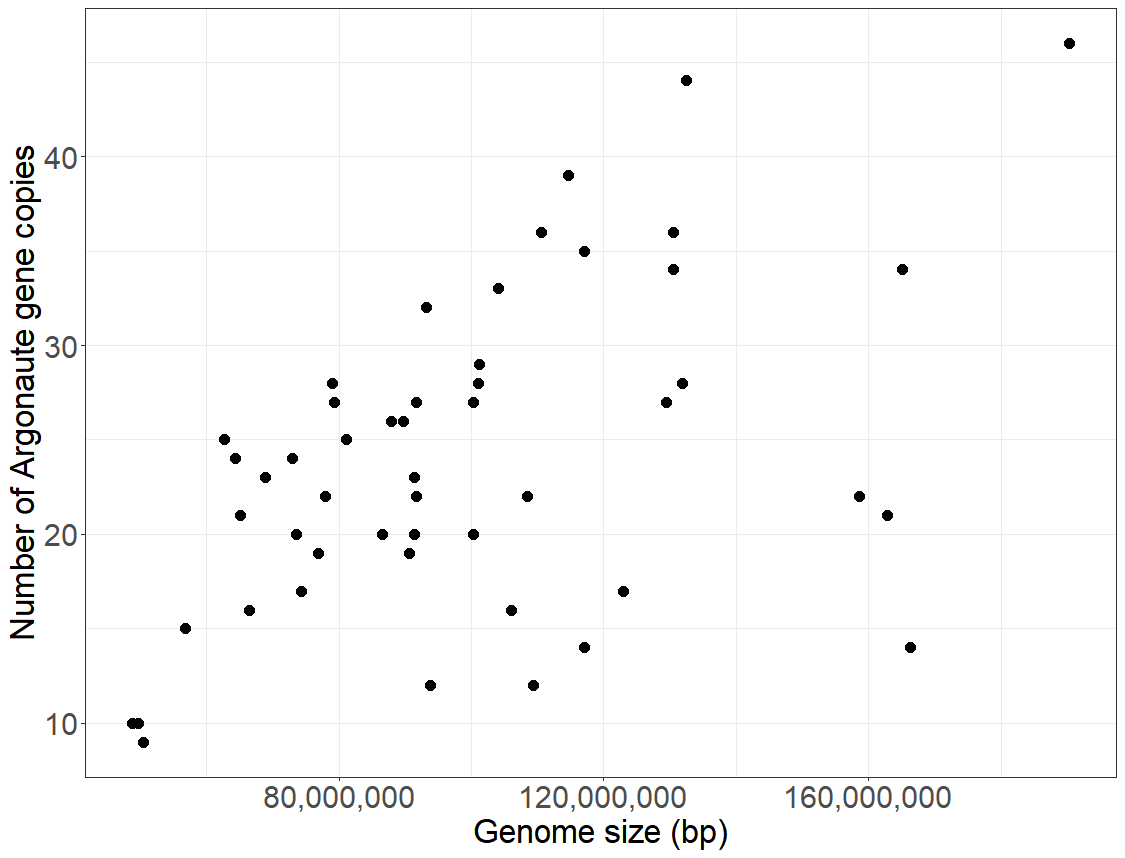

**A**

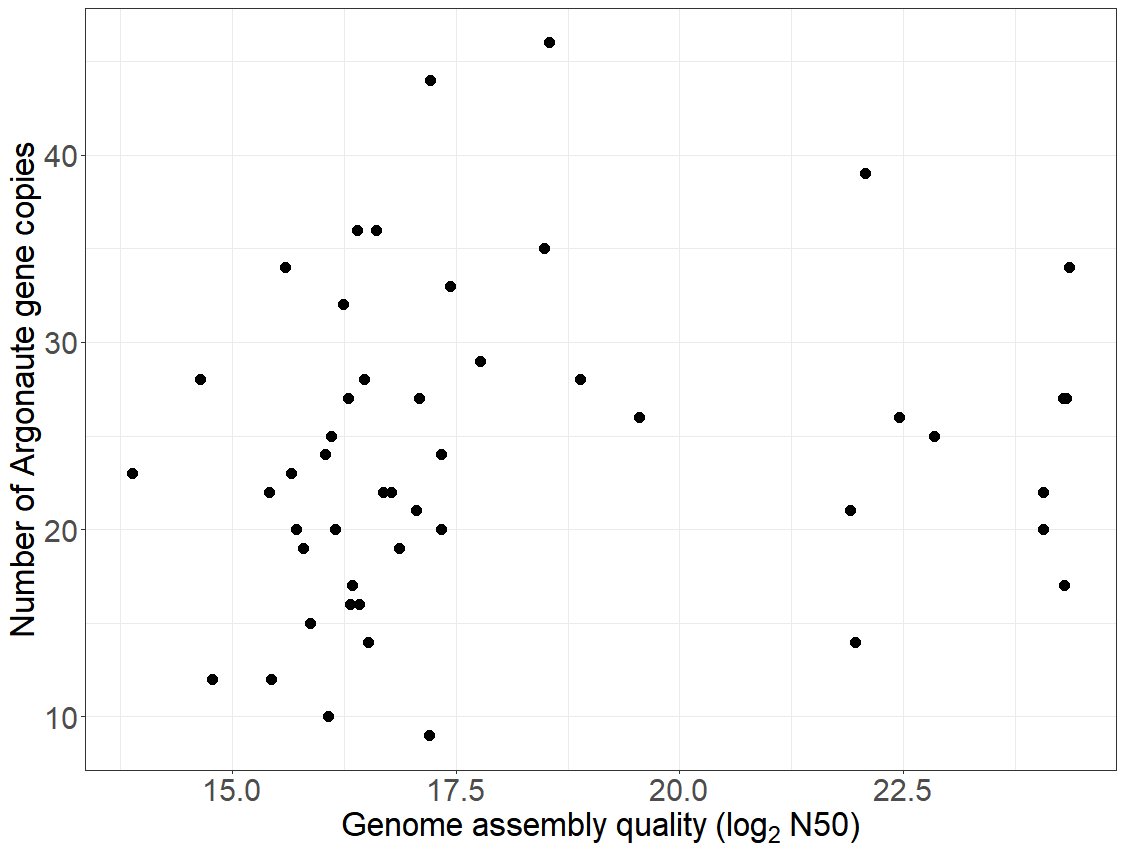

**C**

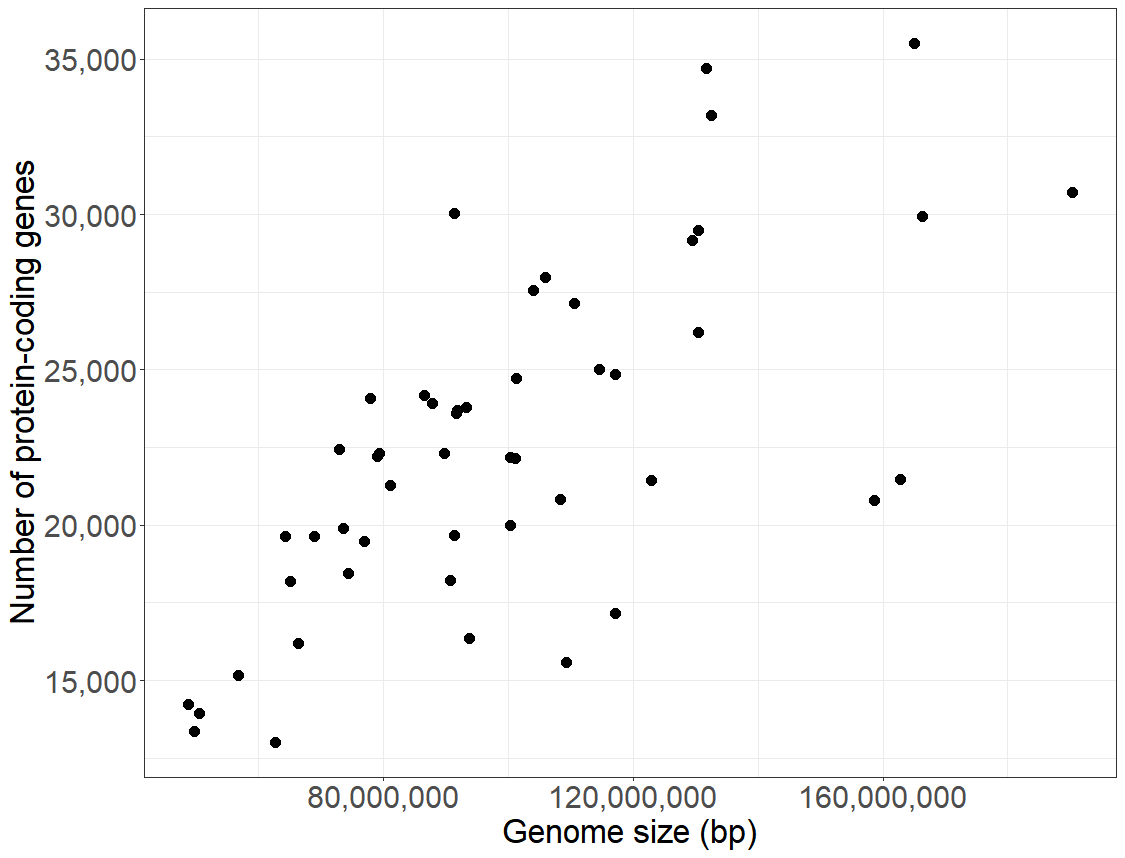

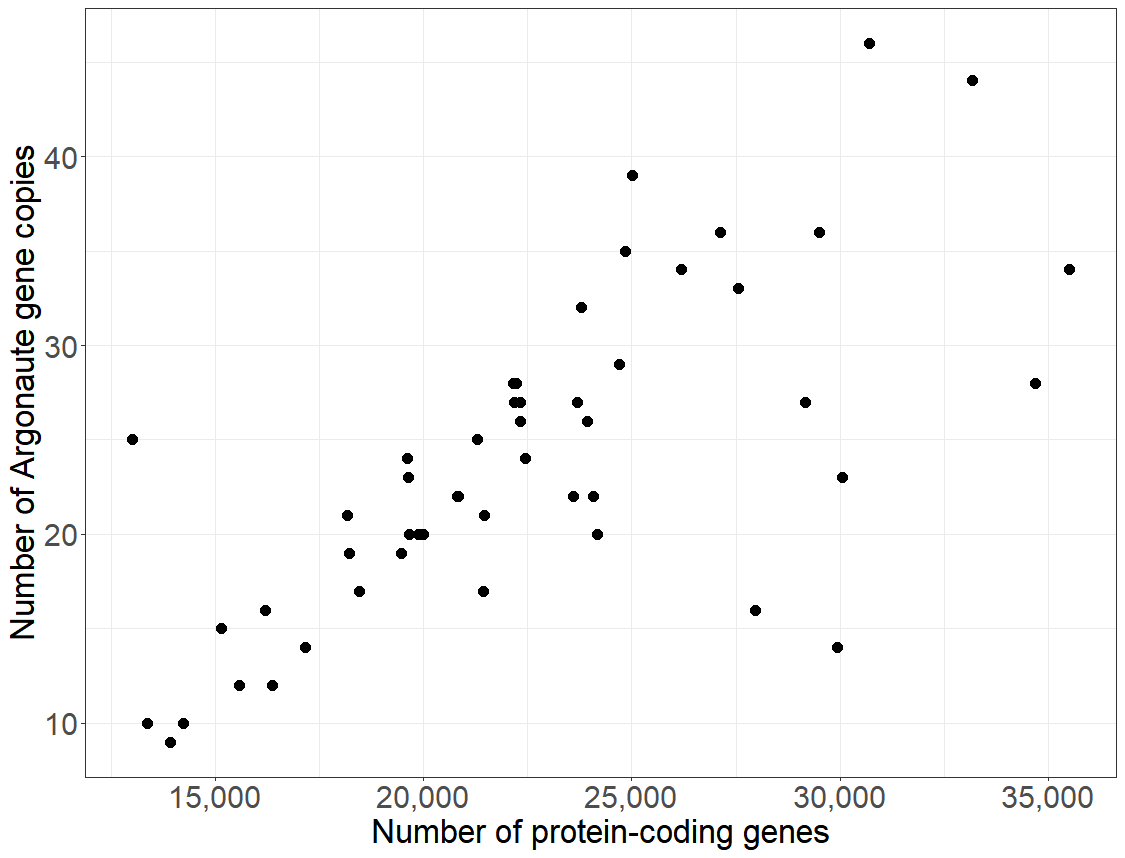

**D**

***P* = 0.0034**

**R^2^ = 0.15**

***P* = 0.33**

**R^2^ = -0.00069**

***P* = 1.6x10^-6^**

**R^2^ = 0.37**

***P* = 0.0010**

**R^2^ = 0.19**

**Figure S3.** Relationship between total number of Argonaute genes and genome size (A), between total number of Argonaute genes and genome assembly N50 (B), between overall number of protein-coding genes and genome size (C), and between total number of Argonaute genes and overall number of protein-coding genes (D) for the 50 species with available genome assemblies.

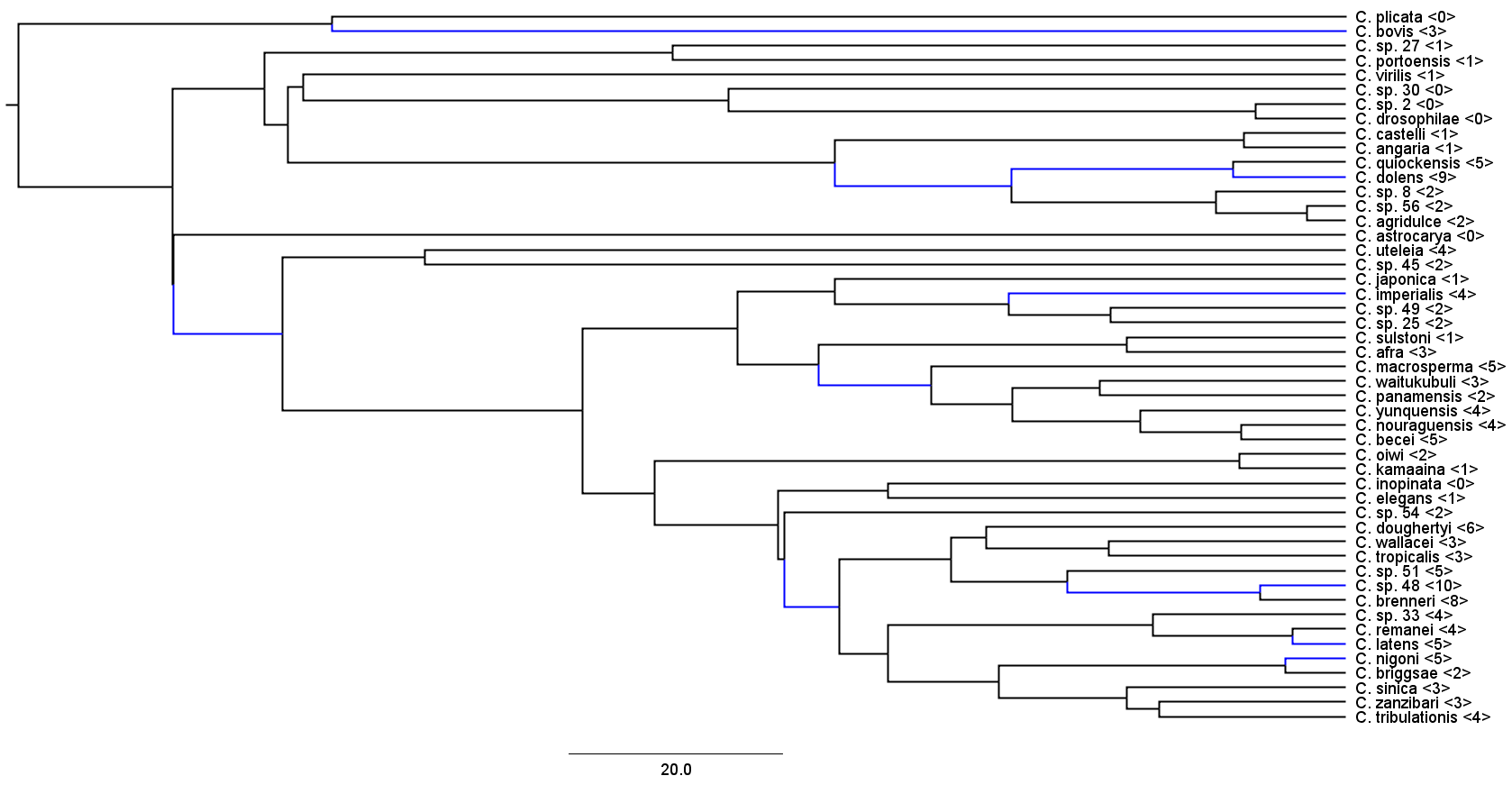

**A**

200

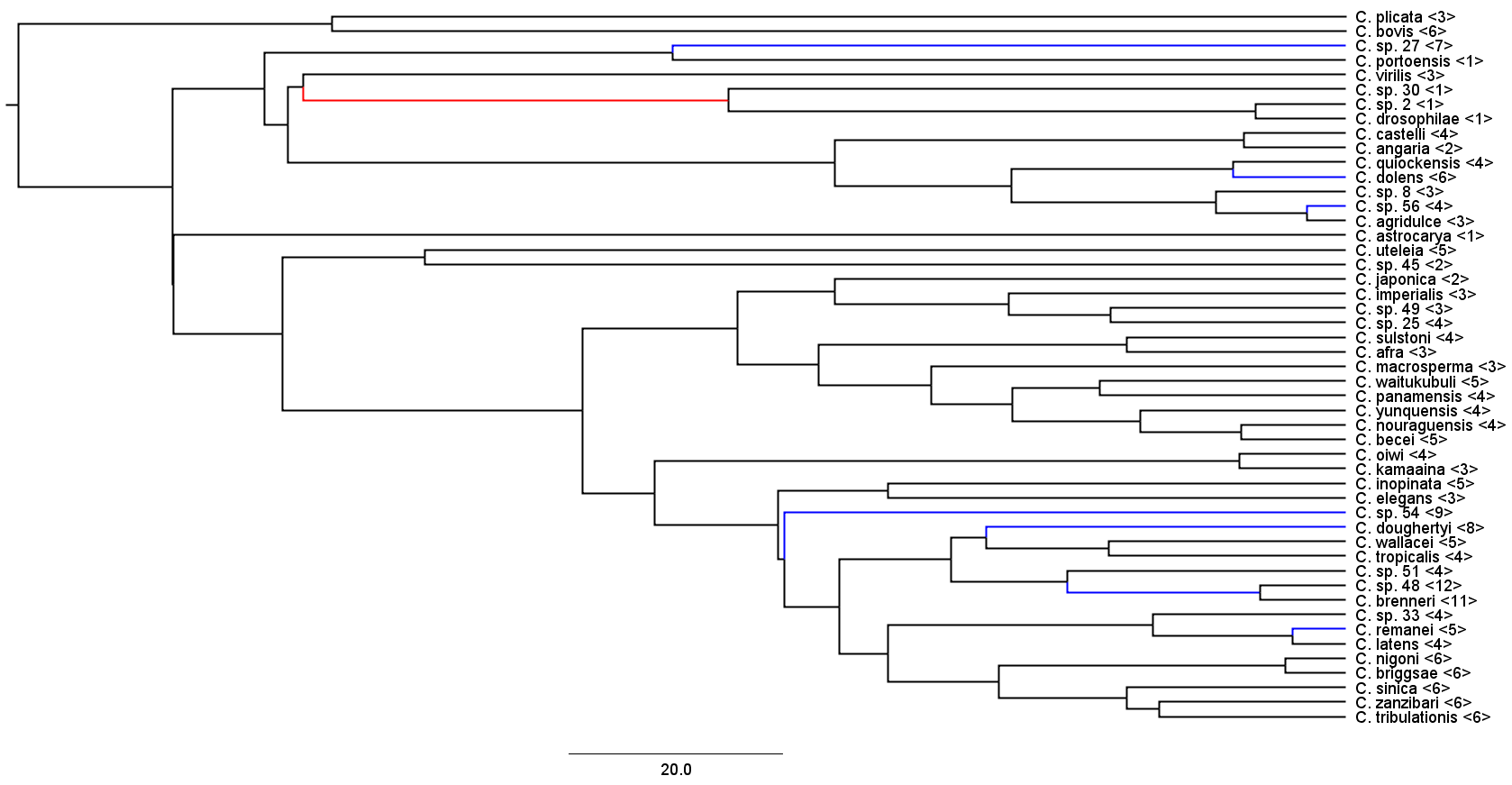

**B**

200

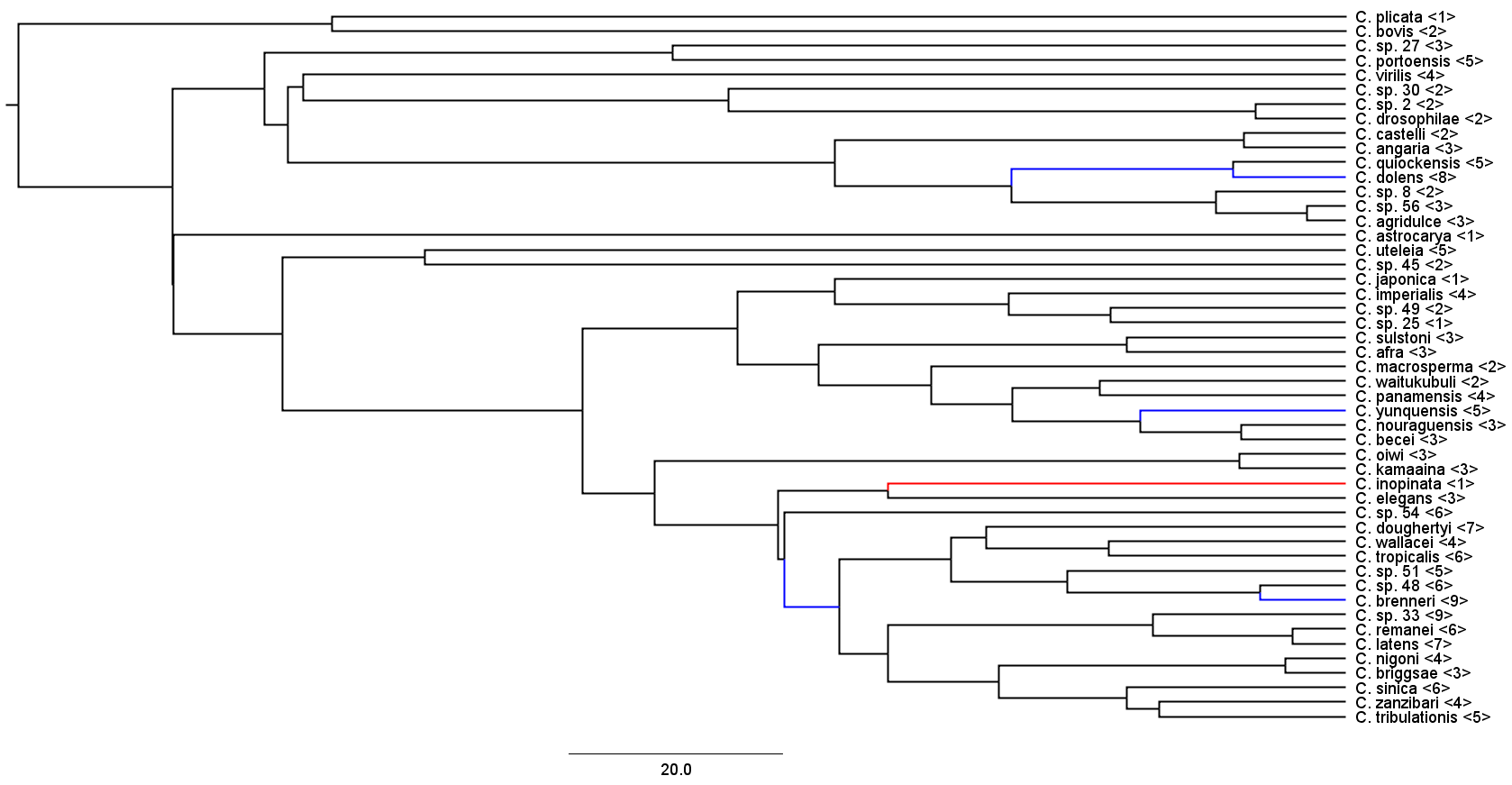

**C**

200

**Figure S4.** Branches on the species tree that have experienced significant gene copy number expansions (blue) or contractions (red) of the ERGO-1 (A), WAGO-6/7/8 (B), and WAGO-9/10/12 (C) subfamilies according to CAFE. Numbers in brackets indicate the number of genes belonging to that subfamily in each species. Branch lengths are in units of millions of generations.

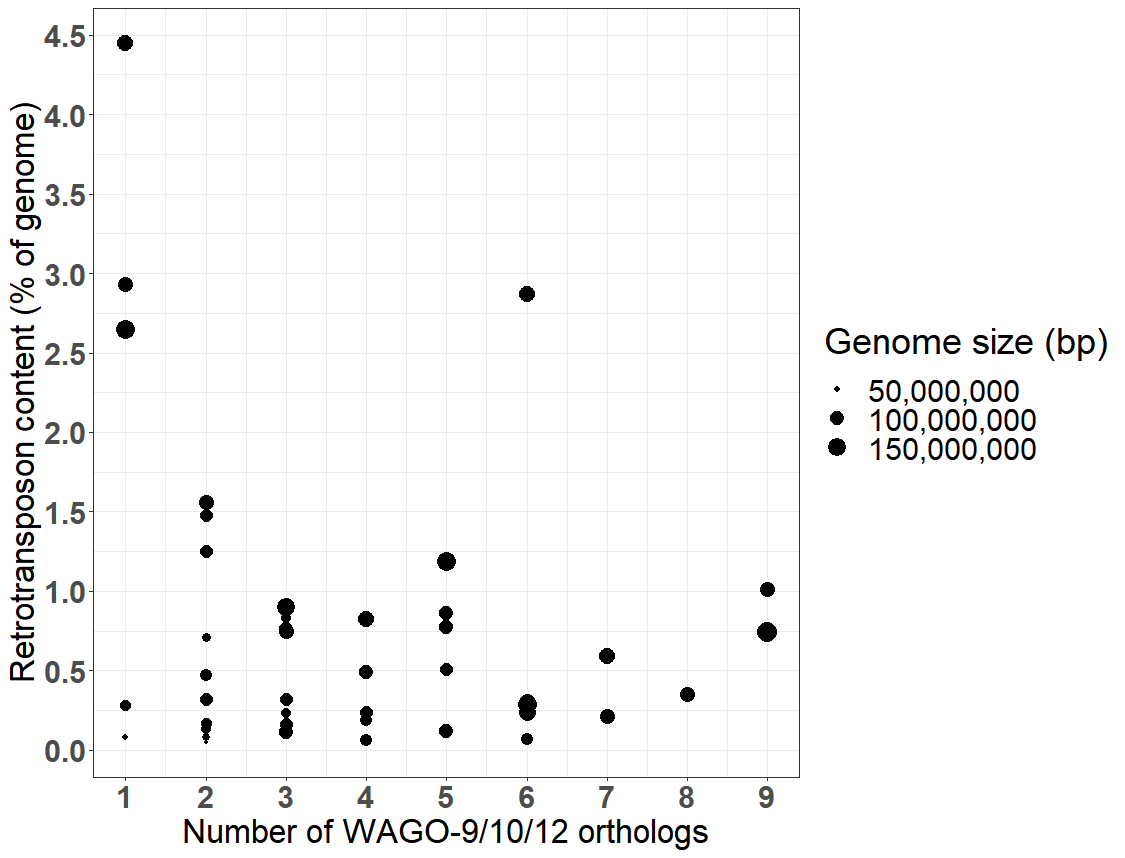

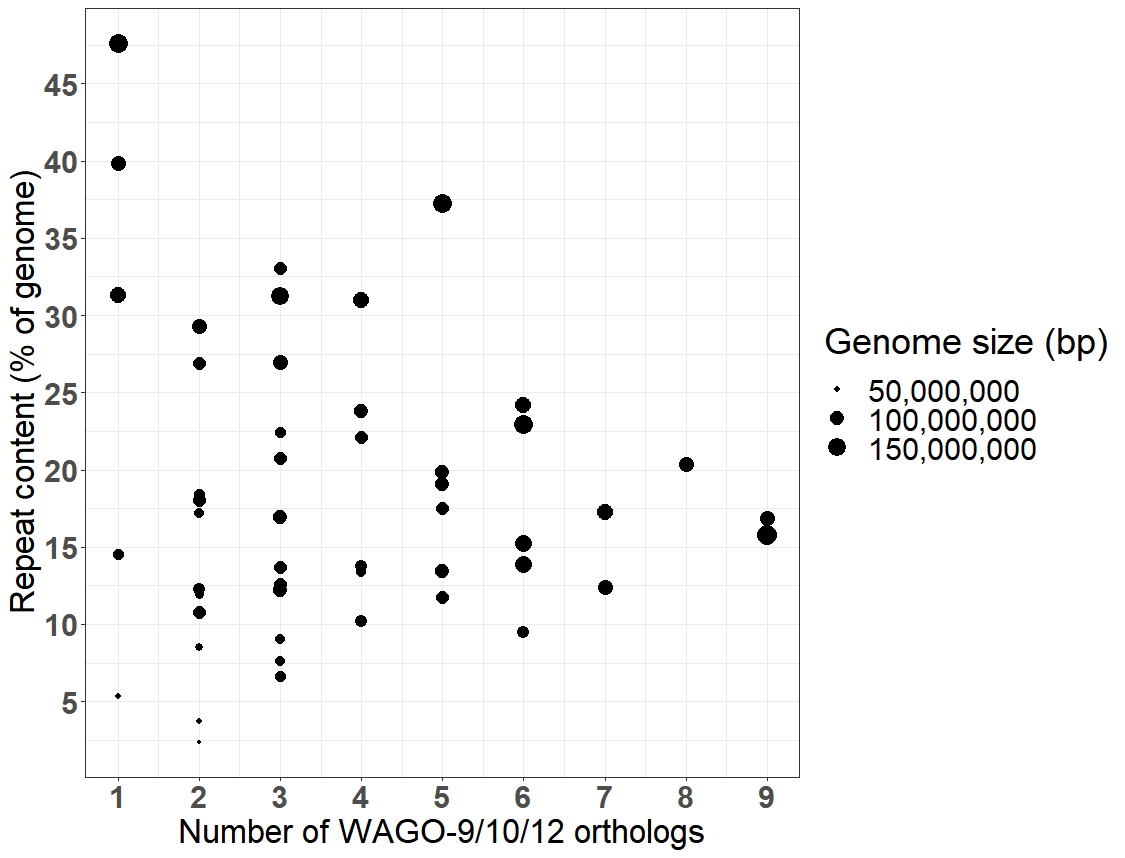

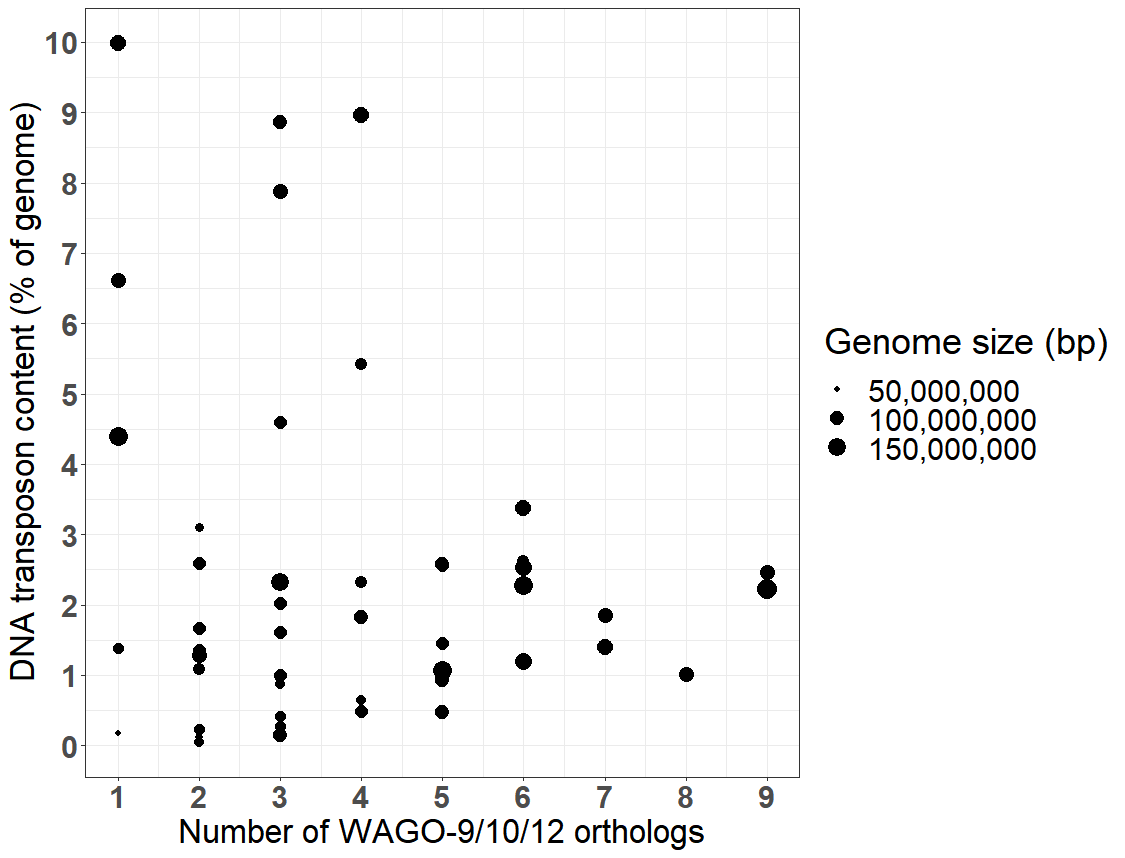

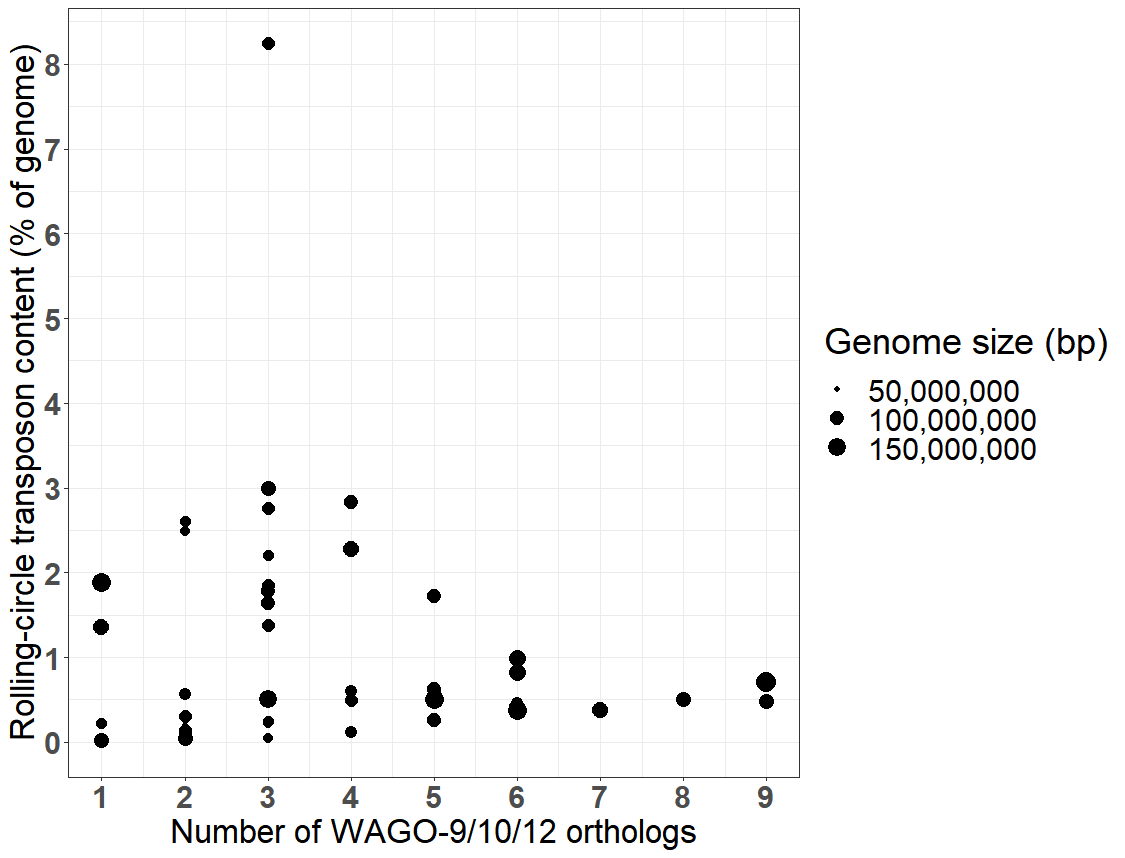

**A**

**B**

**C**

**D**

**Figure S5.** The number of WAGO-9/10/12 orthologs compared to overall repeat content (A), retrotransposon content (B), DNA transposon content (C), and rolling-circle transposon content (D) for the 50 species with available genome assemblies.

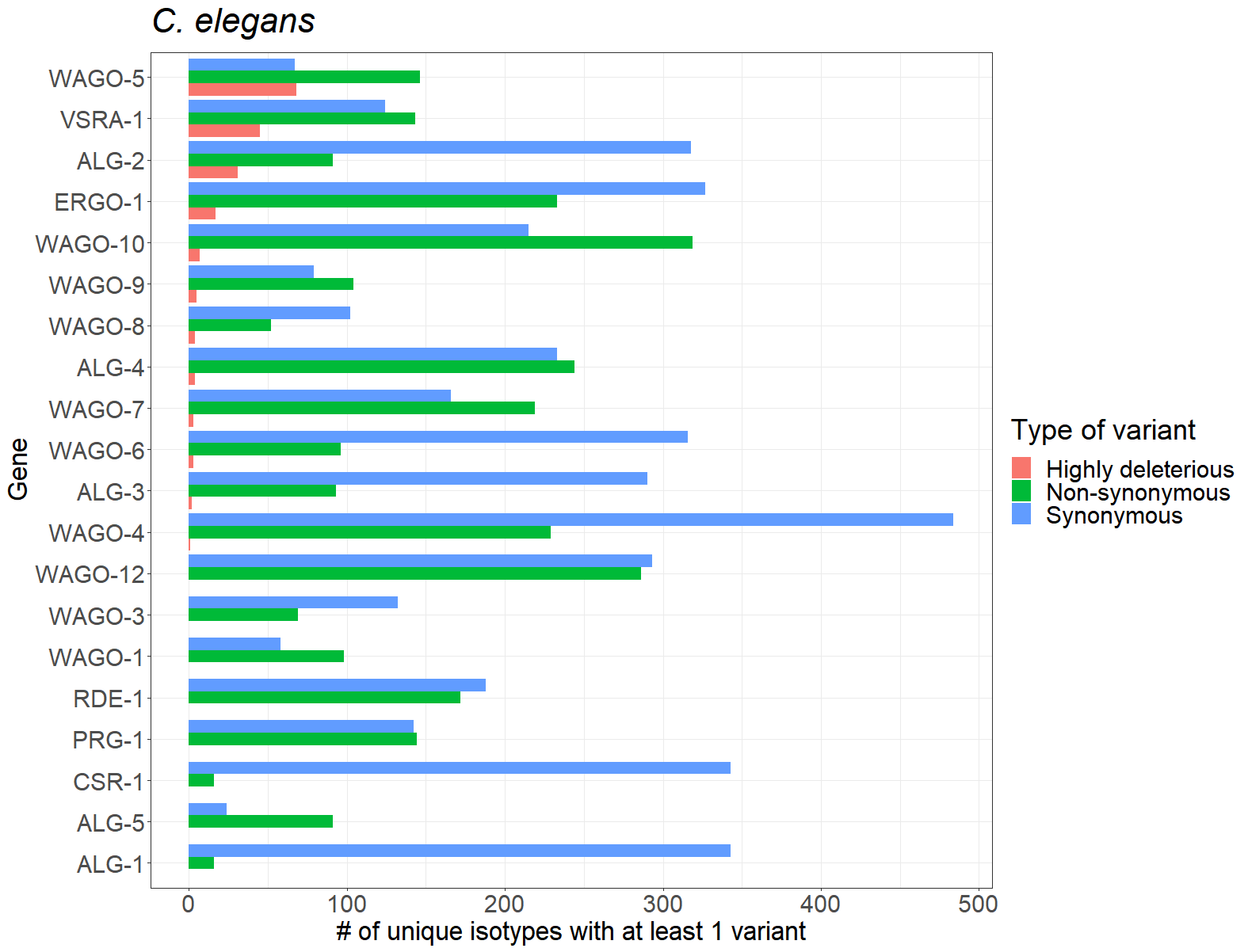

**A**

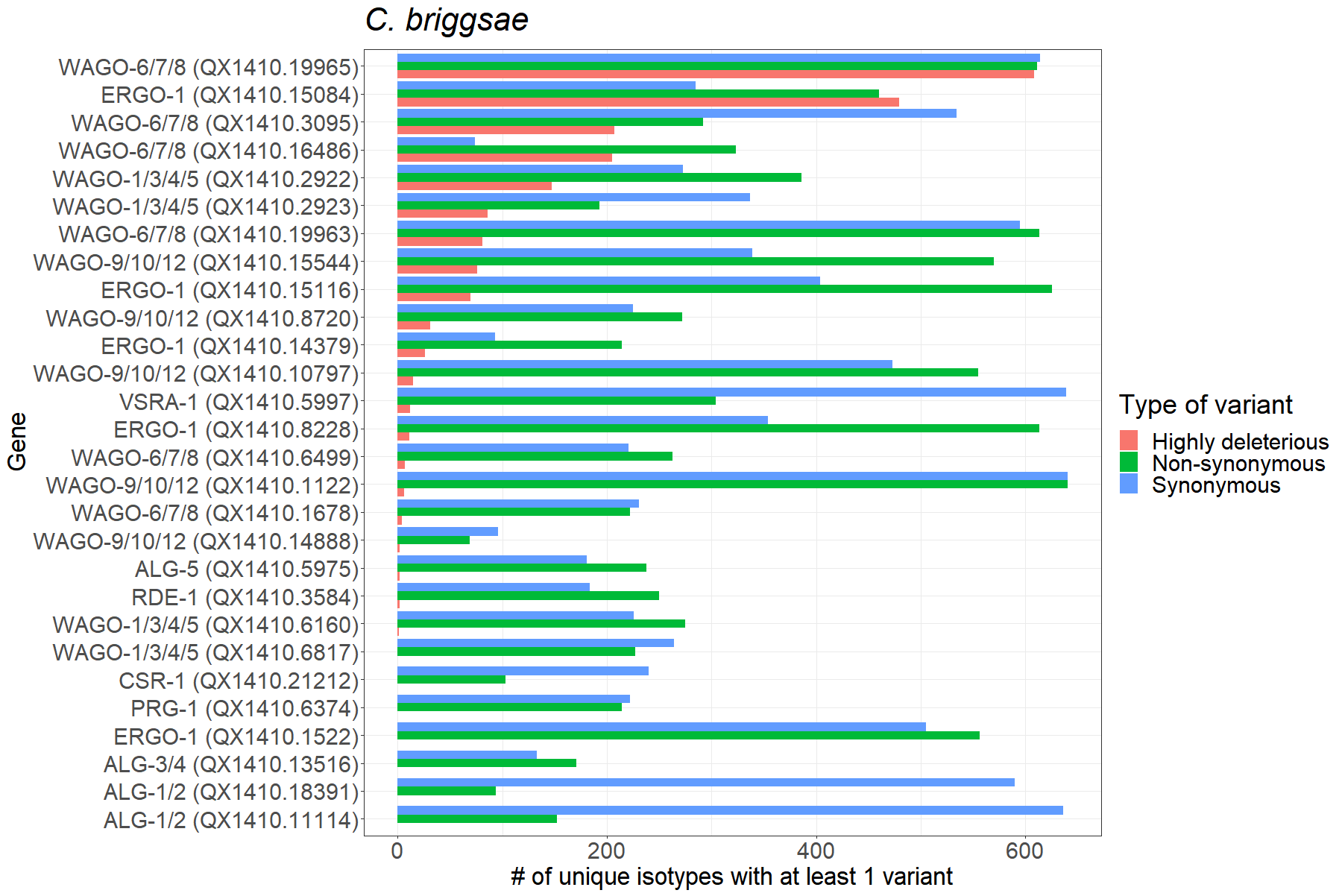

**B**

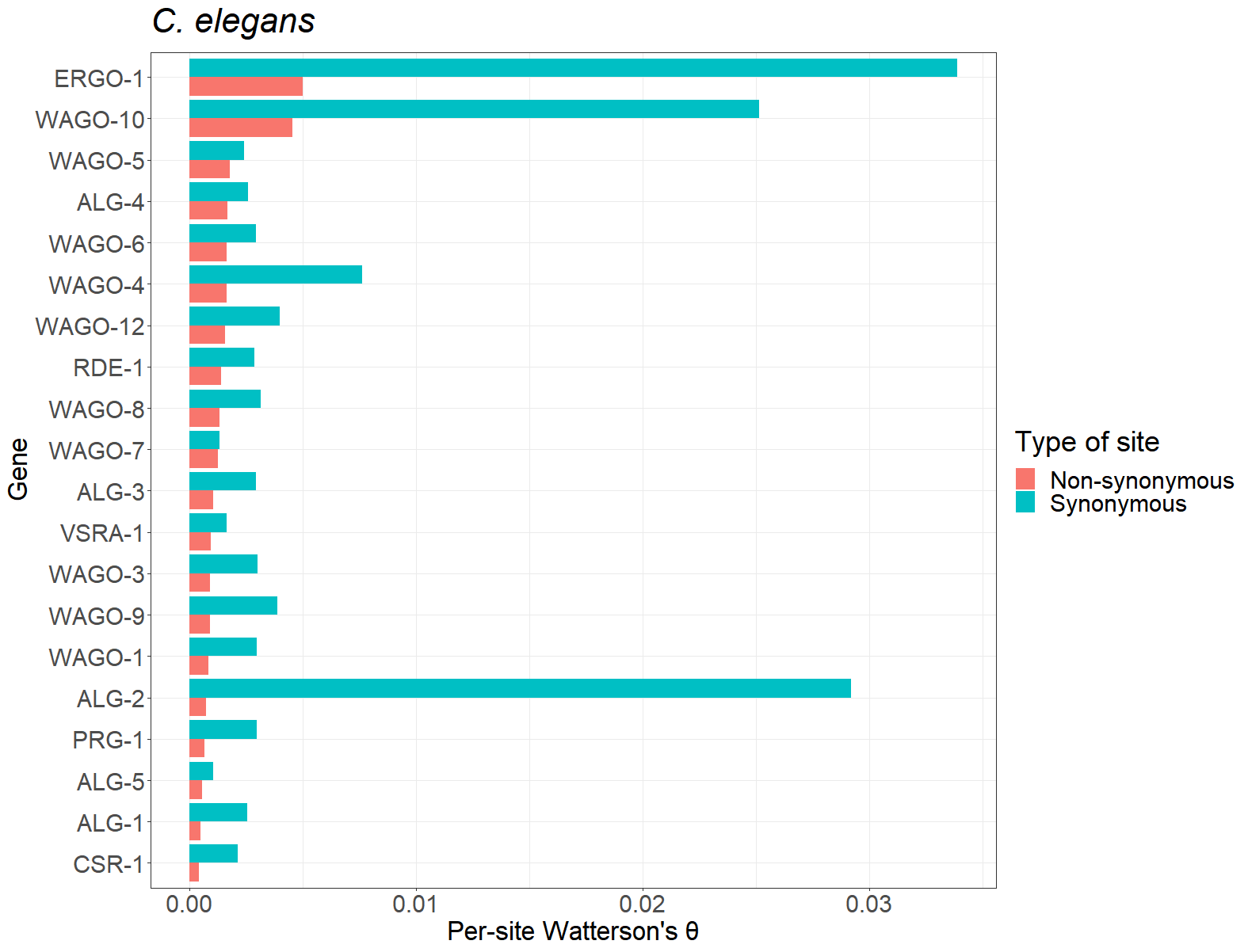

**C**

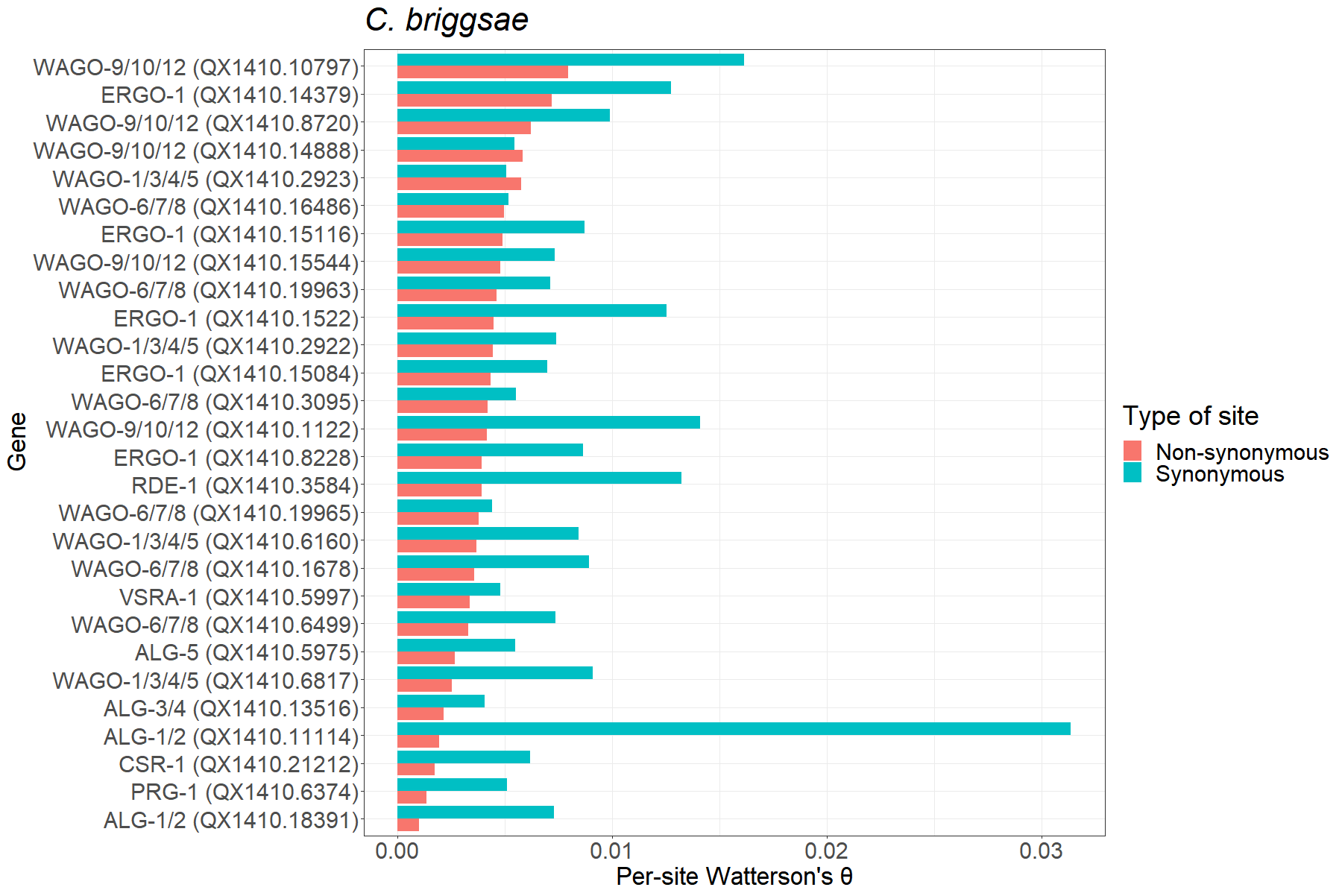

**D**

**Figure S6.** (A, B) The number of unique isotypes that carry at least 1 variant of a given kind, for 3 different kinds of variants, for Argonaute genes in *C. elegans* (A) and *C. briggsae* (B).

(C, D) Per-site Watterson’s θ at synonymous and non-synonymous sites for Argonaute genes in *C. elegans* (C) and *C. briggsae* (D).

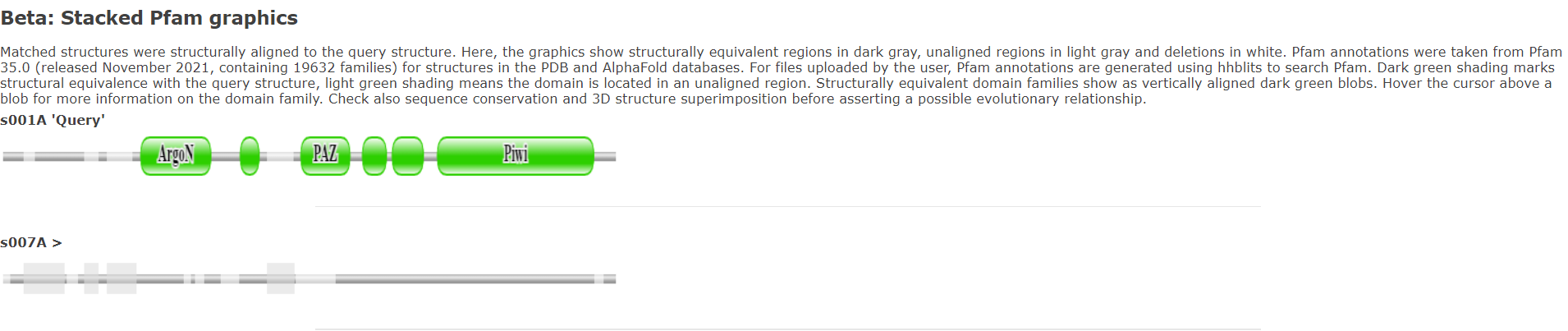

**A**

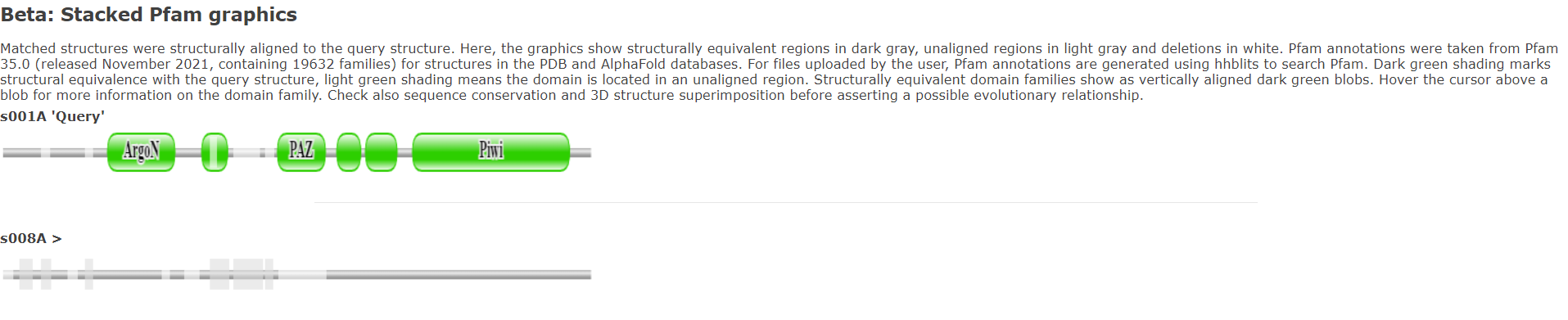

**B**

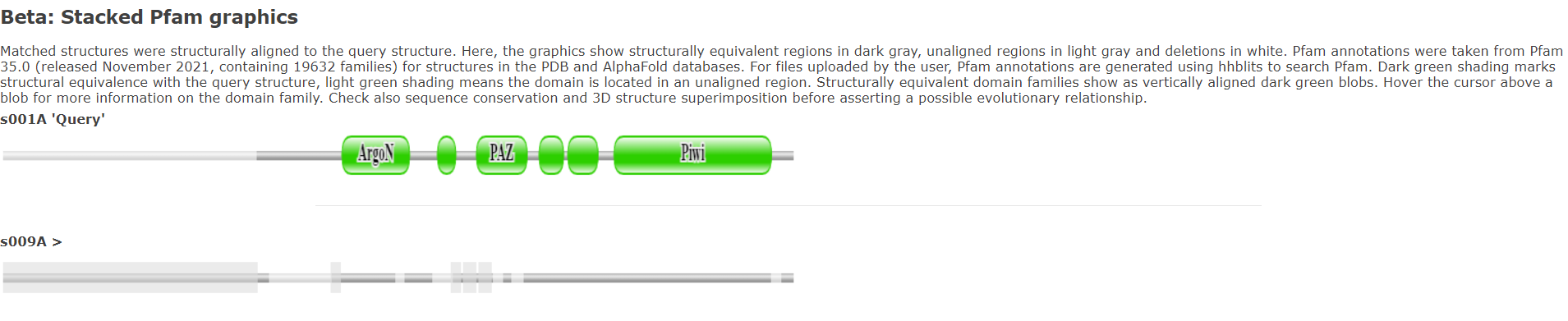

**C**

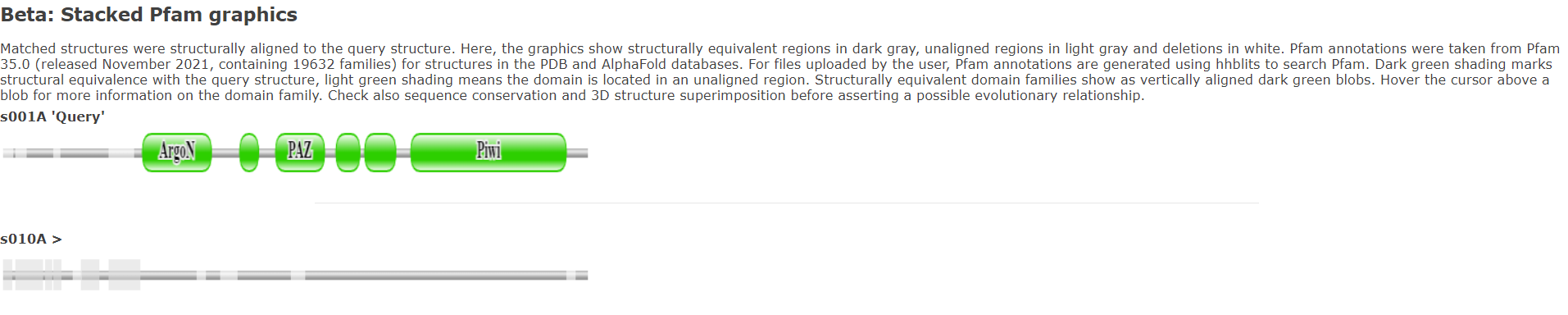

**D**

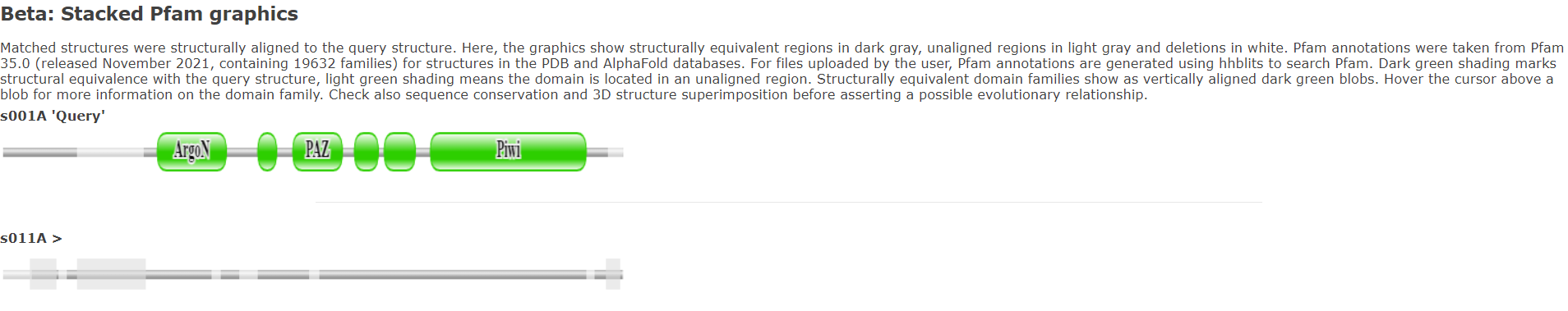

**E**

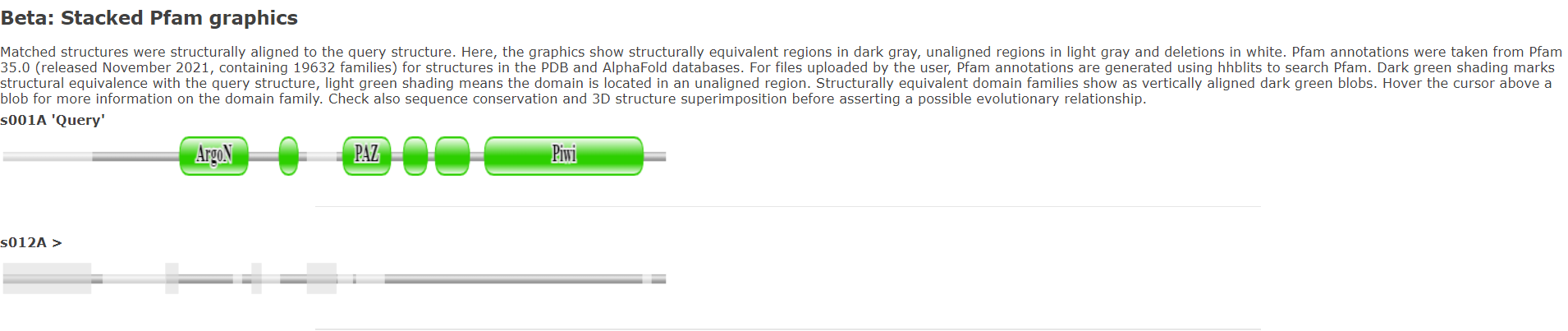

**F**

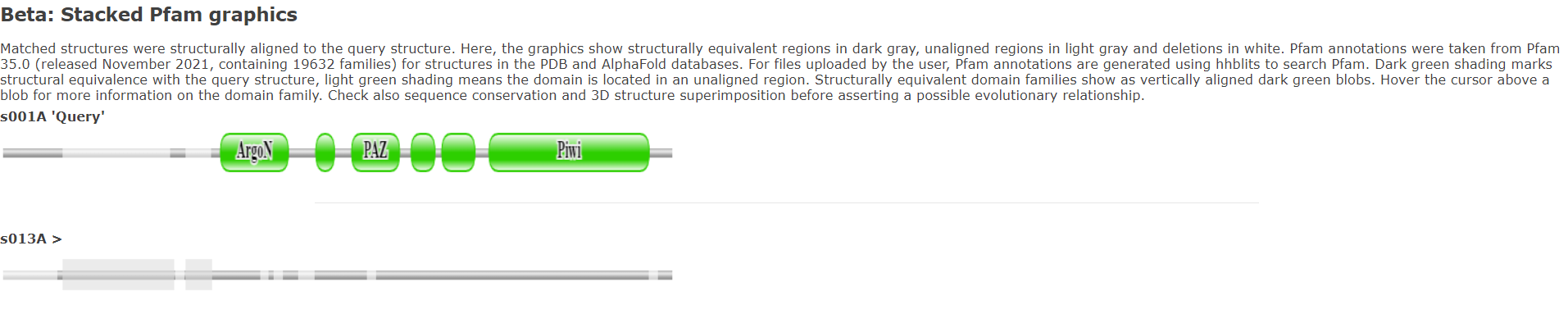

**G**

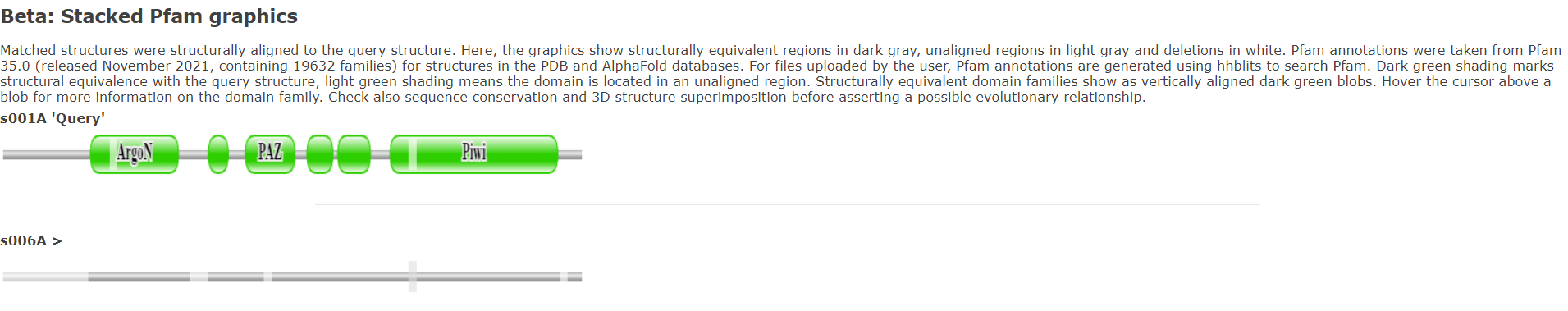

**H**

**Figure S7.** Structural alignments of *C. elegans* ALG-1 (“Query”, top row) to the 7 divergent *C. panamensis* Argonautes (bottom row). The divergent *C. panamensis* Argonautes are CSP28.g8935.t1 (A), CSP28.g16020.t1 (B), CSP28.g16021.t1 (C), CSP28.g16024.t1 (D), CSP28.g16025.t1 (E), CSP28.g16026.t1 (F), and CSP28.g16027.t1 (G). Regions of aligned structures are colored dark grey, with non-aligning regions being light grey and alignment gaps being white. Domains identified by DALI’s Pfam analysis are in green. From left to right, the domains shown for ALG-1 are the N domain, the Argonaute linker 1 domain, the PAZ domain, the Argonaute linker 2 domain, the MID domain, and the PIWI domain. The PAZ domain appears to be mostly or entirely absent in CSP28.g8935.t1, CSP28.g16020.t1, CSP28.g16021.t1, and CSP28.g16026.t1. As a control, *C. panamensis* CSR-1 is also shown (H).

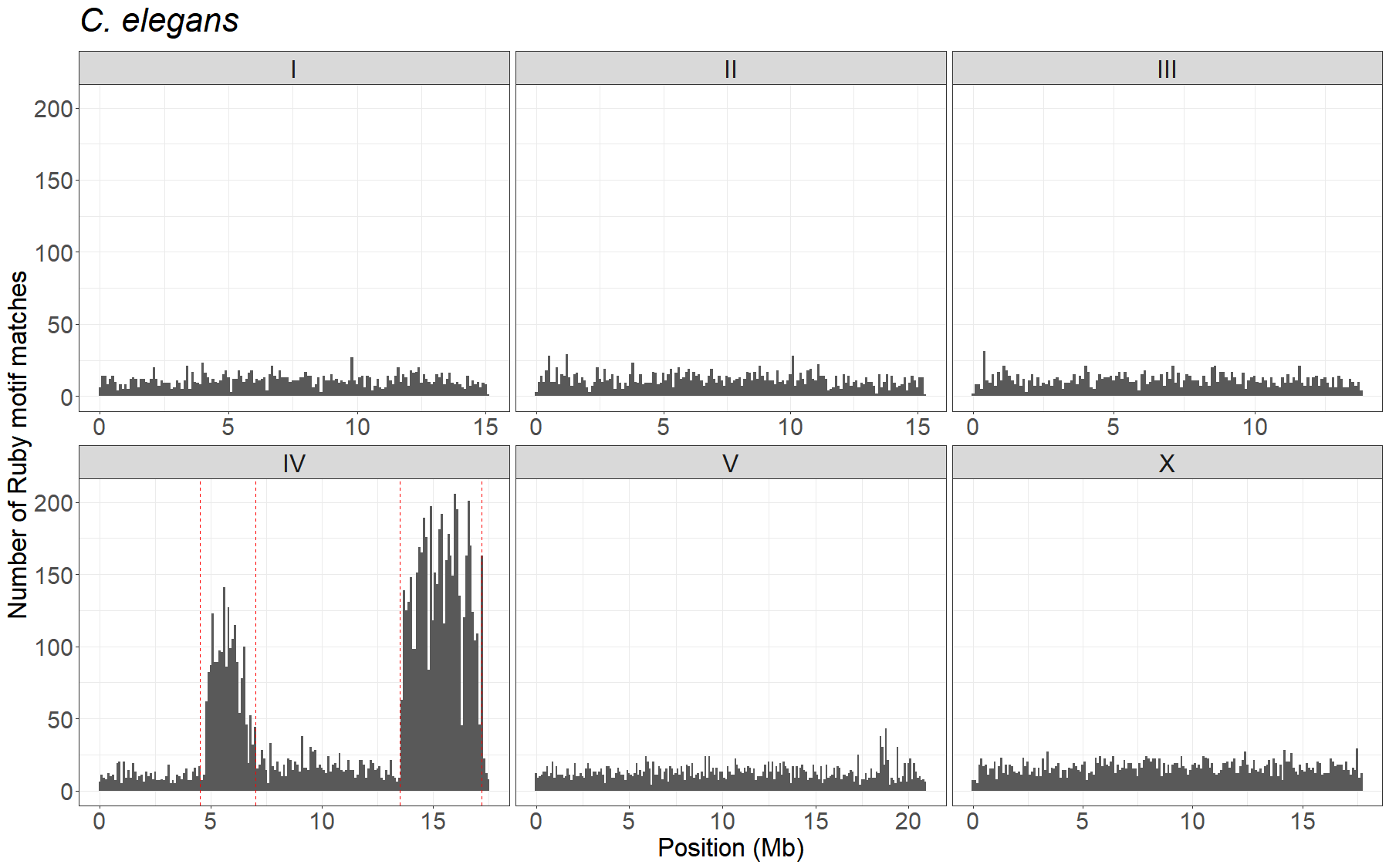

**A**

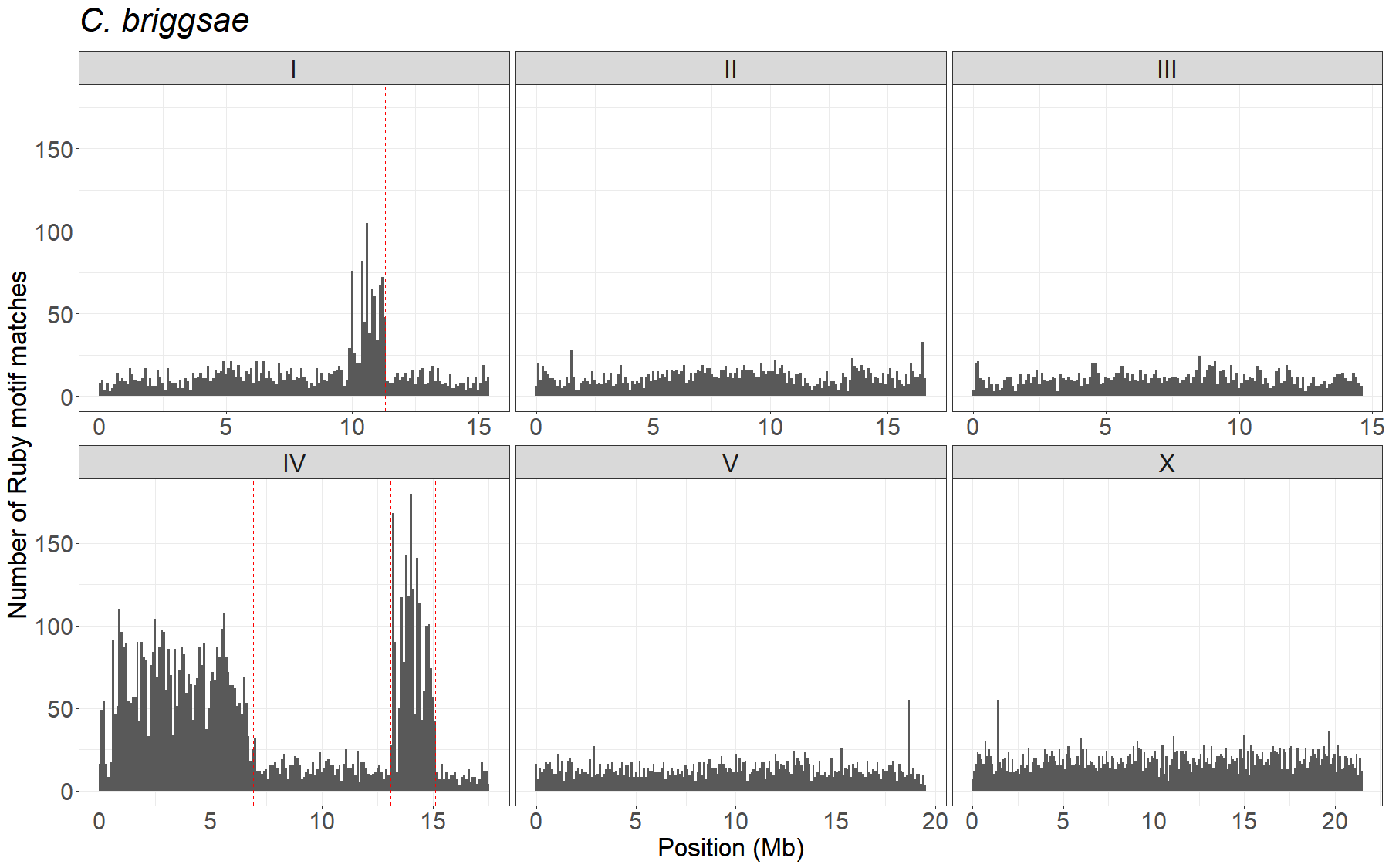

**B**

**Figure S8.** Number of Ruby motifs (CNGTTTCA) in noncoding regions detected in non-overlapping 100kb bins along the genomes of *C. elegans* (A) and *C. briggsae* (B). Dashed vertical lines indicate the boundaries of known piRNA clusters in these species.

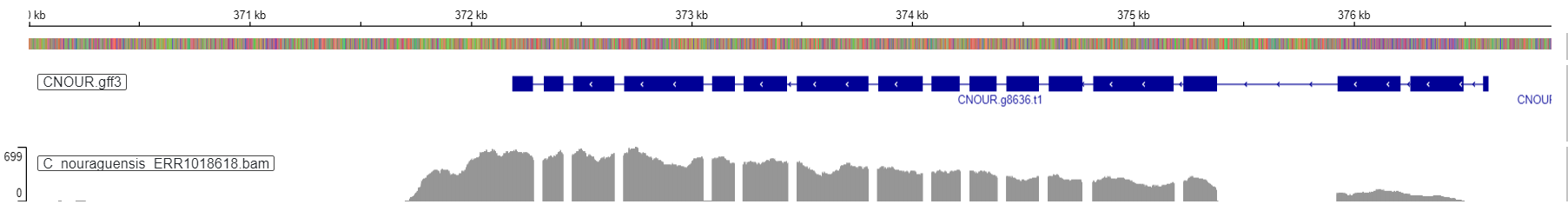

**A**

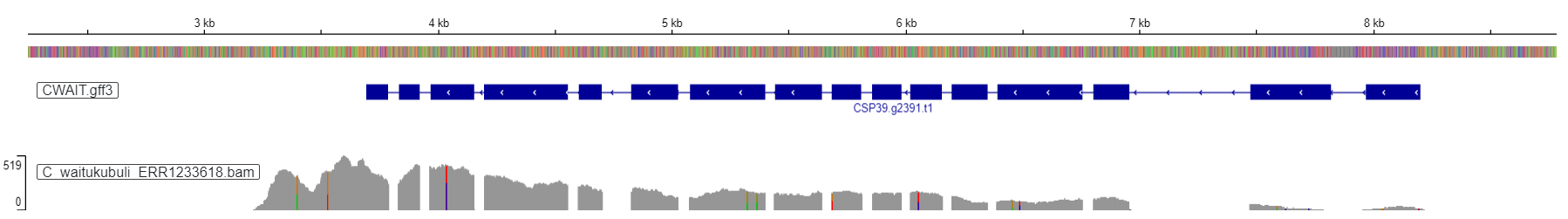

**B**

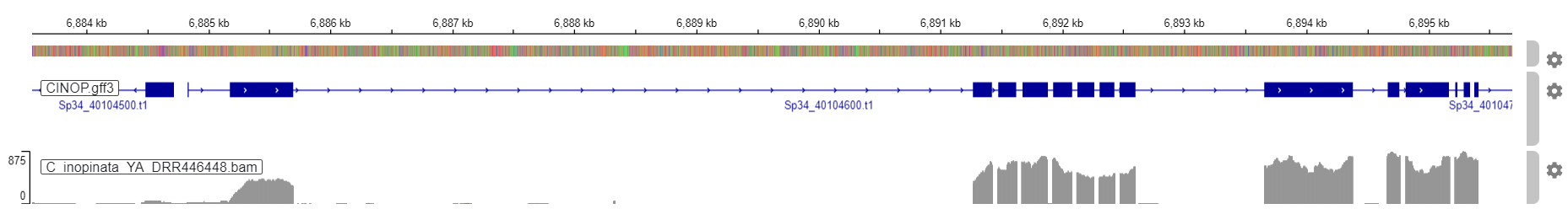

**C**

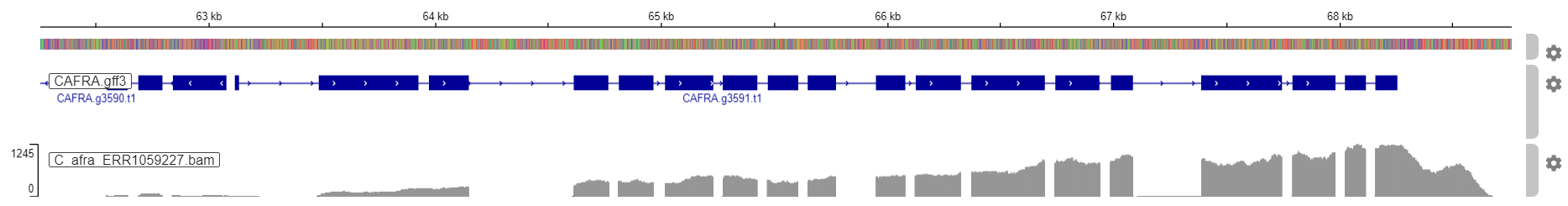

**D**

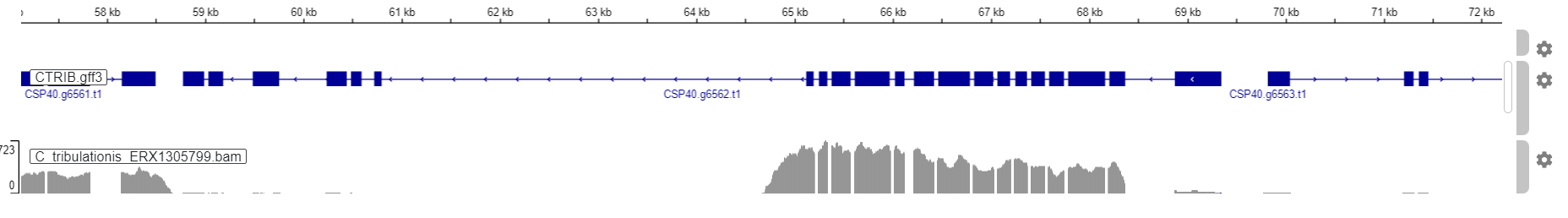

**E**

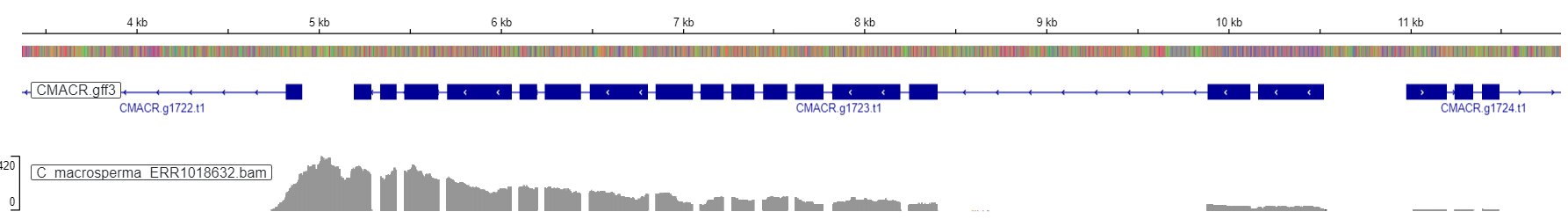

**F**

**Figure S9.** Genome browser tracks showing the number of aligned RNA-seq reads (grey histograms) along the CSR-1 locus in *C. nouraguensis* (A, CNOUR.g8636.t1)*, C. waitukubuli* (B, CSP39.g2391.t1), *C. inopinata* (C, Sp34_40104600.t1)*, C. afra* (D, CAFRA.g3591.t1)*, C. tribulationis* (E, CSP40.g6562.t1)*,* and *C. macrosperma* (F, CMACR.g1723.t1). Red boxes indicate the putative CSR-1a exon(s).

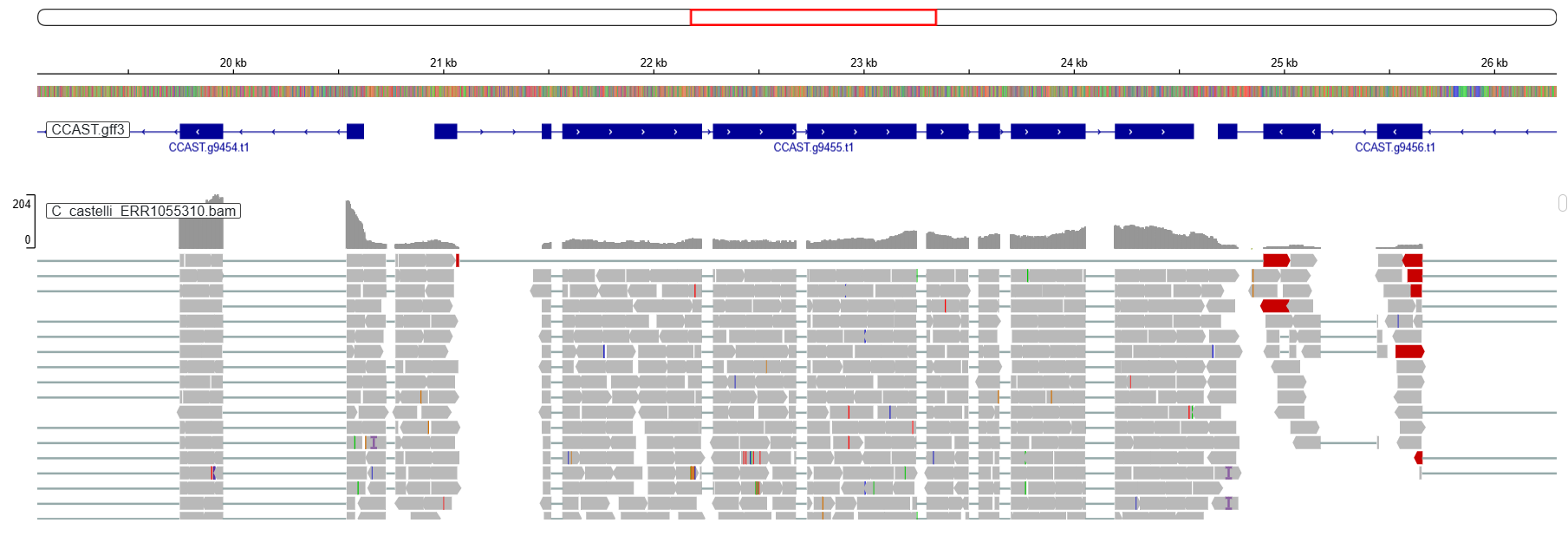

**A**

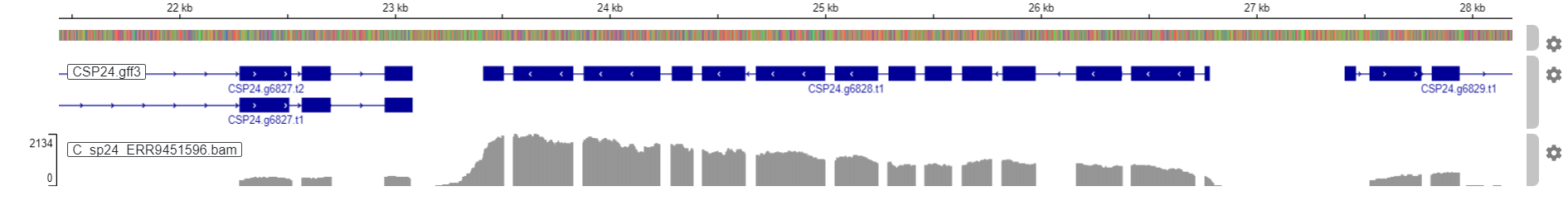

**B**

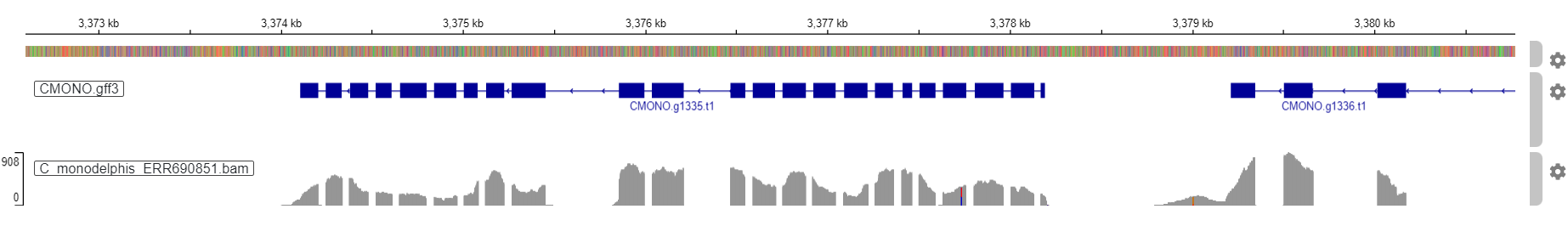

**C**

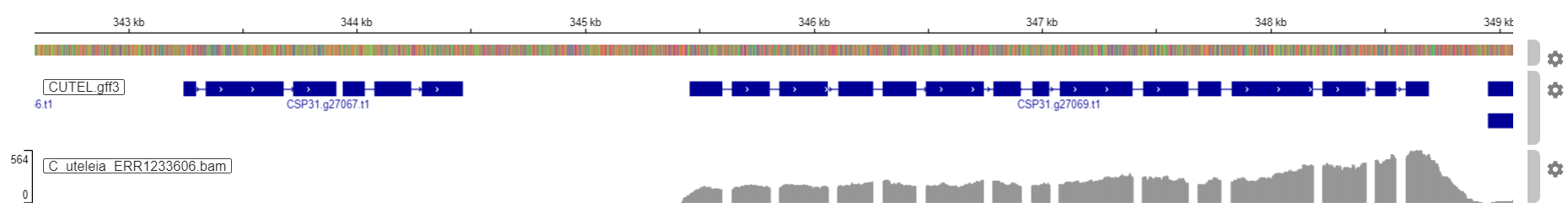

**D**

**Figure S10.** Genome browser tracks showing the number of aligned RNA-seq reads (grey histograms) along the CSR-1 locus in *C. castelli* (A, CCAST.g9455.t1), *C. agridulce* (B, CSP24.g6828.t1), *C. monodelphis* (C, CMONO.g1335.t1), and *C. uteleia* (D, CSP31.g27069.t1). Upstream regions outlined in red are the expected locations of a theoretical unannotated CSR-1a exon, if such an exon existed. The true start codon for *C. castelli* (A) appears to be upstream of the currently-annotated start codon, though this first exon does not contain any RG repeats, and so is likely not the CSR-1a exon.

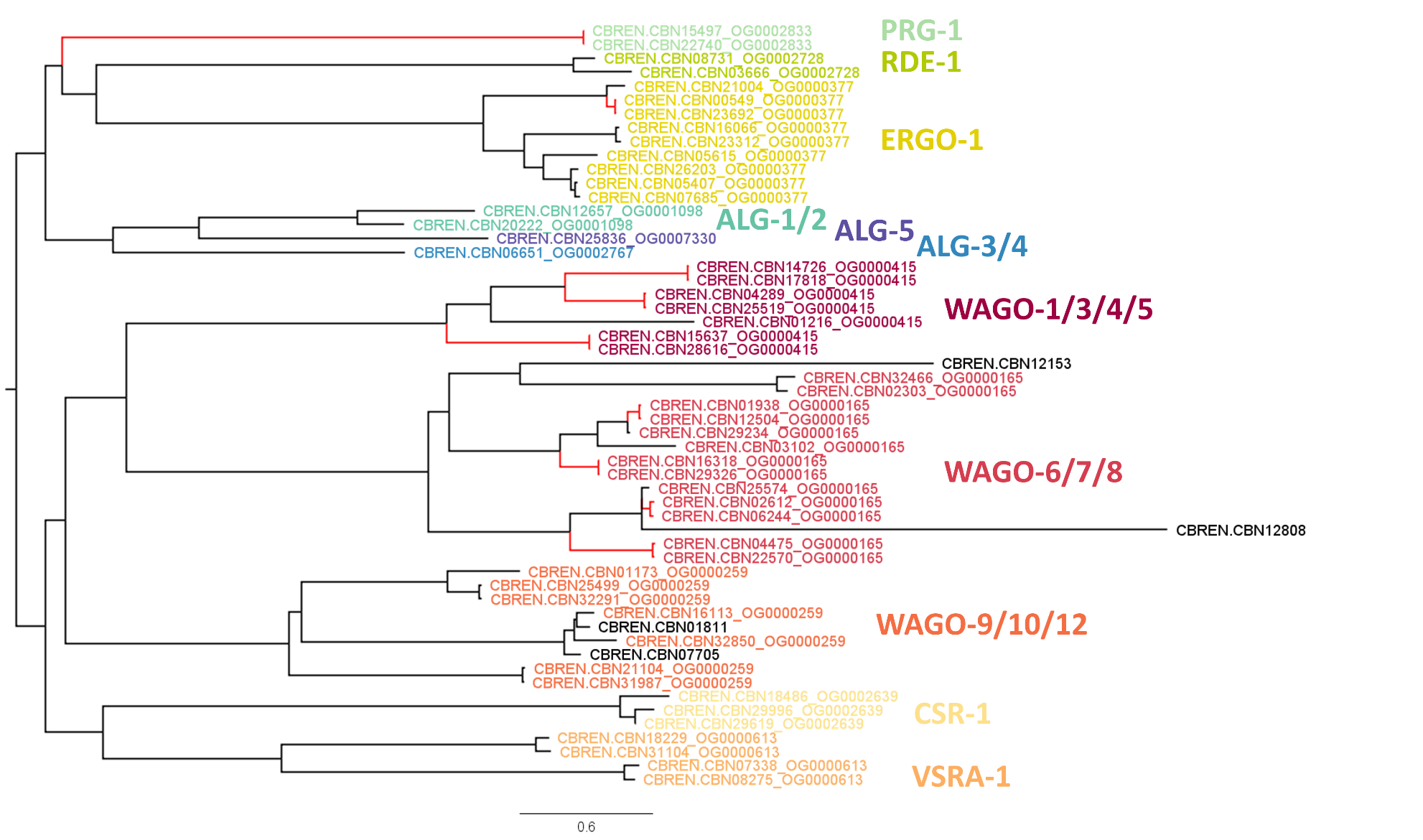

**Figure S11.** Subset of the full gene tree of 1213 Argonaute protein sequences, showing only the Argonaute genes in *C. brenneri*. Red branches indicate the 9 pairs of Argonautes that were each considered to be allelic variants of the same gene (i.e., pairs with protein sequence identity ≥ 95% and K_S_ < 0.141). Branch lengths are in units of amino acid substitutions per site.

Supplementary Methods

Identification of Argonaute genes

Reference genome assemblies for 50 *Caenorhabditis* species, as well as a reference transcriptome for 1 species (*C.* sp. 45), were obtained from a combination of NCBI GenBank [1], WormBase ParaSite [2, 3], and the Caenorhabditis Genomes Project [4] (Table S3). We assessed the quality of all 51 genome and transcriptome assemblies by running BUSCO v5.2.2 [5] to detect the single-copy genes in the nematoda_odb10 gene set (Table S2).

We defined groups of orthologous genes (orthogroups) between all 51 *Caenorhabditis* species with OrthoFinder v2.5.2 [6]. We provided OrthoFinder with the protein sequences of the longest isoform of every annotated gene for each species, as well as a phylogeny of all 51 species (see “Gene family evolution” for details on phylogeny construction). From the output of OrthoFinder, we took any protein belonging to the same orthogroup as a known *C. elegans* Argonaute [7] to be a candidate Argonaute. To supplement this set of Argonautes with genes that may have been missed by OrthoFinder (e.g., Argonaute gene families not present in *C. elegans*), we also ran InterProScan v5.52-86.0 [8] using the Pfam database application in order to annotate protein domains in all of the protein sequences for all 51 species. Proteins that were not identified as Argonautes by OrthoFinder, but that contained at least 1 domain characteristic of Argonaute proteins (“Piwi domain”, “PAZ domain”, “Mid domain of argonaute”, “N-terminal domain of argonaute”, “Argonaute linker 1 domain”, or “Argonaute linker 2 domain”) were added to our set of Argonautes, and assigned to one of the 11 Argonaute subfamilies when possible. The only non-Argonaute protein in *C. elegans* that contains any of these domains is DCR-1 (which has a PAZ domain), and so using our OrthoFinder results we excluded all identified orthologs of *C. elegans* DCR-1 from our set of Argonautes. We also added to our set of Argonautes an ortholog of CSR-1 in *C. parvicauda*, which was not annotated in its genome assembly, but was identified in its transcriptome (see “RNA-seq analysis” section). To remove short gene fragments (which may represent genome sequencing or gene annotation errors) from our set of Argonautes, we excluded all Argonautes identified by OrthoFinder whose protein sequences were shorter than half the length of their shortest *C. elegans* ortholog. For Argonautes that were missed by OrthoFinder but that contained Argonaute domains, we excluded all candidate Argonautes whose protein sequences were shorter than half the length of the shortest *C. elegans* Argonaute (PRG-1, with length 824 amino acids).

A gene tree of all 1213 identified Argonautes that passed our length filtering was inferred by first creating a multiple sequence alignment of their protein sequences using the Clustal Omega web server [9]. The resulting sequence alignment was used as input for IQ-TREE v2.1.2 [10] to estimate a maximum likelihood tree, with 1000 ultrafast bootstrap replicates [11] and selecting the best model using ModelFinder [12]. The Argonaute gene tree was then visualized using FigTree v1.4.4 (https://github.com/rambaut/figtree). In addition to the sequence alignment, Clustal Omega’s output also included the percent protein sequence identity between all pairs of the 1213 aligned Argonautes. After setting values of “NA” (cases where the sequence of an Argonaute only aligns to gaps in the other Argonaute) to 0, these sequence identity scores were used to calculate the median percent identity between all pairs of different Argonautes belonging to the same subfamily, for each of the 11 Argonaute subfamilies. A gene arrow diagram of the *C. panamensis* novel Argonaute cluster was created with gggenes v0.5.1 (https://wilkox.org/gggenes/). To search for ALG-5 in nematode species outside of *Caenorhabditis*, we downloaded the set of annotated proteins for 5 species (*Diploscapter coronatus, Diploscapter pachys*, *Haemonchus contortus*, *Oscheius tipulae*, and *Pristionchus pacificus*) from a combination of WormBase release WS282 (*O. tipulae*, *P. pacificus*) and WormBase ParaSite release WBPS15 (*D. coronatus*, *D. pachys*, *H. contortus*) [2, 3]. We searched each of these protein sets with blastp v2.9.0 [13] using the ALG-5 protein sequence from *C. elegans*, but the best BLAST hit for each species was an ortholog of ALG-1 (upon BLASTing these blastp hits back against the set of *C. elegans* protein sequences, the best blastp hit in *C. elegans* was always ALG-1, with much higher sequence similarity than was seen with ALG-5), suggesting these species lack an ALG-5 ortholog. We additionally searched for ALG-5 in the reference transcriptome for *Caenorhabditis krikudae*, using the set of annotated proteins from the Caenorhabditis Genomes Project [4]. Using blastp v2.9.0 [13], we found a reciprocal best BLAST hit for *C. elegans* ALG-5 in *C. krikudae* (a transcript named TRINITY_DN270_c0_g2_i1.p1), suggesting that ALG-5 is likely present in *C. krikudae*.

We used our identified Argonaute genes to count the number of Argonautes from all 11 subfamilies in each of the 51 species, correcting our counts for identified issues of genome assembly and gene annotation as follows. To verify observed cases where a species did not have any Argonautes from a given Argonaute subfamily, we used tblastn v2.9.0 [13] to search the corresponding genome assembly for that species, using the protein sequence of an ortholog of the gene of interest from a closely related species as the query (e.g., PRG-1 from *C. bovis* to search for PRG-1 in *C. plicata*). While doing so, we identified 4 cases where a species did not appear to have a certain type of Argonaute gene that passed our length cutoff, but which had one or more gene fragments resembling the “missing” Argonaute – these cases were VSRA-1 in *C. japonica* (CJAPO.CJA34825a), ALG-3/4 in *C. angaria* (CANGA.Cang_2012_03_13_00748.g13857.t1, CANGA.Cang_2012_03_13_10466.g22668.t1, and CANGA.Cang_2012_03_13_17541.g25720.t1), ALG-3/4 in *C.* sp. 45 (CSP45.TRINITY_DN19321_c0_g1_i2.p1, CSP45.TRINITY_DN30727_c0_g1_i1.p1, and CSP45.TRINITY_DN23301_c0_g1_i1.p1), and ALG-3/4 in *C. waitukubuli* (CWAIT.CSP39.g21541.t1, CWAIT.CSP39.g19211.t1, CWAIT.CSP39.g10631.t1, CWAIT.CSP39.g10909.t1, CWAIT.CSP39.g11814.t1, CWAIT.CSP39.g13006.t1, and CWAIT.CSP39.g2776.t1). For counting purposes we considered these species to have 1 copy of the corresponding Argonaute (e.g., *C. waitukubuli* has no ALG-3/4 orthologs that passed our length cutoff, but we identified 7 gene fragments in this species that were all highly similar to ALG-3/4, so we consider this species to have 1 copy of ALG-3/4), though the fragments were excluded in the gene tree visualization. For *C. latens*, we counted the gene OZG10323.1 as two copies of ERGO-1 and OZG23985.1 as two copies of VSRA-1, as these genes each appeared to be tandem duplicates that were erroneously annotated as fused genes. Because the draft *C. brenneri* reference genome is known to contain extensive allelism masquerading as paralogy [14, 15], causing this species to have an abnormally high observed rate of single-copy gene duplication (27.7% of BUSCO genes were duplicated, Table S2), we aimed to create more accurate Argonaute counts for *C. brenneri*. We classified pairs of Argonautes as allelic variants of a single gene if their protein sequences were at least 95% identical to each other and they had a synonymous substitution rate (K_S_) less than 0.141, the expected amount of synonymous-site divergence between two *C. brenneri* alleles [15]. Jukes-Cantor corrected values of K_S_ were calculated for pairs of *C. brenneri* Argonautes by using MEGA v11 [16] after first creating codon-based alignments of their coding sequences with MUSCLE [17]. Using this approach we found 9 pairs of Argonautes in *C. brenneri* that appeared to be allelic variants of the same gene, and we therefore only counted one Argonaute from each of these 9 pairs (Table S4, Fig. S11). As an example, *C. brenneri* has 2 annotated orthologs of PRG-1 (CBN15497 and CBN22740) that have 100% identical protein sequences, an unusually low level of synonymous-site divergence (K_S_ = 0.0842), and whose directly adjacent genes are also highly similar to each other, and so we count these PRG-1 orthologs as likely allelic variants of a single gene. We repeated this procedure of identifying allelic variants in *C.* sp. 48, the sister species of *C. brenneri*, but only identified a set of 3 genes (CSP48.g8025.t1, CSP48.g8029.t1, and CSP48.g8031.t1) that may be alleles of a single gene (protein sequence identity ranges from 97.52 – 97.81%, K_S_ ranges from 0.112 – 0.124). However, given that these K_S_ values are close to our *C. brenneri*-specific cutoff, and that the expected amount of synonymous-site divergence between two *C.* sp. 48 alleles may be lower than this, we elected to count these 3 genes as 3 separate Argonautes.

Within-species Argonaute sequence variation

Within-species variation data for *C. elegans* and *C. briggsae* Argonautes were downloaded from the Variant Annotation tool on CaeNDR release 20220216 (for *C. elegans*) and release 20240129 (for *C. briggsae*) [18]. For each Argonaute gene in both species, we retrieved all unique variants annotated as being “synonymous” or “missense” (i.e., non-synonymous), or whose annotations suggest they may be pseudogene-causing (all unique variants carrying the annotation “frameshift”, “start_lost”, “stop_gained”, “stop_lost”, “splice_acceptor”, or “splice_donor”). As the reference genome assembly and gene annotations used on CaeNDR for *C. briggsae* are for a different strain (QX1410) compared to the assembly we used to detect Argonautes (AF16), we made use of the orthology calls between the AF16 and QX1410 genome assemblies from [19] (https://github.com/AndersenLab/briggsae_gene_models_MS/blob/main/AFQX_orthology/Orthogroups.tsv). Any QX1410 gene called as being orthologous to a known Argonaute in AF16 was also classified as an Argonaute for the purposes of retrieving variants from CaeNDR.

For each Argonaute in both *C. elegans* and *C. briggsae*, we calculated the number of unique isotypes carrying at least 1 variant of each type (synonymous, non-synonymous, and potentially pseudogene-causing). We additionally calculated per-site Watterson’s θ (the normalized number of variants per site) at both synonymous and non-synonymous sites for each Argonaute. To calculate the total number of synonymous and non-synonymous sites in each Argonaute for this calculation, we used degenotate v1.3 (https://github.com/harvardinformatics/degenotate) to count the number of 0-fold, 2-fold, 3-fold, and 4-fold degenerate sites in the coding sequence of each Argonaute. The number of synonymous sites in a gene is equal to (number of 4-fold sites) + (1/3 * number of 2-fold sites) + (2/3 * number of 3-fold sites). Similarly, the number of non-synonymous sites in a gene is equal to (number of 0-fold sites) + (2/3 * number of 2-fold sites) + (1/3 * number of 3-fold sites). Watterson’s θ was calculated assuming a sample of 550 diploid individuals for *C. elegans* and 641 diploid individuals for *C. briggsae*, based on the number of unique isotypes included in each CaeNDR release.

3D structural analysis

Files of computationally-predicted 3D structures for all 20 *C. elegans* Argonaute proteins, as well as *C. elegans* DCR-1 (for use as an outgroup) were downloaded from the AlphaFold Protein Structure Database [20] (Table S5). For each of the 7 *C. panamensis* Argonaute proteins that potentially belong to a novel subfamily of ALGs (as well as CSR-1 from *C. panamensis* as a control), we generated AlphaFold predictions of their 3D structures using ColabFold v1.5.5, by running the AlphaFold2_mmseqs2 Google Colab notebook with default settings [21, 22]. Prior to structure prediction, we manually trimmed the protein sequences of two of these putatively novel Argonautes (CSP28.g8935.t1 and CSP28.g16021.t1) to remove sequences from adjacent genes that were erroneously included in these gene models. To determine which Argonaute proteins had the most similar structures to our “novel” *C. panamensis* Argonautes, we used the DALI web server [23] to perform structural alignments between all pairs of our 29 protein structures (21 from *C. elegans* and 8 from *C. panamensis*), and used the reported Z-scores for each alignment as our measure of structural similarity. We additionally used these DALI structural alignments to assess whether each protein structure contained an N, PAZ, PIWI, and MID domain. Given that DALI provided Pfam annotations for all of these domains in ALG-1 from *C. elegans*, we viewed the pairwise structural alignments of each of our protein structures against *C. elegans* ALG-1, and determined whether the N, PAZ, PIWI, and MID domains of ALG-1 aligned to corresponding structures in the other protein (as opposed to these domains aligning to a gap or a disordered region). Heatmaps of protein sequence identity and structural similarity were created using ComplexHeatmap v2.12.1 [24], clustering rows and columns with hierarchical clustering based on sequence identity. 3D protein structures were visualized using the Mol* Viewer [25] on RCSB Protein Data Bank [26].

RNA-seq analysis

Raw gene expression data from SRA was retrieved for 14 different *Caenorhabditis* species: *C. afra, C.* *agridulce*, *C. castelli*, *C. inopinata, C. macrosperma, C. monodelphis*, *C. nouraguensis, C. parvicauda*, *C. plicata*, *C. quiockensis*, *C. remanei*, *C. tribulationis, C. uteleia,* and *C. waitukubuli* (Table S6), using seqtk v1.3 (https://github.com/lh3/seqtk) to split paired-end FASTQ files into separate files for the first and second reads in each pair when necessary. The quality of each FASTQ file was assessed with FastQC v0.11.7 (https://www.bioinformatics.babraham.ac.uk/projects/fastqc/), and sequencing adapters were trimmed from reads using TrimGalore v0.6.6 (https://github.com/FelixKrueger/TrimGalore) with a stringency value of 3 and using paired-end mode for all paired-end sequencing files. Trimmed RNA-seq reads were aligned to their respective reference genomes using STAR v2.5.3a [27]. Read alignments were filtered with the “view” function from SAMtools v1.8 [28] to exclude reads that did not align uniquely (-q option set to 255) or (for paired-end files) that did not align as a proper pair (-f option set to 0x2). SAMtools was additionally used to remove duplicate reads from paired-end files using the “sort”, “fixmate”, and “markdup” functions. Filtered RNA-seq read alignments at the CSR-1 locus for each species were visualized with the IGV web server [29]. Additionally, we used Trinity v2.13.2 [30] to make a de novo transcriptome assembly for each of the 14 species, using the trimmed FASTQ files as input. To verify apparent gene losses in species with available RNA-seq data (e.g., CSR-1 in *C. parvicauda*, PRG-1 in *C. plicata*), we used tblastn v2.9.0 [13] to search our transcriptome assembly for that species, using the protein sequence of an ortholog of the gene of interest from a closely related species as the query (e.g., PRG-1 from *C. bovis* to search for PRG-1 in *C. plicata*).

Worm culture and DNA isolation

Molecular biology confirmation was conducted in the following species: *C. drosophilae* (DF5112), *C. macrosperma* (JU1857), *C. panamensis* (QG702), *C. plicata* (SB355), and *C.* sp. 30 (DF5152). *C. elegans* (N2) and *C. virilis* (JU1528) were used as controls for species inside and outside the Elegans supergroup, respectively. All strains were cultured on 10 cm NGM-agar plates seeded with OP50 *Escherichia coli* and maintained at room temperature (~24ºC), except *C. elegans* which was maintained at 20ºC [31, 32]. Prior to DNA extractions, strains were cleaned and age synchronized through hypochlorite treatment [32]. In our hands, *C. drosophilae* did not lay sufficient eggs to withstand hypochlorite treatment. Therefore, adult populations were cleaned by soaking adult worms in M9 buffer with 100ug/mL carbenicillin and 400ug/mL kanamycin for 3 hours. This antibiotic treatment was done for two successive generations prior to DNA isolation. A complete list of strain information can be found in Table S7.

High quality genomic DNA was purified using the Monarch Genomic DNA Purification Kit (NEB). Briefly, at least 1,000 adult worms were washed into 15mL falcon tubes with M9 buffer and then rinsed with clean M9 buffer three additional times. Worms were then pelleted and transferred to 1.5mL microfuge tubes and washed with an additional 1mL of buffer. After the final wash the supernatant was removed, leaving a pelleted worm volume of approximately 200uL. Following the manufacturer recommendations for animal tissues, 200uL of tissue lysis buffer and 10uL of Proteinase K were added to each sample and incubated in a shaking incubator at 56ºC and 220rpm for 3 hours. The samples were then pelleted and the supernatant transferred to a fresh microfuge tube for RNase A treatment. The manufacturer instructions were followed for the DNA binding and elution steps. The eluted DNA was further cleaned using the DNA Clean and Concentrator-5 kit (Zymo). DNA concentration and purity was assessed using Nanodrop (Table S7).

Genomic DNA for *C. virilis* was purified by isolating 10 adult worms in 6uL of 20mg/uL Proteinase K in elution buffer. A total of eight samples (n = 80 worms) were collected. Samples were freeze-cracked at -80ºC for 30 minutes two times. They were then incubated at 58ºC for 60 minutes followed by 10 minutes at 95ºC to inactivate the Proteinase K.

Molecular biology

PCR products were generated using the Q5 High-Fidelity Master Mix (NEB) and 100ng of purified DNA in accordance with manufacturer instructions. We verified that the DNA isolation process was successful in all species by amplifying a 535bp product in the 18S ribosomal DNA region (Table S8). Additionally, we tested the amplification of the conserved gene *act-1* using *C. elegans* specific primers in *C. elegans*, *C. drosophilae*, *C. plicata*, and *C.* sp. 30. Despite having 2-4 mismatches in the primer sequences, all non-*elegans* species had correct amplification, which indicated that primers designed in conserved regions have the potential to amplify across divergent species. A complete list of primers used in this study can be found in Table S8.

To verify the putative duplicate copy of *csr-1* in *C. macrosperma*, we attempted to amplify through the entire gene, starting 220bp upstream of the start codon through 154bp downstream of the stop codon. We additionally attempted to amplify exons 1-3 and exons 12-17 (i.e., the PIWI domain). PCR products of approximately the correct size were gel extracted and purified using the Zymoclean Gel DNA Recovery Kit (Zymo) for verification by Sanger sequencing.

To verify the putative duplicate copy of *prg-1* in *C.* sp. 30, we amplified through the entire gene, starting 219bp upstream of the start codon through 172bp downstream of the stop codon. Additionally, we amplified products starting upstream of the start codon through exon 5 and through exon 10. PCR products for all primer combinations were gel extracted and purified using the Zymoclean Gel DNA Recovery Kit (Zymo) for verification by Sanger sequencing.

To examine the potential loss of *prg-1* in *C. drosophilae* and *C. plicata*, we first attempted to use *C. elegans* specific primers to amplify exons 3-6. As positive controls, DNA from species where *prg-1* was successfully annotated (i.e., *C. elegans*, *C. macrosperma*, *C. panamensis*, and *C.* sp. 30) were included. Only *C. elegans* DNA amplified. We then redesigned primers using a reduced alignment of seven species for which we had DNA along with *C. bovis*. From this alignment, we designed degenerate primers with no more than four ambiguous base pairs in each 20nt forward and reverse primer that amplified exons 4-8. PCRs using the degenerate primers were run at three temperatures: the lowest potential Tm, the highest potential Tm, and an intermediate Tm. All of the positive controls (*C. elegans*, *C. macrosperma*, *C. panamensis*, *C.* sp. 30, and *C. virilis*) amplified correctly for at least one temperature.

To verify the divergent *C. panamensis* ALG duplicate CSP28.g8935.t1, we first attempted to amplify a product starting 117bp upstream of the start codon through 106bp downstream of the stop codon. This reaction produced product. We then attempted to amplify the coding sequence only (exons 1-15). PCR products of approximately the correct size were gel extracted and purified using Zymoclean Gel DNA Recovery Kit (Zymo) for verification by Sanger sequencing. The remaining 6 divergent *C. panamensis* ALG duplicates (which formed a cluster of tandem duplicates) were analyzed using three approaches. First, we attempted to amplify between genes by anchoring the forward primer in the final exon of the upstream gene and the reverse primer in the first or second exon of the downstream gene. We additionally attempted to amplify ~1kb upstream and downstream of each gene independently. Finally, we attempted to amplify the coding sequence of each gene for verification by Sanger sequencing.

Gene family evolution

The species tree for *Caenorhabditis* used in this study was taken from Fusca et al. (in preparation). Briefly, gene trees of BUSCO single-copy genes were made with IQ-TREE v2.1.0 [10] and used to create a species tree with ASTRAL v5.6.3 [33]. Branch lengths on this species tree were then converted to units of absolute time using BEAST v2.7.5 [34], calibrating the tree using strict molecular clock estimates of divergence time based on 1:1 orthologs of the 10 most recently-diverged species pairs.

Significant gene family expansions and contractions across *Caenorhabditis* were detected with CAFE v5.1 [35], using our dated species tree. The number of genes in each orthogroup reported by OrthoFinder were used as the gene family size for CAFE (aside from the 11 Argonaute orthogroups, for which we instead used our manually-corrected gene counts). As CAFE only tests gene families that are inferred to be present at the root of the species tree, we excluded *C. monodelphis* and *C. parvicauda* in order to test all 11 Argonaute families, since these early-diverging species lack copies of ALG-5 and ERGO-1 respectively. CAFE was run using 4 gamma rate categories for the gene birth/death rate, and we excluded the 50 gene families that showed the largest difference in gene family size between species (corresponding to gene families where one species had over 40 more gene copies than another species), in order for CAFE to initialize reasonable values for the gene birth/death rate. We additionally reran CAFE while also excluding the 5 species with the lowest-quality gene annotations according to BUSCO (either fewer than 85% of BUSCO genes complete and single-copy, or over 5% of BUSCO genes complete but duplicated) – these 5 species are *C. angaria*, *C. brenneri*, *C. japonica*, *C.* sp. 49, and *C. waitukubuli*. CAFE detected significant gene families using a P-value threshold of 0.05, and we additionally performed multiple-testing correction on these P-values by calculating False Discovery Rates in R v4.2.1 [36], taking gene families with an FDR ≤ 0.05 to be significant.

To model genomic repeat content as a function of Argonaute gene copy number, we ran RepeatModeler v2.0.3 [37] to identify repeat families in each of our 50 reference genome assemblies, using the -LTRStruct option to also detect LTR transposons. The coordinates of repeats in each genome were then identified and masked using RepeatMasker v4.1.2 (https://www.repeatmasker.org/) with the -s and -nolow options. We defined repeat content for a species as the proportion of that species’ genome that was masked by RepeatMasker (Table S1). All phylogenetic generalized least squares (PGLS) regressions in this study were performed in R with caper v1.0.2 (https://cran.r-project.org/web/packages/caper/index.html), using our dated species tree as the phylogeny and excluding *C.* sp. 45 (as we only have a transcriptome assembly for this species). PGLS regressions were performed using the following 6 linear models:

1. Total Argonaute number ~ genome size
2. Total Argonaute number ~ genome assembly N50
3. Total protein-coding gene number ~ genome size
4. Total Argonaute number ~ total protein-coding gene number
5. Repeat content ~ genome size + total number of non-Argonaute protein-coding genes + total number of Argonaute genes
6. Repeat content ~ genome size + total number of non-Argonaute protein-coding genes + number of RDE-1 orthologs + number of ALG-1/2 orthologs + number of ALG-3/4 orthologs + number of ALG-5 orthologs + number of CSR-1 orthologs + number of ERGO-1 orthologs + number of PRG-1 orthologs + number of VSRA-1 orthologs + number of WAGO-1/3/4/5 orthologs + number of WAGO-6/7/8 orthologs + number of WAGO-9/10/12 orthologs

Both regressions involving repeat content were repeated after excluding the 5 species with the lowest-quality gene annotations (*C. angaria*, *C. brenneri*, *C. japonica*, *C.* sp. 49, and *C. waitukubuli*). Regressions involving repeat content were also repeated for three specific types of repeats (retrotransposons, DNA transposons, and rolling-circle transposons), taking the dependent variable to be the proportion of the genome annotated by RepeatMasker as belonging to that repeat type.

piRNA pathway analysis

The presence or absence of 7 genes involved in piRNA transcription and processing (*prde-1*, *snpc-4*, *tofu-4*, *tofu-5*, *pid-1*, *henn-1*, and *parn-1*) [38] in all 51 species was determined based on whether OrthoFinder was able to identify any orthologs of these genes in each species. Apparent gene absences were then verified by using tblastn v2.9.0 [13] to search for the gene in the corresponding genome sequence (as well as our assembled transcriptome, when possible) as described previously for Argonautes. Doing so allowed us to identify several piRNA pathway genes that were missed by OrthoFinder, but still present in certain species’ genomes (e.g., *parn-1* in *C. bovis*).

Ruby motifs were detected in the noncoding portion of all 50 available reference genomes (i.e., every species except for *C.* sp. 45) by first using the “maskfasta” function of BEDTools v2.30.0 [39] to mask all sequences belonging to exons for each genome. The “locate” function of SeqKit v2.3.0 [40] was then used to find and count all instances of the sequence “CNGTTTCA” (where “N” can be any nucleotide) and its reverse complement in each masked genome sequence, based on the known Ruby motif sequence in *Caenorhabditis* [41]. To compare our counts of Ruby motifs to what would be expected under random chance, we used the “compseq” function from EMBOSS v6.6.0 [42] to calculate the frequencies of all 16 dinucleotides (e.g., “AC”, “GT”) in each masked genome sequence. The number of expected Ruby motifs in each species was then calculated as the probability of a random 8-mer in a species being “CNGTTTCA” or its reverse complement given the dinucleotide content of its noncoding genome, multiplied by the number of 8-mers in a DNA sequence as long as that species’ noncoding genome (i.e., noncoding genome size – 7). We verified that this approach provides an accurate expectation by simulating 1000 random DNA sequences of equal length and dinucleotide content in R using the “rDNAbin” function in ape v5.6.2 [43], and found that the mean number of Ruby motifs observed in the random genomes matched our calculated expectation. For species with unusually low levels of the Ruby motif (*C. drosophilae*, *C.* sp. 2, *C. monodelphis*, *C. bovis*, *C. plicata*, *C. parvicauda*, *C.* sp. 30, and *C. japonica*), we repeated this counting procedure using the 6-mer “GTTTCA”, which is the core of the Ruby motif and is more widely conserved across nematodes outside of *Caenorhabditis* [41]. However, we observed similarly low levels of the Ruby motif in these species even when using this shorter sequence, and so we only present results for the full CNGTTTCA motif. Genomic locations of known piRNA clusters in *C. elegans* and *C. briggsae* were taken from [44] and [45], respectively.

CSR-1 sequence evolution

DNA coding sequences were retrieved for all identified CSR-1 orthologs, aside from very highly-diverged CSR-1 duplicates in *C. parvicauda* (CSP21.g381.t1) and *C. macrosperma* (CMACR.g1069.t1) which may be genome assembly artifacts, resulting in a total of 58 CSR-1 sequences. For 5 species (*C. drosophilae*, *C.* sp. 2, *C.* sp. 51, *C. sulstoni*, and *C. tribulationis*), the annotated CSR-1 orthologs appeared to be fused to the adjacent gene FRPR-10. Given that RNA-seq data in *C. tribulationis* showed no evidence of CSR-1 and FRPR-10 being expressed as a single transcript, these fusions are likely annotation errors, and for these 5 species the portions of the CSR-1 coding sequence corresponding to the FRPR-10 fusion were manually removed. In addition to this set of 58 sequences, we also created a subset that contained only a single CSR-1 ortholog for each of the 51 species, by keeping only a single CSR-1 ortholog from the species that had multiple orthologs (CPLIC.g9864.t1 for *C. plicata*, CBN29619 for *C. brenneri*, CSP48.g7722.t1 for *C.* sp. 48, CSP54.g17215.t1 for *C.* sp. 54, and CDOUG.g9300.t1 for *C. doughertyi*). Both sets of CSR-1 sequences were separately aligned using PRANK v.170427 [46] in codon-alignment mode. Codons that could not be aligned reliably were filtered out from both alignments with GUIDANCE v2.0.2 [47], using 50 bootstrap runs and a cutoff score of 0.93. For the alignment containing all 58 CSR-1 sequences, a maximum-likelihood gene tree was constructed by running IQ-TREE v2.1.2 [10] with 1000 ultrafast bootstrap replicates [11] and selecting the best model using ModelFinder [12].

Branch-site tests of positive selection were performed on both the full 58-sequence alignment and the reduced 51-sequence alignment using both aBSREL v2.5 [48] and BUSTED v4.1 [49], hosted on the Datamonkey web server [50]. For the corresponding phylogeny needed for these analyses, we used our CSR-1 gene tree for the full 58-sequence alignment (rooting the tree using the *C. monodelphis* ortholog as the outgroup), and our *Caenorhabditis* species tree for the 51-sequence alignment. aBSREL was run on all branches of the corresponding phylogeny, using aBSREL’s built-in corrections for multiple testing. BUSTED was run either only on the branch leading to the Elegans group, or only on the branch leading to the Elegans supergroup, allowing for site-to-site synonymous rate variation in the branch-site model, but not multinucleotide substitutions. The aBSREL tests using the 51-sequence alignment were repeated after trimming off alignment columns in a highly gap-filled region containing what remained of the CSR-1a exon after GUIDANCE filtering, such that the trimmed alignment begins 39 bases into exon 2 of CSR-1 in *C. elegans*.

The gene structure of CSR-1 across the genus was visualized using TBtools v1.119 [51], again taking only a single representative CSR-1 ortholog for species with multiple orthologs, exactly as was done for our branch-site tests. The locations of arginine-glycine (RG) repeats in each CSR-1 sequence were detected using the “locate” function of SeqKit v2.3.0 [40]. We attempted to detect the CSR-1a exon in all of the species within the Elegans supergroup that did not have it annotated (*C. japonica*, *C.* sp. 25, *C. afra*, *C. sulstoni*, *C. panamensis*, and *C. becei*), as well as a subset of species outside of the Elegans supergroup (*C. uteleia*, *C. castelli*, *C. portoensis*, *C. dolens*, *C. monodelphis*, and *C. astrocarya*). To do so, we retrieved the 5kb of DNA upstream of the CSR-1 start codon using the “getfasta” function of BEDTools v2.30.0 [39], translated this DNA in all 3 reading frames using the Expasy web server [52], and searched for stretches of amino acids beginning with an “M” and containing multiple RG repeats without any intervening stop codons. For this gene structure visualization, we manually corrected the CSR-1 gene models for these Elegans supergroup species to contain the identified putative CSR-1a exon: for *C. japonica* we reclassified the annotated 5′ UTR as a coding exon (starting at the start codon located 12 bases into the exon and removing 1 base from the end to avoid introducing a frameshift); for *C.* sp. 25, *C. sulstoni*, *C. panamensis*, and *C. becei* we took the upstream RG-rich open reading frame to be the CSR-1a exon (beginning with the start codon and ending with the final RG repeat; for *C. sulstoni* we extended this exon to also include the currently-annotated short first exon); and for *C. afra* we took the true start codon to be the ATG codon in the second exon (as the first exon is unusually short, and the second exon contains many RG repeats when translated in this new reading frame) and removed 1 base from the end of the second exon to avoid introducing a frameshift. We found several species (*C. macrosperma*, *C. waitukubuli*, *C. nouraguensis*, and one *C. brenneri* ortholog) where the putative CSR-1a exon annotation appears split into two adjacent exons, though gene expression data indicates these are likely annotation errors in the *C. macrosperma* and *C. nouraguensis* genomes. We used tblastn v2.9.0 and blastp v2.9.0 [13] to search for the first 163 amino acids of *C. elegans* CSR-1 (corresponding to the CSR-1a exon) in the genomes (or transcriptome, for *C.* sp. 45) and annotated proteomes of 5 species outside of the Elegans supergroup (*C. uteleia*, *C.* sp. 45, *C. astrocarya*, *C. castelli*, and *C.* sp. 27).

Supplementary References

1. Sayers EW, Bolton EE, Brister JR, Canese K, Chan J, Comeau DC, et al. Database resources of the national center for biotechnology information. Nucleic Acids Res. 2022;50(D1):D20-D6. pmid:34850941.
2. Howe KL, Bolt BJ, Shafie M, Kersey P, Berriman M. WormBase ParaSite - a comprehensive resource for helminth genomics. Mol Biochem Parasitol. 2017;215:2-10. pmid:27899279.
3. Davis P, Zarowiecki M, Arnaboldi V, Becerra A, Cain S, Chan J, et al. WormBase in 2022-data, processes, and tools for analyzing *Caenorhabditis elegans*. Genetics. 2022;220(4). pmid: 35134929.
4. Stevens L. Data associated with the Caenorhabditis Genomes Project (caenorhabditis.org); 2024 [cited 2024 July 4]. Database: Zenodo [Internet]. Available from: https://doi.org/10.5281/zenodo.12633738
5. Manni M, Berkeley MR, Seppey M, Simão FA, Zdobnov EM. BUSCO Update: Novel and Streamlined Workflows along with Broader and Deeper Phylogenetic Coverage for Scoring of Eukaryotic, Prokaryotic, and Viral Genomes. Mol Biol Evol. 2021;38(10):4647-54. pmid:34320186.
6. Emms DM, Kelly S. OrthoFinder: phylogenetic orthology inference for comparative genomics. Genome Biol. 2019;20(1):238. pmid:31727128.
7. Youngman EM, Claycomb JM. From early lessons to new frontiers: the worm as a treasure trove of small RNA biology. Front Genet. 2014;5:416. pmid:25505902.
8. Jones P, Binns D, Chang HY, Fraser M, Li W, McAnulla C, et al. InterProScan 5: genome-scale protein function classification. Bioinformatics. 2014;30(9):1236-40. pmid:24451626.
9. Madeira F, Pearce M, Tivey ARN, Basutkar P, Lee J, Edbali O, et al. Search and sequence analysis tools services from EMBL-EBI in 2022. Nucleic Acids Res. 2022;50(W1):W276-W9. pmid:35412617.
10. Minh BQ, Schmidt HA, Chernomor O, Schrempf D, Woodhams MD, von Haeseler A, et al. IQ-TREE 2: New Models and Efficient Methods for Phylogenetic Inference in the Genomic Era. Mol Biol Evol. 2020;37(5):1530-4. pmid:32011700.
11. Hoang D, Chernomor O, von Haeseler A, Minh B, Vinh L. UFBoot2: Improving the Ultrafast Bootstrap Approximation. Molecular Biology and Evolution. 2018;35(2):518-22. pmid:29077904.
12. Kalyaanamoorthy S, Minh BQ, Wong TKF, von Haeseler A, Jermiin LS. ModelFinder: fast model selection for accurate phylogenetic estimates. Nat Methods. 2017;14(6):587-9. pmid:28481363.
13. Camacho C, Coulouris G, Avagyan V, Ma N, Papadopoulos J, Bealer K, et al. BLAST+: architecture and applications. BMC Bioinformatics. 2009;10:421. pmid:20003500.
14. Barrière A, Yang SP, Pekarek E, Thomas CG, Haag ES, Ruvinsky I. Detecting heterozygosity in shotgun genome assemblies: Lessons from obligately outcrossing nematodes. Genome Res. 2009;19(3):470-80. pmid:19204328.
15. Dey A, Chan CK, Thomas CG, Cutter AD. Molecular hyperdiversity defines populations of the nematode *Caenorhabditis brenneri*. Proc Natl Acad Sci U S A. 2013;110(27):11056-60. pmid:23776215.
16. Tamura K, Stecher G, Kumar S. MEGA11: Molecular Evolutionary Genetics Analysis Version 11. Mol Biol Evol. 2021;38(7):3022-7. pmid:33892491.
17. Edgar RC. MUSCLE: multiple sequence alignment with high accuracy and high throughput. Nucleic Acids Res. 2004;32(5):1792-7. pmid:15034147.
18. Crombie TA, McKeown R, Moya ND, Evans KS, Widmayer SJ, LaGrassa V, et al. CaeNDR, the *Caenorhabditis* Natural Diversity Resource. Nucleic Acids Res. 2024;52(D1):D850-D8. pmid:37855690.
19. Moya ND, Stevens L, Miller IR, Sokol CE, Galindo JL, Bardas AD, et al. Novel and improved *Caenorhabditis briggsae* gene models generated by community curation. BMC Genomics. 2023;24(1):486. pmid:37626289.
20. Varadi M, Anyango S, Deshpande M, Nair S, Natassia C, Yordanova G, et al. AlphaFold Protein Structure Database: massively expanding the structural coverage of protein-sequence space with high-accuracy models. Nucleic Acids Res. 2022;50(D1):D439-D44. pmid:34791371.
21. Jumper J, Evans R, Pritzel A, Green T, Figurnov M, Ronneberger O, et al. Highly accurate protein structure prediction with AlphaFold. Nature. 2021;596(7873):583-9. pmid:34265844.
22. Mirdita M, Schütze K, Moriwaki Y, Heo L, Ovchinnikov S, Steinegger M. ColabFold: making protein folding accessible to all. Nat Methods. 2022;19(6):679-82. pmid:35637307.
23. Holm L. Dali server: structural unification of protein families. Nucleic Acids Res. 2022;50(W1):W210-W5. pmid:35610055.
24. Gu Z, Eils R, Schlesner M. Complex heatmaps reveal patterns and correlations in multidimensional genomic data. Bioinformatics. 2016;32(18):2847-9. pmid:27207943.
25. Sehnal D, Bittrich S, Deshpande M, Svobodová R, Berka K, Bazgier V, et al. Mol* Viewer: modern web app for 3D visualization and analysis of large biomolecular structures. Nucleic Acids Res. 2021;49(W1):W431-W7. pmid:33956157.
26. Burley SK, Bhikadiya C, Bi C, Bittrich S, Chao H, Chen L, et al. RCSB Protein Data Bank (RCSB.org): delivery of experimentally-determined PDB structures alongside one million computed structure models of proteins from artificial intelligence/machine learning. Nucleic Acids Res. 2023;51(D1):D488-D508. pmid:36420884.
27. Dobin A, Davis CA, Schlesinger F, Drenkow J, Zaleski C, Jha S, et al. STAR: ultrafast universal RNA-seq aligner. Bioinformatics. 2013;29(1):15-21. pmid:23104886.
28. Li H, Handsaker B, Wysoker A, Fennell T, Ruan J, Homer N, et al. The Sequence Alignment/Map format and SAMtools. Bioinformatics. 2009;25(16):2078-9. pmid:19505943.
29. Robinson JT, Thorvaldsdóttir H, Winckler W, Guttman M, Lander ES, Getz G, et al. Integrative genomics viewer. Nat Biotechnol. 2011;29(1):24-6. pmid:21221095.
30. Grabherr MG, Haas BJ, Yassour M, Levin JZ, Thompson DA, Amit I, et al. Full-length transcriptome assembly from RNA-Seq data without a reference genome. Nat Biotechnol. 2011;29(7):644-52. pmid:21572440.
31. Brenner S. The genetics of *Caenorhabditis elegans*. Genetics. 1974;77(1):71-94. pmid:4366476.
32. Kenyon C. The nematode *Caenorhabditis elegans*. Science. 1988;240(4858):1448-53. pmid:3287621.
33. Zhang C, Rabiee M, Sayyari E, Mirarab S. ASTRAL-III: polynomial time species tree reconstruction from partially resolved gene trees. BMC Bioinformatics. 2018;19(Suppl 6):153. pmid:29745866.
34. Bouckaert R, Vaughan TG, Barido-Sottani J, Duchêne S, Fourment M, Gavryushkina A, et al. BEAST 2.5: An advanced software platform for Bayesian evolutionary analysis. PLoS Comput Biol. 2019;15(4):e1006650. pmid:30958812.
35. Mendes FK, Vanderpool D, Fulton B, Hahn MW. CAFE 5 models variation in evolutionary rates among gene families. Bioinformatics. 2021;36(22-23):5516-8. pmid:33325502.
36. R Core Team. R: A language and environment for statistical computing. R Foundation for Statistical Computing, Vienna, Austria. 2022. URL https://www.R-project.org/.
37. Flynn JM, Hubley R, Goubert C, Rosen J, Clark AG, Feschotte C, et al. RepeatModeler2 for automated genomic discovery of transposable element families. Proc Natl Acad Sci U S A. 2020;117(17):9451-7. pmid:32300014.
38. Almeida MV, Andrade-Navarro MA, Ketting RF. Function and Evolution of Nematode RNAi Pathways. Noncoding RNA. 2019;5(1). pmid:30650636.
39. Quinlan AR, Hall IM. BEDTools: a flexible suite of utilities for comparing genomic features. Bioinformatics. 2010;26(6):841-2. pmid:20110278.
40. Shen W, Le S, Li Y, Hu F. SeqKit: A Cross-Platform and Ultrafast Toolkit for FASTA/Q File Manipulation. PLoS One. 2016;11(10):e0163962. pmid:27706213.
41. Beltran T, Barroso C, Birkle TY, Stevens L, Schwartz HT, Sternberg PW, et al. Comparative Epigenomics Reveals that RNA Polymerase II Pausing and Chromatin Domain Organization Control Nematode piRNA Biogenesis. Dev Cell. 2019;48(6):793-810.e6. pmid:30713076.
42. Rice P, Longden I, Bleasby A. EMBOSS: the European Molecular Biology Open Software Suite. Trends Genet. 2000;16(6):276-7. pmid:10827456.
43. Paradis E, Schliep K. ape 5.0: an environment for modern phylogenetics and evolutionary analyses in R. Bioinformatics. 2019;35(3):526-8. pmid:30016406.
44. Ruby JG, Jan C, Player C, Axtell MJ, Lee W, Nusbaum C, et al. Large-scale sequencing reveals 21U-RNAs and additional microRNAs and endogenous siRNAs in *C. elegans*. Cell. 2006;127(6):1193-207. pmid:17174894.
45. Shi Z, Montgomery TA, Qi Y, Ruvkun G. High-throughput sequencing reveals extraordinary fluidity of miRNA, piRNA, and siRNA pathways in nematodes. Genome Res. 2013;23(3):497-508. pmid:23363624.
46. Löytynoja A, Goldman N. Phylogeny-aware gap placement prevents errors in sequence alignment and evolutionary analysis. Science. 2008;320(5883):1632-5. pmid:18566285.
47. Sela I, Ashkenazy H, Katoh K, Pupko T. GUIDANCE2: accurate detection of unreliable alignment regions accounting for the uncertainty of multiple parameters. Nucleic Acids Res. 2015;43(W1):W7-14. pmid:25883146.
48. Smith MD, Wertheim JO, Weaver S, Murrell B, Scheffler K, Kosakovsky Pond SL. Less is more: an adaptive branch-site random effects model for efficient detection of episodic diversifying selection. Mol Biol Evol. 2015;32(5):1342-53. pmid:25697341.
49. Lucaci AG, Zehr JD, Enard D, Thornton JW, Kosakovsky Pond SL. Evolutionary Shortcuts via Multinucleotide Substitutions and Their Impact on Natural Selection Analyses. Mol Biol Evol. 2023;40(7). pmid:37395787.
50. Weaver S, Shank S, Spielman S, Li M, Muse S, Pond S. Datamonkey 2.0: A Modern Web Application for Characterizing Selective and Other Evolutionary Processes. Molecular Biology and Evolution. 2018;35(3):773-7. pmid:29301006.
51. Chen C, Chen H, Zhang Y, Thomas HR, Frank MH, He Y, et al. TBtools: An Integrative Toolkit Developed for Interactive Analyses of Big Biological Data. Mol Plant. 2020;13(8):1194-202. pmid:32585190.
52. Duvaud S, Gabella C, Lisacek F, Stockinger H, Ioannidis V, Durinx C. Expasy, the Swiss Bioinformatics Resource Portal, as designed by its users. Nucleic Acids Res. 2021;49(W1):W216-W27. pmid:33849055.
